# Differential encoding in prefrontal cortex projection neuron classes across cognitive tasks

**DOI:** 10.1101/2020.03.14.991018

**Authors:** Jan H. Lui, Nghia D. Nguyen, Sophie M. Grutzner, Spyros Darmanis, Diogo Peixoto, Mark J. Wagner, William E. Allen, Justus M. Kebschull, Ethan B. Richman, Jing Ren, William T. Newsome, Stephen R. Quake, Liqun Luo

## Abstract

Single-cell transcriptomics has been widely applied to classify neurons in the mammalian brain, while systems neuroscience has historically analyzed the encoding properties of cortical neurons without considering cell types. Here we examine how specific transcriptomic types of mouse prefrontal cortex (PFC) projection neurons relate to axonal projections and encoding properties across multiple cognitive tasks. We found that most types projected to multiple targets, and most targets received projections from multiple types, except PFC→PAG (periaqueductal gray). By comparing Ca^2+^-activity of the molecularly homogeneous PFC→PAG type against two heterogeneous classes in several two-alternative choice tasks in freely-moving mice, we found that all task-related signals assayed were qualitatively present in all examined classes. However, PAG-projecting neurons most potently encoded choice in cued tasks, whereas contralateral PFC-projecting neurons most potently encoded reward context in an uncued task. Thus, task signals are organized redundantly, but with clear quantitative biases across cells of specific molecular-anatomical characteristics.

## INTRODUCTION

Achieving a ‘ground truth’ understanding of neuronal types is an important step in dissecting the function of complex neuronal circuits (Jorgenson et al., 2015; Luo et al., 2018). Molecular neuroscience has seen a recent explosion in neuronal cell type classification using single-cell RNA sequencing technologies (reviewed in Zeng and Sanes, 2017). Because transcriptomic data reflects cellular function, is high dimensional, and can be quantitatively compared across brain regions (Tasic et al., 2018) and species (Tosches et al., 2018; Hodge et al., 2019; Kebschull et al., bioRxiv 10.1101/2020.06.25.170118), it is often considered as a foundation to all other properties. However, it is challenging to reconcile cell type definitions from transcriptomic data with those determined by other properties such as developmental history, connectivity patterns, electrophysiological properties, and the encoding of signals related to behavior. Most transcriptomic studies have not investigated the encoding of behaviorally relevant signals in discovered neuronal cell types *in vivo.* Furthermore, what constitutes a neuronal type in many regions of the mammalian brain is a topic of intense debate (Yuste et al., 2020). Explicitly testing whether a given transcriptomic classification possesses any characteristic anatomical and physiological properties can help determine whether the classification is functionally relevant.

This issue is beginning to be addressed in a variety of neural systems. Applying single-cell transcriptomic analysis to systems with a pre-existing ground truth of cell types and function (e.g., mouse retina: Shekhar et al., 2016; fly olfactory system: Li et al., 2017) has resulted in relatively faithful mapping between molecular and functional types. Interrogation of two subcortically-projecting transcriptomic types in mouse anterolateral motor cortex (ALM) during a motor planning task revealed differences in the encoding of preparatory activity (Economo et al., 2018). Transcriptomic profiling of neuronal diversity in the mouse hypothalamus revealed high levels of transcriptomic diversity (Moffitt et al., 2018; Kim et al., 2019) but little one-to-one matching between transcriptomic types, behavior-specific activation, and connectivity (Kim et al., 2019). Transcriptomic analysis of the mouse striatum revealed that continuous variation in gene expression is overlaid on discrete cell types over space, with both continuous and discrete variation contributing to circuit function (Gokce et al., 2016; Stanley et al., 2020). Thus, the level of correspondence between molecular and functional properties can differ substantially across neuron types and brain regions.

Neurons of the mammalian prefrontal cortex (PFC) serve at a critical transition between sensation and action, bias diverse sensory signals towards appropriate downstream targets, and underlie cognitive processes such as reward-guided decision-making and behavioral flexibility (Fuster, 2008; Miller and Cohen, 2001; Rushworth et al., 2011; Euston et al., 2012; Zingg et al., 2014). While traditional systems neuroscience techniques have shown how complex task-related signals can be encoded at the single-neuron as well as population levels (e.g., Asaad et al., 1998; Mante et al., 2013), these studies are typically blind to cell type and their projection patterns. Furthermore, cognitive functions, by their nature, operate among a wide range of task demands, and the measured complexity of how PFC encodes task signals depends heavily on the behavioral assays used. It is therefore important to achieve an integrated picture of how specific transcriptomic cell types relate to their projection patterns, and together underlie a well-defined repertoire of task signals.

To this end, recent studies in mice have suggested that nucleus accumbens-projecting medial PFC (mPFC→NAc) neurons have different roles in the conjunctive encoding of social and spatial targets (Murugan et al., 2017), the restraint of reward seeking (Kim et al., 2017), and the representation of reward predicting cues (Otis et al., 2017). mPFC→PAG neurons were reported to be a key node that dopamine acts on to modulate the signal-to-noise ratio in encoding of aversive stimuli (Vander Weele et al., 2018), and also exhibited activity signatures that underlie compulsive alcohol drinking (Siciliano et al., 2019). During a sensory discrimination task, dorsomedial PFC cells of the same inhibitory neuron classes had more similar encoding properties, but excitatory neurons had diverse task encoding that correlated with different layers that contained heterogeneous populations (Pinto and Dan, 2015). This picture however remains incomplete as the molecular heterogeneity of these populations is unknown and the behavioral task repertoire is diverse, thus making the match between cell type and function a continuing challenge.

Here, we use the mouse PFC as a case study to address the extent to which the encoding of cognitive task-related signals is predicted by transcriptomic and projection properties, starting from the foundation of a transcriptomic analysis. Single-cell RNA-sequencing of neurons labeled by *Rbp4Cre* (most Layer 5 excitatory projection neurons; Gerfen et al., 2013) identified seven transcriptomic types, but projection mapping revealed that most types projected to multiple targets and most targets received projections from multiple types. Building on this foundation, we leveraged a unique property that projection-defined mPFC→PAG (periaqueductal gray) neurons all derived from a single transcriptomic type, and assessed the diversity of task-related signals present in these neurons by performing Ca^2+^ imaging in freely-moving mice during several two-alternative choice tasks. This allowed us to determine the degree of heterogeneity in the signal encoding of these transcriptomically homogenous neurons. We contrasted mPFC→PAG neurons to two other classes of PFC neurons: those that project to contralateral PFC (comprising three transcriptomic types), and those labeled by *Rbp4Cre* (comprising all seven transcriptomic types). The use of single-cell RNA sequencing data to drive the subsequent analysis of encoding properties in cognitive tasks bridges an important gap between molecular and systems neuroscience, furthers our understanding of PFC function, and extrapolates principles of how task information is organized in a cell type framework.

## RESULTS

### Single-cell Transcriptomes of *Rbp4Cre*-labeled Projection Neurons in Prefrontal Cortex

Rodent PFC is distinguished by a lack of Layer 4, and contains a thick Layer 5 that targets diverse subcortical and intracortical areas (Gabbott et al., 2005). We thus focused on using the *Rbp4Cre* line, which is known to label most Layer 5 excitatory projection neurons (Gerfen et al., 2013), as a foundation for our dataset. To define transcriptomic cell types, as well as uncover potential differences in gene expression across PFC regions, we first profiled tissue broadly from dorsomedial (dmPFC), ventromedial (vmPFC), and orbitofrontal (OFC) regions (see STAR Methods for subregion descriptions).

We crossed the *Rbp4Cre* mouse line with a reporter mouse expressing tdTomato in Cre+ cells (*Ai14;* Madisen et al., 2010), dissected and dissociated tissue from postnatal day 34 to 40 (P34–P40) double transgenic progeny, and performed fluorescence-activated cell sorting (FACS) and plate-based single-cell RNA sequencing on tdTomato+ cells using SMART-Seq2 (Picelli et al., 2014) (Figure 1A). We analyzed 3139 high-quality cells (Figure S1A) pooled from all three regions (dmPFC: 910 cells, n = 3 mice; vmPFC: 1234 cells, n = 4 mice; OFC: 995 cells, n = 4 mice) and performed Seurat-based unbiased clustering and batch normalization (Butler et al., 2018; Stuart et al., 2019; STAR Methods) at multiple resolutions (Figure 1E) to understand the organization of cell classification. Classification with a relatively low clustering resolution gave seven clusters represented in tSNE space, each of which was defined on the basis of co-expression of multiple genes. For simplicity, we highlight only one exemplar marker gene from each set of co-expressed genes: *Cd44, Figf, Otof, Pld5, Cxcr7, Npr3,* and *Tshz2,* respectively for the seven clusters (Figures 1B, 1D, Table S1 Tab 1), but emphasize that the definition of these clusters relies on multi-dimensional data.

**Figure 1.**
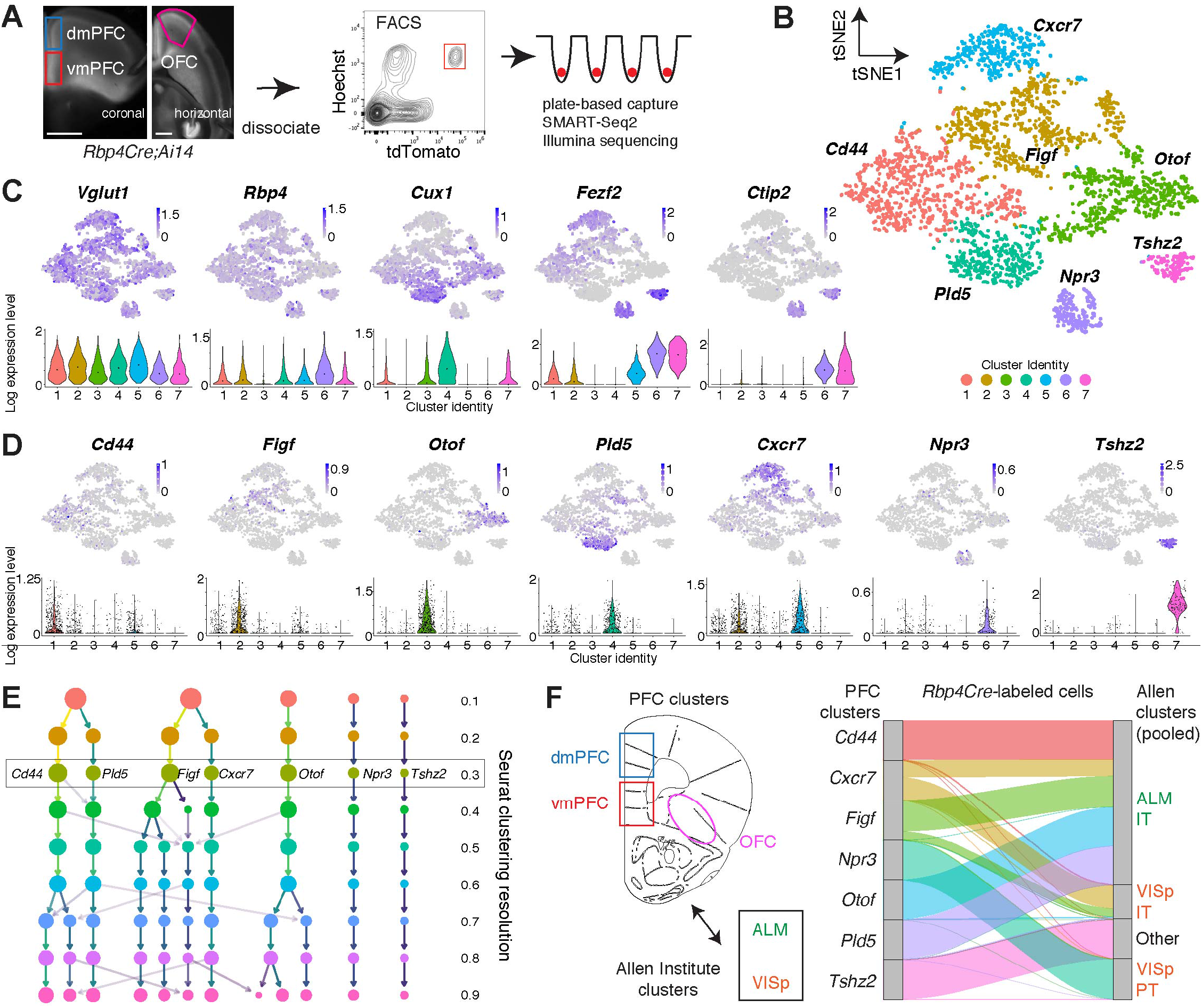
Transcriptomic Map of *Rbp4Cre*-labeled Prefrontal Cortex (PFC) Projection Neurons. (A) Procedure to isolate cells from three PFC subregions in *Rbp4Cre;Ai14* animals for single-cell RNA sequencing. Tissue was dissociated into a single cell suspension, FAC-sorted into either 96- or 384-well plates, and processed using the SMART-Seq2 protocol. Scale bar, 1 mm. (B) Unbiased clustering of 3139 high-quality projection neurons (median ~7000 genes/cell, ~1–2 million reads/cell) based on transcriptomic criteria, shown in dimensionally reduced t-distributed stochastic neighbor embedding (tSNE) space. This embedding was generated using the Seurat pipeline with batch correction (STAR Methods). The seven clusters are labeled based on the expression of marker genes determined from differential expression analysis across clusters. (C) Feature plots in tSNE space (top) and violin plots (bottom) showing the single-cell gene expression of known marker genes, which validate that the dataset is focused on excitatory pyramidal neurons (*Vglut1*+) Neurons are separated by upper vs. deeper layer markers *Cux1* vs. *Fezf2*, and the subcortically-projecting neuron marker *Ctip2*. Color scale of feature plots and y-axis of violin plots in this and other panels are in the unit of ln[1+ (reads per 10000)]. Dots within each violin plot are the median of the distribution. (D) Feature plots in tSNE space and cluster-specific violin plots showing the single-cell gene expression of specific marker genes (STAR Methods) identified to best distinguish clusters through differential expression analysis (see Table S1 Tab 1). Dots within each violin plot are individual data points representing cells. (E) ‘Clustree’ flowchart (Zappia and Oshlack, 2018) showing how cell classifications change across different Seurat clustering resolution parameters (0.1–0.9). Color intensity of arrows indicates the size of the population moving between levels. The relatively low resolution of 0.3 was chosen because it is a level where clusters could still be distinguished on the basis of 1–2 marker genes. Note the relative stability of the *Otof, Pld5, Cxcr7,* and in particular the *Npr3* and *Tshz2* clusters. (F) Determination of Seurat nearest neighbor mapping (TransferData; Stuart et al., 2019) between *Rbp4Cre*-labeled types in PFC (defined here) and *Rbp4Cre*-labeled types in ALM (anterolateral motor cortex) or VISp (primary visual cortex) (defined in Tasic et al., 2018). An alluvial diagram (right) shows the mapping results of how cells from the 7 PFC clusters were categorized on the basis of the 20 ALM and VISp clusters (full list in Figure S1D, summarized into four groups here), with normalization to the same population size for each PFC cluster. IT: intratelencephalic, PT: pyramidal tract (subcortical). See also Figure S1.

All clusters expressed *Slc17a7,* which encodes the vesicular glutamate transporter Vglut1, confirming their excitatory neuron identity. The cluster map could also be divided coarsely into deeply vs. superficially located cells based on expression of marker genes such as *Fezf2* and *Cux1* (Greig et al., 2013; Lein et al., 2007), respectively (Figure 1C). Putative *Ctip2+* subcortically projecting neurons were further divided into two discrete clusters expressing *Npr3* or *Tshz2.* By contrast, gene expression in the remaining five clusters had more continuous variation (Figures 1B, 1D). We refer to these seven clusters operationally as transcriptomic types hereafter. Testing the robustness of this classification using a different method (Allen Institute, scrattch.hicat; Tasic et al., 2018) revealed a similar data structure (Figures S1B, S1C), but with *Figf* and *Cxcr7* clusters more finely subdivided into additional groups.

We observed that few clusters from PFC were highly region-specific even in the case of finely subdivided clusters (Figure S1B), and the majority had more mixed contribution. This was in contrast to a comparison of anterolateral motor (ALM) vs. primary visual (VISp) cortex transcriptomic types, where glutamatergic types were found to be highly region-specific (Tasic et al., 2018). Both of these observations could be explained trivially by continuous spatial differences in gene expression across cortex, which would differentially affect regions that are close together vs. far apart. To examine whether this is the case, we compared *Rbp4Cre*>tdTomato+ cells among PFC and ALM/VISp (Tasic et al., 2018), and determined the cluster from the Allen Institute annotation that was the nearest neighbor to each PFC cell (using TransferData; Stuart et al., 2019). We found that transcriptomically, PFC neurons were not always nearest neighbors with the more physically proximal ALM cells (Figure 1F, ALM mapping: 62.5%, VISp mapping: 37.5%; Allen clusters were pooled based on lists in Figure S1D), indicating that gene expression features defining cortical neuron clusters are not simply explained by physical distance alone. This highlights the complex mapping of homologous cell types between cortical regions and suggests that subtype composition cannot be inferred based only on physical location. However, our data do emphasize that our assayed PFC subregions are relatively similar, and that the most obvious differences are driven by changes in cell proportion.

### Anatomical Locations of Transcriptomic Types

We next examined the spatial location and co-localization of marker expression among the PFC transcriptomic types using hybridization chain reaction-based fluorescence *in situ* hybridization (HCR-FISH; Choi et al., 2018). Focusing on mPFC because of its clear laminar structure in the coronal plane, we quantified marker expression within *Vglut1*-labeled cell soma (Figures 2A, 2B, S2A). *Otof* labeling was highly specific to Layer 2/3 [and negative for *Rbp4* in the sequencing data despite being *Rbp4Cre>tdTomato+* (Figures 1C, 1D), possibly because *Rbp4Cre* is transiently expressed in *Otof+* cells during development] (Figures 2A, 2B, S2A). *Npr3+* and *Tshz2+* cells were located deeper in Layer 5 with similar spatial distributions, but co-localization analysis validated that they were distinct (Figure 2C). Co-localization analysis of *Otof/Figf/Cxcr7* and *Otof/Cd44/Cxcr7* demonstrated a complex variety of single, double, and triple labeled cells distributed throughout Layers 2/3 and 5, consistent with the continuous variation present in the sequencing data (Figures 2D, S2B, similar to Stanley et al., 2020). *Cd44/Cxcr7/Tshz2* triple labeling analysis validated the distinctness of *Tshz2* with the other two markers (Figure S2C). Overall, we found that both transcriptomic and spatial organization was similar between dmPFC and vmPFC, and the major difference distinguishing OFC from mPFC was enrichment of *Pld5* (Figure S2D) and *Cxcr7(5-2)* (Figure S1B) cluster cells. Because vmPFC is narrower than dmPFC, our data indicate that neuronal heterogeneity in mPFC is best summarized as a complex mixture of laminar expression that is increasingly compressed and intermixed from dorsal to ventral, causing extensive overlap of transcriptomic types in Layer 5.

**Figure 2.**
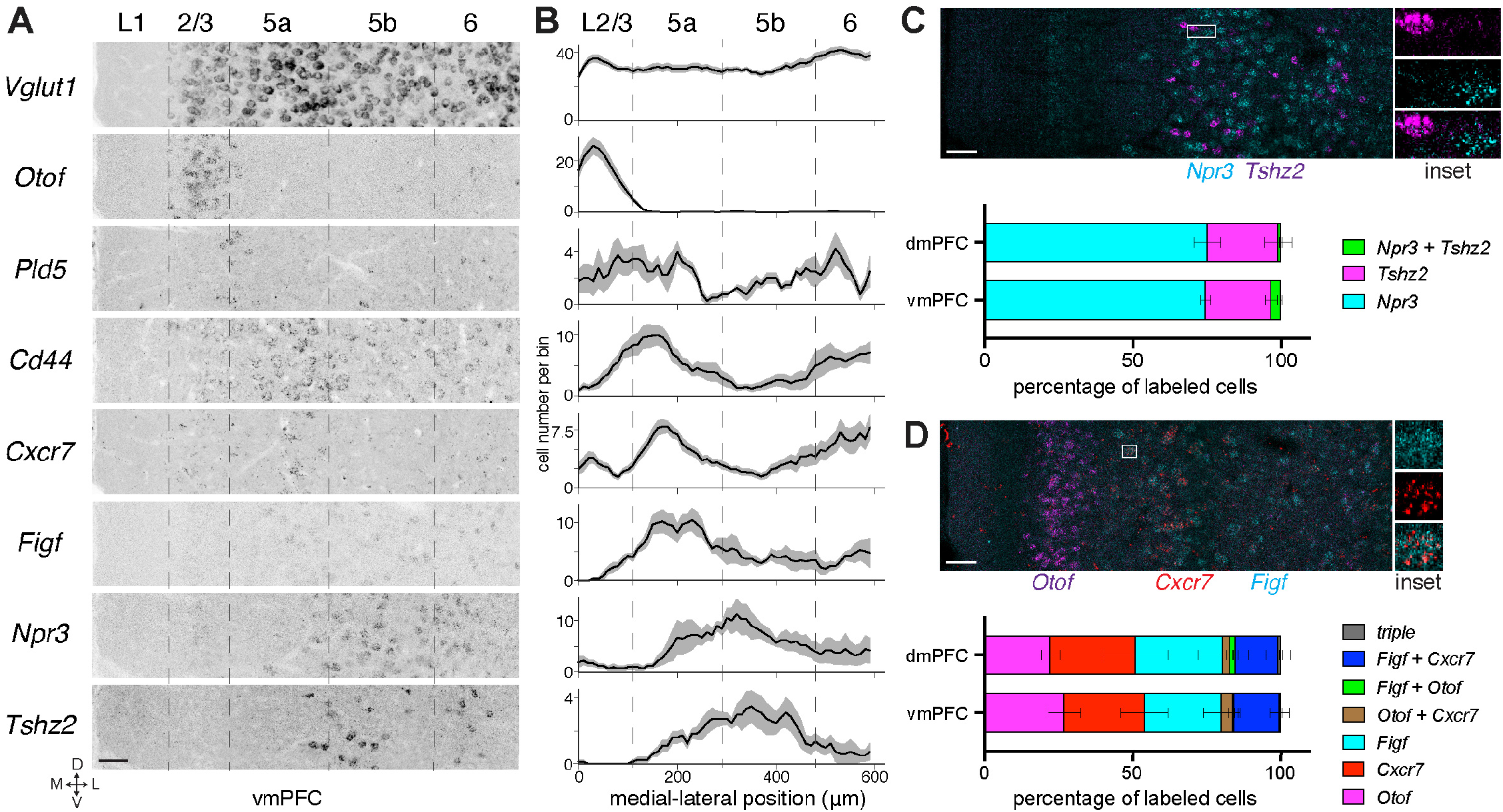
Anatomical Locations of PFC Transcriptomic Types. (A) Images of HCR-FISH of cluster-specific marker genes in vmPFC (A–P ~1.95 mm, D–V 2.35 mm), demonstrating both laminar and non-laminar expression across the cortical wall. Dashed lines are approximate cortical layer boundaries according to the Allen Atlas (beginning of Layer (L) 2/3, 120 μm; L5a, 230 μm; L5b, 410 μm; L6: 600 μm from midline). Scale bar, 50 μm. (B) Laminar distribution of cells expressing cluster-specific marker genes across vmPFC. *Vglut1* counterstaining was used to segment cell soma in order to quantify expression of marker genes. Distributions of cell abundance were aligned medially at the beginning of Layer 2/3, counted with a sliding window (width = 50 μm, step size = 10 μm), and then averaged over the medial–lateral extent of the image (STAR Methods). Traces are averaged across n = 4 animals, with 1–2 images collected per animal. Y-axis represents cells per bin. Layer boundaries are the same as (A) but begin at L2/3. Error bars are SEM. (C) Double HCR-FISH between *Npr3* and *Tshz2* in vmPFC. Quantification of overlap is shown for both dmPFC and vmPFC, and averaged across n = 4 animals (252 dmPFC, 322 vmPFC cells). Error bars are SEM. Scale bar, 50 μm. (D) Triple HCR-FISH between *Otof*, *Cxcr7* and *Figf* in vmPFC, Quantification of overlap is shown for both dmPFC and vmPFC, and averaged across n = 4 animals (540 dmPFC, 591 vmPFC cells). Error bars are SEM. Scale bar, 50 μm. In this and all subsequent figures, all stereotactic coordinates are in millimeters (mm) with respect to bregma. See also Figure S2A–D.

While the exact homology between primate and rodent PFC is a topic of continual debate, in the remainder of this study we focused only on rodent vmPFC (centered on the infralimbic region) because: 1) it fulfills the traditional anatomical definition of PFC in rodents based on dense innervation from mediodorsal thalamus, which is less clearly the case for dmPFC (Rose and Woolsey, 1948; Preuss, 1995; Uylings et al., 2003; Oh et al., 2014); and 2) the full diversity of cell types is more compactly represented in a small space, making tracing and imaging studies more specific.

### Complex Correspondence between Projection Targets and Transcriptomic Types

To examine the relationship between transcriptomic types and specific targets of vmPFC projection neurons, we injected retrograde traveling Cre-expressing virus (*CAV-Cre* or *AAVretro-Cre*, with the caveat of potential differences in viral tropism, see Limitations section of STAR Methods for discussion of this and other caveats), which transduces axon terminals and is transported back to cell bodies, into *Ai14* mice at 6 major known PFC target sites: ipsilateral dorsal striatum (DS), nucleus accumbens (NAc), hypothalamus (Hypo), periaqueductal gray (PAG), amygdala (Amyg), and contralateral PFC (cPFC) (Figure 3A). We dissected vmPFC containing retrogradely-labeled tdTomato+ cells (Figure S2E) and performed single-cell sequencing as before. Appending retrogradely-labeled cells to the dataset from Figure 1 roughly recapitulated the previous clustering, with the exception of a new *Syt6+* cluster derived mostly from hypothalamus-projecting cells and additional *Figf+* cells (Figure S3A). Other than the *Syt6+* (a known Layer 6 marker, Gerfen et al., 2013) cluster, we classified all remaining retrograde cells based on similarity to the seven reference transcriptomic types of Figure 1 (using TransferData; Stuart et al., 2019).

**Figure 3.**
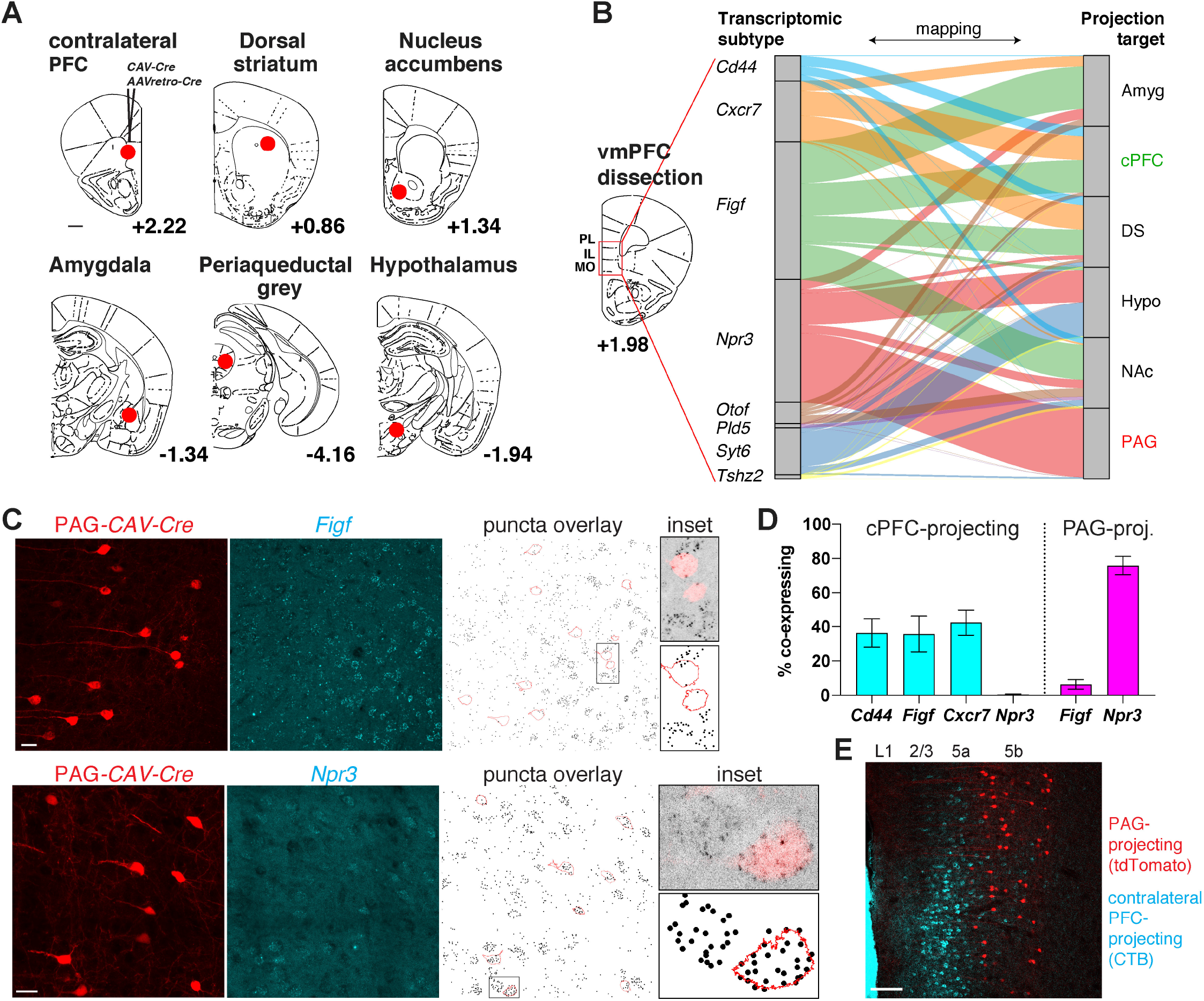
Relationship between Axon Projection Patterns and Transcriptomic Types in vmPFC. (A) Retrograde labeling from major known targets of vmPFC (injection sites indicated by red circles) for singlecell RNAseq, using *CAV-Cre* or *AAV_retro_-Cre* virus injection into *Ai14* animals. tdTomato*+* cells were collected from vmPFC one week after injection for sequencing. Numbers indicate distance in mm in the A–P axis from bregma. Scale bar, 500 μm. (B) Nearest neighbor mapping of retrograde cells collected from vmPFC (query cells: cPFC n = 440, DS n = 129, NAc n = 93, Amyg n = 290, PAG n = 94, Hypo n = 109), to the 7 different transcriptomic cluster identities from Figure 1 (reference), with normalization to the same population size for each projection target. Retrograde cells belonging to a Layer 6 *Syt6* cluster (see Figure S3A) were classified as such, and the remaining retrograde cells were classified using the TransferData function in Seurat against the 7 cluster identities. An alluvial diagram shows the results of the categorization. Similar results were obtained from mapping retrogradely-labeled cells to reference datasets with higher clustering resolution (Figure S3B), of only *Rbp4+* cells, or of only vmPFC cells (data not shown). PL, prelimbic cortex; IL, infralimbic cortex; MO, medial orbital cortex. (C) Confocal images of cells labeled by retrograde tracing from PAG (red) and HCR-FISH *in situ* validation to determine if PAG-projecting cells (tdTomato) express *Figf* or *Npr3* (cyan). HCR-FISH signal was converted to binary puncta and overlaid with tdTomato cell outlines for quantification. Inset shows high magnification of the boxed region. Scale bars: 25 μm. (D) Quantification of the percentage of retrograde cells (cPFC-projecting or PAG-projecting) that co-localized with markers for several transcriptomic clusters (*Cd44, Figf, Cxcr7, Npr3*, n = 3 animals for each). Error bars are SEM. (E) Confocal image showing lack of overlap between PAG-projecting (tdTomato) and cPFC-projecting (CTB-488) cells in vmPFC, labeled in the same section. Scale bar: 100 μm. See also Figures S2E–F, and S3.

To analyze how projection-defined cells were distributed among the transcriptomic types, we visualized the mapping between these two categorical variables (normalized to the size of each projection-defined population). We found that most projection-defined populations consisted of multiple transcriptomic types (Figure 3B, right), indicating that multiple transcriptomic types target individual brain regions. Conversely, each transcriptomic type collectively projects to multiple targets (Figure 3B, left, colored by transcriptomic type). Re-mapping retrogradely-labeled cells to a higher resolution version of the transcriptomic clusters reached similar conclusions (Figure S3B). Despite the divergence and convergence of projections, the targets of any given transcriptomic type exhibited specific biases. Importantly, our data revealed a special case: >97% of vmPFC→PAG neurons mapped to the *Npr3* cluster (15-fold enrichment over random). While we refer to these neurons as ‘PAG-projecting’, it is important to note that PAG is not the only target site. Instead, vmPFC→PAG neurons extend collateral branches to multiple subcortical target sites, including the hypothalamus as predicted in Figure 3B (Vander Weele et al., 2018; and our unpublished data).

We next validated some of these observations by staining sections with PAG-projecting or cPFC-projecting tdTomato*+* cells using markers predicted to either co-label or not. cPFC-projecting cells co-labeled with the markers *Cd44, Cxcr7,* and *Figf* (Figure S3C) at proportions similar to what was observed in the sequencing data, but not with *Npr3* (Figure S3D). Conversely, PAG-projecting cells co-labeled with *Npr3* with high frequency, but not with *Figf* (Figures 3C, 3D) and were distributed throughout mPFC but not OFC (Figure S2F). Finally, double labeling of PAG-projecting and cPFC-projecting cells in the same brain confirmed that these populations did not overlap (Figure 3E), as expected.

Thus far, our study has parsed out different neuronal populations by their transcriptomic and projection signatures. However, the observation that vmPFC→PAG neurons are a highly transcriptomically homogeneous population gave a unique opportunity to examine the signal encoding properties of a neuronal population with high molecular homogeneity. As comparisons to the vmPFC→PAG neurons (hereafter also referred to as the vmPFC→PAG class), we also sought to examine the functional properties of two additional classes: the nonoverlapping vmPFC→cPFC class, consisting mainly of three transcriptomic types (Figure 3B), as well as the entire population of *Rbp4Cre*-labeled neurons as a third class—thereby applying transcriptomic insights to the study of signal encoding.

### Silencing vmPFC Interferes with A Two-Alternative Forced Choice Task

We sought to understand how our characterized organization of vmPFC cell classes and circuit organization contributed to the core function of decision-making (Euston et al., 2012) by imaging neural activity at the single-cell level. We adapted a two-alternative forced choice (2AFC) task in freely moving animals (Figure 4A; Uchida et al., 2003; Feierstein et al., 2006), which could reveal differences in how cells represent diverse sensory, motor, and cognitive signals. We trained water-restricted mice to nose poke into a center port, which triggered the immediate release of one of two odor cues. Each odor was associated with a 4-μL water reward delivered from a port either to the left or right side, which the animal would receive only upon licking the correct port (Figure 4A). The task was self-paced, and mice freely initiated trials and reported decisions. Over a 2–3 week period, mice learned to robustly discriminate the two different odors and earn water rewards (valeric acid or VA → left; 1-hexanol or 1H → right) at high accuracy (>90%) over hundreds of trials (268 ± 10 trials in a single session, mean ± SEM; STAR Methods).

**Figure 4.**
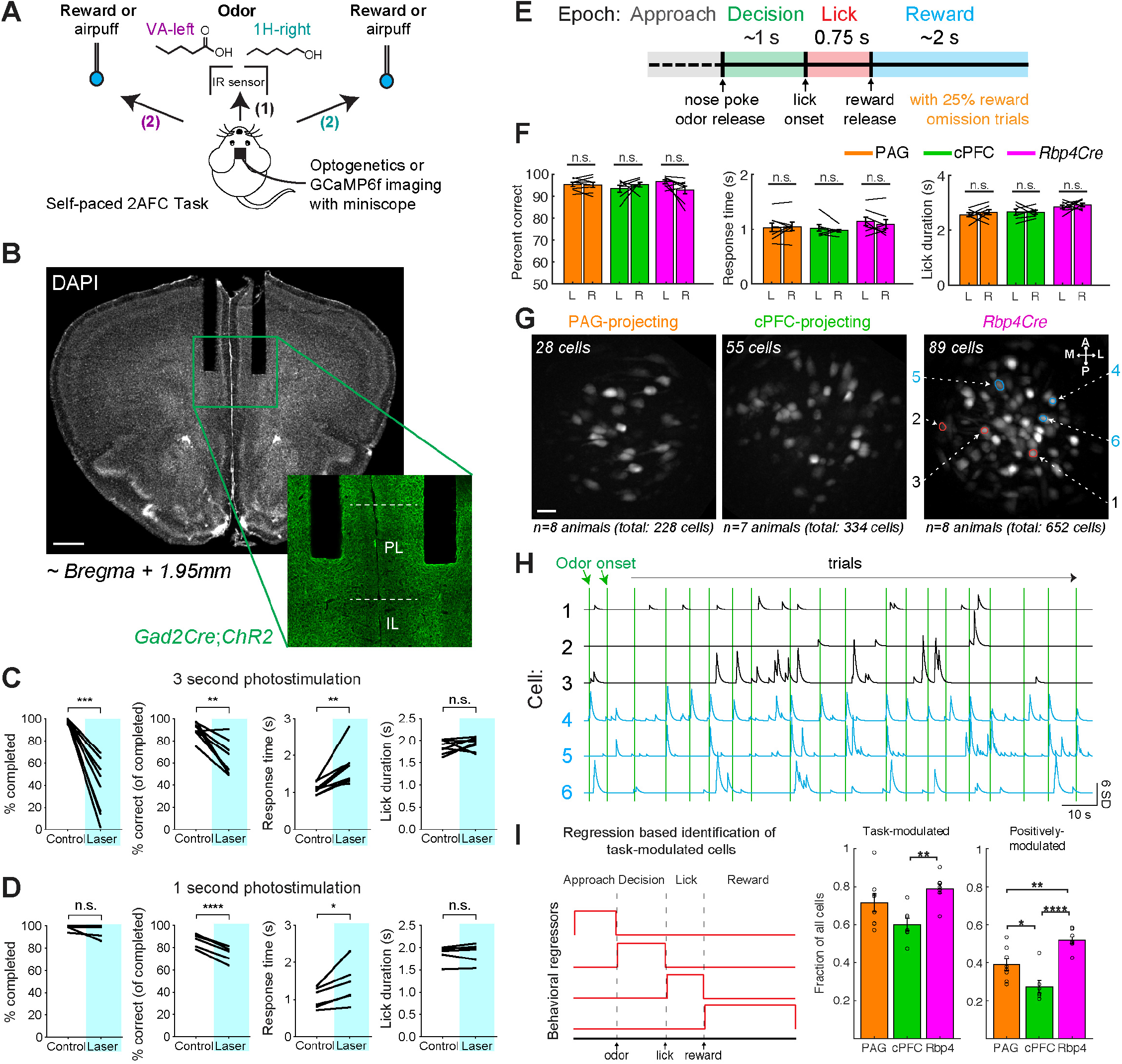
vmPFC is Engaged by A Two-Alternative Forced Choice Task. (A) Self-paced two-alternative forced choice task design for freely moving mice. Mice were trained to nose poke in a center port (1), discriminate between two odor cues (valeric acid or VA → left; 1-hexanol or 1H → right; odor presented for up to 1 s), and move to the correct reward port to obtain a 4-μL water reward (2). Incorrect cue-outcome associations resulted in a brief air puff punishment. (B) Bilateral optogenetic fibers (shown for 200-μm core diameter, 0.39 NA cannulae) implanted into prelimbic cortex (PL; A–P: +1.95, M–L: ±0.35, D–V: −2.3) of animals expressing ChR2(H134R) in all inhibitory neurons (*Gad2Cre:Ai32*). IL, infralimbic cortex. Scale bar: 500 μm. (C) Behavioral effects of vmPFC optogenetic inhibition during task. Photostimulation persisted for 3 s beginning at nose poke (473 nm laser, ~5–6 mW at each fiber tip, 50 Hz, 10 ms pulse width) randomly interleaved on 25% of trials. Trials were considered completed if a response was reported during the allotted 4-s window. Response time was calculated on all completed trials, and lick duration was calculated only for correct trials as animals received air puff punishments on incorrect trials. Paired t-test was used for significance between control and photostimulation trials, *, P < 0.05; **, P < 0.01; ***, P < 0.001; ****, P < 0.0001. (D) Same as (C), but for studies in which photostimulation persisted only for 1 s beginning at nose poke. (E) Trial structure and definition of 4 task epochs for imaging experiments. Water reward was omitted in a random 25% of trials. (F) Performance metrics of animals during imaging demonstrate no sided bias between left and right trial types, on average (*Rbp4Cre:* n = 8, PAG-projecting: n = 8, and cPFC-projecting: n = 7 animals; mean ± SEM, n.s.: not significant, P > 0.05, paired t-test). Left: L, Right: R. (G) Example fields of view for PAG-projecting, cPFC-projecting, and *Rbp4Cre* cell classes, from 2-odor task imaging. Cre-dependent GCaMP6f expression (*Ai148)* was initiated from *CAV-Cre* injections at target sites or from crossing to *Rbp4Cre.* Original images acquired through Inscopix ProView lens probes (500 μm width, 6.1 mm length, 0.5 NA, implanted at A–P: +1.95, M–L: +0.4, D–V: −2.4) using the nVoke endoscopic system. Displayed images are maximum intensity projections of cells in a typical field of view. Colored rings in the *Rbp4Cre* image are example regions-of-interest (ROIs) derived from CNMF-E (see STAR Methods). Scale bar, ~25 μm. (H) Ca^2+^ fluorescence signals (CNMF-E denoised) for six cells highlighted in (G, right panel). Each vertical green line denotes odor onset following the animal’s voluntary nose poke for each trial. Blue traces highlight cells time-locked to the task (see STAR Methods). Black traces are examples from non-time-locked cells. Ca^2+^ signals are z-scored with standard deviation (SD) as the unit. (I) Determination and quantification of cells with task-modulated activity. Set of four behavioral regressors used for linear regression, considered separately for left and right trial types (left). Average fraction of imaged cells that were significantly modulated (middle: both positively and negatively; right: only positively) for each cell class (circles represent individual animals, mean ± SEM; *, P < 0.05; **, P < 0.01; ***, P < 0.001; ****, P < 0.0001; one-way ANOVA, *post-hoc* Tukey’s HSD test; PAG-projecting n = 8, cPFC-projecting n = 7, *Rbp4Cre* n = 8 animals). See also Figure S4.

We first tested whether neural activity in vmPFC of trained animals was important for performance of the 2AFC task. Optogenetic activation of cortical inhibitory neurons has been used as an effective strategy to silence specific cortical areas (Pfeffer et al., 2013; Guo et al., 2014b). We implanted bilateral fiber optic cannulae above vmPFC (Figures 4B, S4A) of *Gad2Cre;Ai32* double transgenic mice (Taniguchi et al., 2011; Madisen et al., 2012), which expresses channnelrhodopsin (ChR2) in all cortical GABAergic inhibitory neurons. We found that photostimulation for 3 seconds beginning at odor onset decreased task performance in multiple ways (Figure 4C). Photostimulation trials had a significantly reduced completion rate: animals often failed to report a choice by licking at any reward port within the allotted time (4 seconds) despite receiving an odor cue. Among completed trials, the error rate and response time were increased. The duration of licking for water reward on correct trials was not affected, however, arguing against a general motor defect. Photostimulation for 1 second beginning at odor onset did not affect the proportion of completed trials, but the error rate and response time were similarly increased (Figure 4D), suggesting disrupted decision-making.

Together, these data suggested that vmPFC is required for the proper execution of the 2AFC task, although we cannot rule out a minor potential contribution of nearby cortical areas from spread of optogenetic stimulation, or distant areas due to the presence of a small fraction of GABAergic projection neurons [He et al., 2016; our own estimate from retrograde-labeling is ~0.5% (5 of 1160 cells)]. This prompted us to characterize the diversity of task-related signals encoded by different classes of projection neurons in vmPFC.

### Imaging Task-Relevant Ca^2+^ Dynamics in Defined vmPFC Classes

We modified the task design for imaging by adding a delay period to better de-correlate signals such as licking and reward consumption. Animals first approached the center port (Approach epoch, ~1 s), discriminated 2 odors to choose between left and right (Decision epoch, ~1 s), reported their choice by licking (Lick epoch, 0.75 s), and received a 4-μL water reward 0.75 s after the first lick (Reward epoch, ~2 s) (Figure 4E). To isolate cell activity correlated with reward, we also randomly omitted reward in 25% of the trials (omission trials) and collectively refer to this design as the ‘2-odor’ task. Animals achieved high levels of performance (>90%) and those that were utilized for imaging on average did not exhibit any significant biases in their performance (accuracy, response time, lick duration) that correlated with the side (Figure 4F).

To determine whether neural activity varied by cell class in vmPFC during task performance, we utilized a *Cre-*dependent GCaMP6f mouse (*Ai148;* Daigle et al., 2018) to label each of the three neuronal classes of interest in separate cohorts of mice: PAG-*CAV-Cre* (one transcriptomic type), cPFC-*CAV-Cre* (three transcriptomic types that exclude the PAG-*CAV-Cre* type), and *Rbp4Cre* (seven transcriptomic types including the above two classes). Experimental animals were implanted with a gradient-index lens (GRIN: 500 μm width) above vmPFC (Figure S4B), through which we performed Ca^2+^ imaging of neural activity at cellular resolution using the Inscopix miniature endoscopic system (Ghosh et al., 2011; Stamatakis et al., 2018). Figures 4G and S4C show fields of view (FOVs) for all imaged mice (*n* = 7–8 animals per cell class). As an example, Figure 4H shows fluorescence traces of 6 cells (FOV in Figure 4G, right panel) whose Ca^2+^ transients correlated (blue) or did not correlate (black) with odor onset.

To determine task-relevant activity, we first defined a set of behavioral regressors representing the four task epochs (Approach, Decision, Lick, and Reward) for left and right trial types separately (Figure 4I, left) and then performed linear regression with the observed cell activity. Cells were considered task-modulated if at least one regression coefficient was significant when compared to shuffled data that randomized the regressor–activity relationship. By quantifying the number of task-modulated cells as a proportion of all imaged cells on a per animal basis, we found that across all three cell classes, ~60–80% of all cells were modulated in at least one epoch, suggesting that all three classes contribute significantly to task representation, but with cPFC-projecting cells less likely to be modulated compared to the other two classes (Figure 4I).

### Cell Classes Are Differentially Recruited by Task Epoch

Do cells of different classes have enriched activity in different epochs? To address this, we first analyzed the Ca^2+^ signal of single cells in individual trials and calculated trial-averaged traces aligned to odor onset or first lick. We found cells with activity significantly elevated during each of the four epochs (Approach, Decision, Lick, Reward) in all three classes of neurons we imaged (Figures 5A, S5A). Thus, there is no absolute relationship between neuron class and the time within the task that it is active. Next, for each class, we identified all positively-modulated cells across all animals, and sorted cells by the time of their maximum trial-averaged activity aligned to odor onset and first lick (Figure 5B). In all three cases, peak response times tiled the entire task. The tiling pattern was not an artifact of sorting, as cross validation of averaged data from only even trials but subsequently aligned to odd trials produced nearly identical patterns (Figure S5B). Thus, even a molecularly homogeneous class of neurons (PAG-projecting) encodes diverse signals.

**Figure 5.**
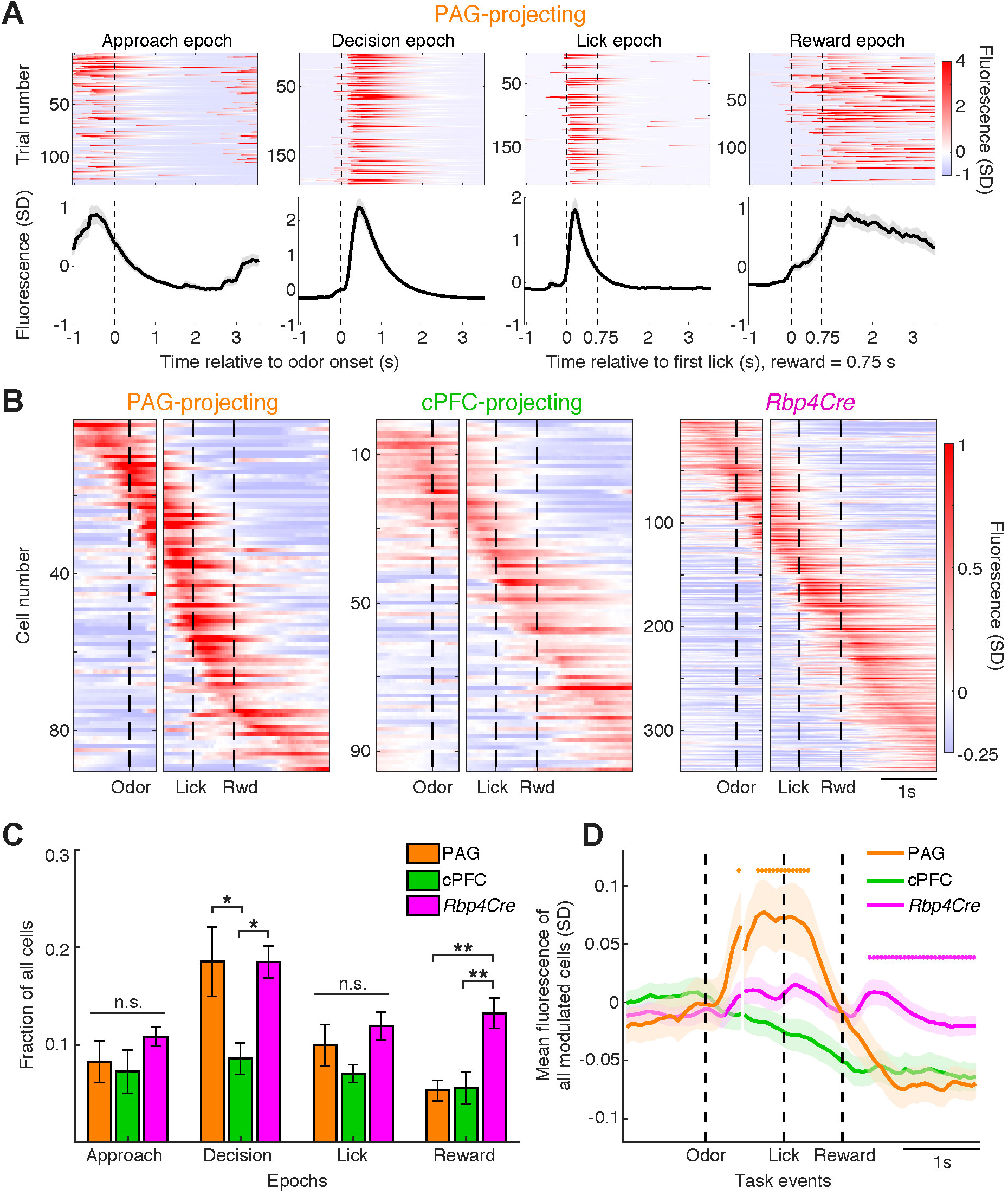
Differential Enrichment of Activity Across Epochs Between Cell Classes. (A) Example single-trial activity (top) and corresponding trial-averaged activity (bottom, mean ± SEM) of four significantly modulated PAG-projecting cells during the four task epochs as defined in Figure 4E. Traces include all correct rewarded trials. Reward was delivered 0.75 s after the first lick. Vertical dashed line in Approach/Decision epochs denotes odor onset. Vertical dashed lines in Lick/Reward epochs denote first lick response and reward delivery. Fluorescence intensity is z-scored, with standard deviation (SD) as the unit. Unless otherwise specified, all activity in this and subsequent figures is represented as the z-score of the fluorescence intensity signal of single cells. (B) Trial-averaged activity of all positively-modulated cells sorted by time of maximum activity, and grouped by cell class. Panels are aligned to odor onset (left) and first lick (right) (n = 90 PAG-projecting cells, n = 95 cPFC-projecting cells, n = 339 *Rbp4Cre*-labeled cells). (C) Cells positively-modulated in each of the four task epochs (Approach, Decision, Lick, Reward) as a fraction of all cells, on a per-animal basis (mean ± SEM; n.s., P > 0.05; *, P < 0.05; **, P < 0.01; ***, P < 0.001, oneway ANOVA, *post-hoc* Tukey’s HSD test). (D) Average activity trace of all task-modulated cells aligned to odor onset and first lick, for each cell class (n = 168 PAG-projecting cells, n = 205 cPFC-projecting cells, n = 518 *Rbp4Cre*-labeled cells; mean ± SEM). Orange and magenta dots represent timepoints where the PAG-projecting and *Rbp4Cre* traces are significantly greater than the other two traces, respectively (P < 0.05, one-way ANOVA, *post-hoc* Tukey’s HSD test). See also Figure S5.

We then classified and counted positively-modulated cells based on the epoch wherein statistical significance was first reached. On a per-animal basis, the PAG-projecting and *Rbp4Cre* classes had a relatively greater proportion of cells modulated within the Decision epoch compared to the cPFC-projecting class. Conversely, the PAG- and cPFC-projecting classes had a significantly lower proportion of cells modulated during the Reward epoch compared to *Rbp4Cre* (Figure 5C). Therefore, although activity of cells in each class tiled the entire trial, the classes varied in the amount of modulation in each epoch. To test whether cell classes exhibit quantitative differences in their activity across task epochs at the population level, we computed the average signal for all task-modulated cells across the three classes (Figure 5D). The *Rbp4Cre* trace exhibited two periods of elevated Ca^2+^ signal: following odor and reward onset. By contrast, the PAG-projecting trace had elevated Ca^2+^ signal after odor onset, whereas the predominant effect in the cPFC-projecting trace was that of negative modulation. Furthermore, the magnitude of the PAG-projecting trace was significantly greater than the other two traces during the Decision and Lick epochs, whereas the *Rbp4Cre* trace was greater during the Reward epoch. Finally, by comparing activity during rewarded (75%) and reward omission trials (25%) (Figures S5C–E), we found that reward omission is represented across each class by a net loss of activity, with the largest change in *Rbp4Cre*-labeled cells. These results demonstrate that activity in each class, despite heterogeneity at the single cell level, has distinct signatures at the population level.

### vmPFC→ PAG Neurons Contain the Most Information about Choice Direction

Choice-specific information has been observed across multiple regions of cortex (Feierstein et al., 2006; Harvey et al., 2012; Guo et al., 2014b; Li et al., 2015; Driscoll et al., 2017; Wagner et al., 2019). Similarly, we found that many vmPFC cells exhibited activity selective not only for the task epoch, but also for left or right choice directions (Figures 6A, S6A). To explore the relative abundance of this important task signal across different cell classes, we first analyzed data at the population level by pooling imaged cells from all animals. To visualize how choice direction-selective activity evolved over time within a typical trial, we computed trial-averaged activity traces for each cell separately on left and right correct trials, and grouped all cells into a time-varying high-dimensional neural activity trajectory, where each axis represents the activity of a single neuron (Churchland et al., 2006; Shenoy et al., 2013). We used principal component analysis (PCA) for dimensionality reduction and plotted neural activity trajectories using the first 3 principal components, which accounted for ~70% of the variance in the data (Figures 6B, S6B). Prior to odor onset, neural trajectories of the left and right trials were very similar. Upon odor onset, these trajectories rapidly diverged, and persisted through the reward epoch.

**Figure 6.**
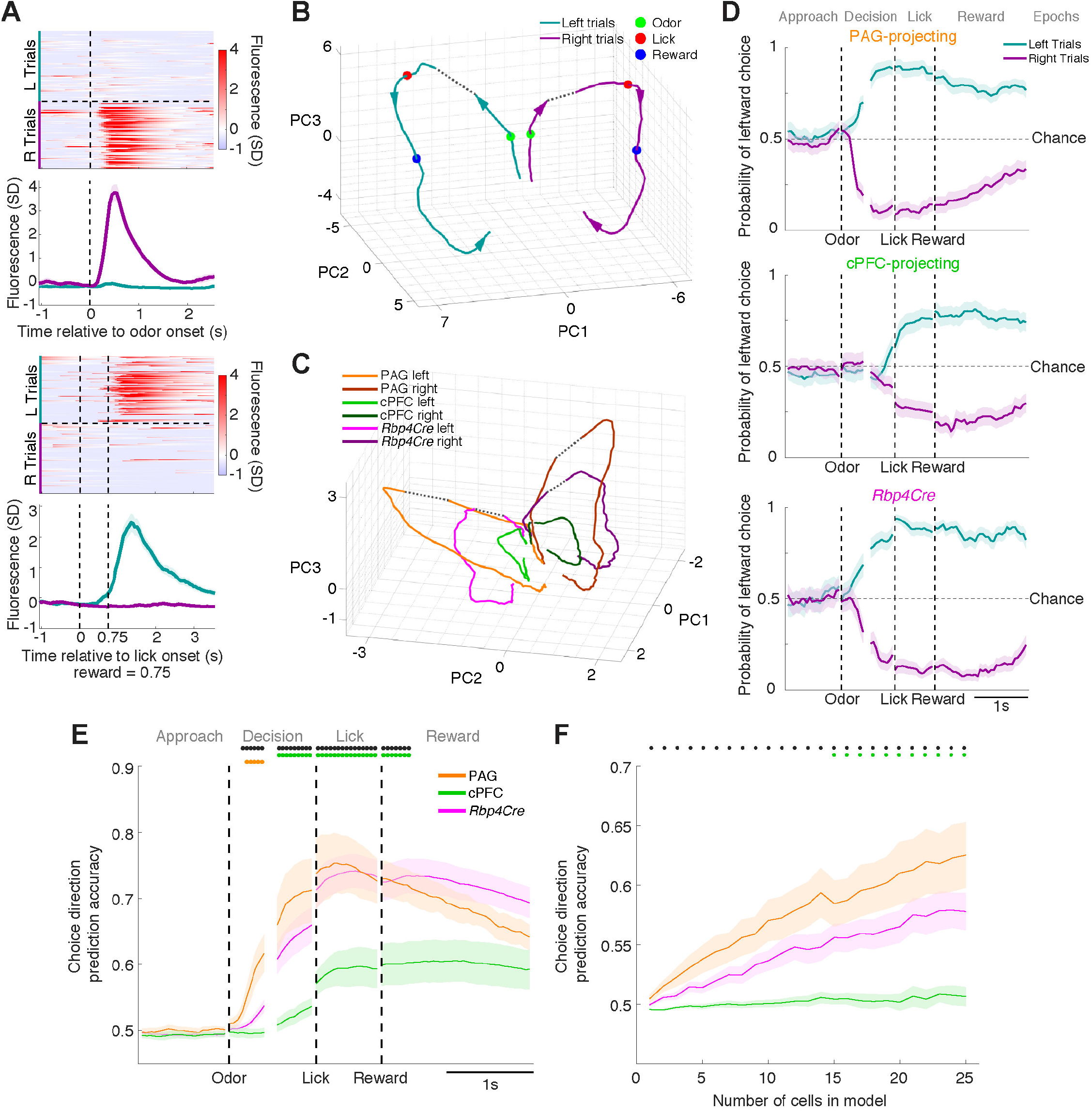
Choice Direction-Specific Information Differs Quantitatively between Cell Classes. (A) Example single-trial activity (upper panels) and corresponding trial-averaged activity (lower panels; mean ± SEM) of two representative cells. The top cell has positive modulation on right trials during the Decision epoch. The bottom cell has positive modulation on left trials during the Reward epoch. (B) Population neural activity trajectories of trial-averaged correct left and right trials represented using the first three principal components (PCs) in activity state space. Arrows denote the direction of time as the animal progresses through the task. Green, red, and blue dots represent onset of odor, lick, and reward delivery. All cells imaged are included regardless of cell class (n = 1214 cells). Dotted lines connect the data between the two alignment points. (C) Similar to (B), except that neural activity trajectories are subdivided by cell class (color-coded) and randomly subsampled to 200 cells for each class. (D) Accuracy of choice direction prediction using a logistic regression model, shown over time across the four epochs, with one example animal for each cell class. Values towards 1 or 0 (mean ± SEM) indicate accurate left or right choice direction prediction, respectively. Models were computed independently for each epoch. Data from both correct and error trials were used in the Approach and Decision epochs. Data from only correct trials was used in the Lick and Reward epochs. (E) Average choice direction prediction accuracy across animals (PAG-projecting n = 5, cPFC-projecting n = 5, and *Rbp4Cre* n = 8 animals), from data randomly subsampled to 25 cells per animal (mean ± SEM). Black dots represent timepoints where the PAG-projecting trace is significantly greater than the cPFC-projecting trace. Orange dots represent timepoints where the PAG-projecting trace is significantly greater than the other two traces, and green dots represent timepoints where the cPFC-projecting trace is significantly lower than the other two traces (P < 0.05, one-way ANOVA, *post-hoc* Tukey’s HSD test). (F) Average choice direction prediction accuracy during the Decision epoch across animals (PAG-projecting n = 5, cPFC-projecting n = 5, and *Rbp4Cre* n = 8 animals), as a function of the number of cells included in the logistic regression analysis (mean ± SEM). Black dots represent cell numbers where the model for PAG-projecting cells is significantly more accurate than the model for cPFC-projecting cells. Green dots represent timepoints where the model for cPFC-projecting cells is significantly less accurate than the other two (P < 0.05, one-way ANOVA, *post-hoc* Tukey’s HSD test). See also Figure S6A–B.

We next asked whether the cell classes contributed differentially to the aggregate trajectory. By using random subsampling to control for differences in the number of imaged cells between classes, we computed trajectories for left and right correct trials within the same PC space, while segregating the data by cell class (Figure 6C). Cell class-specific trajectories all showed a similar profile to the aggregate trajectory, and the number of principal components (dimensionality) required to capture the variance in the data was similar between classes (Figure S6B), again demonstrating that each class had a similar diversity of activity patterns. However, the magnitude of neural trajectory divergence between left and right trials differed: PAG-projecting cells diverged the most after odor onset, followed by *Rbp4Cre*-labeled cells, and then cPFC-projecting cells. This suggested that the PAG-projecting class contains the most choice direction-specific information.

We sought to confirm that these observations held true across individual mice where cells were simultaneously imaged. For each mouse, we performed logistic regression to compute a time-varying prediction of choice direction in each epoch (similar to Kiani et al., 2014, using both correct and error trials, 5-fold cross validation; STAR Methods). Data from PAG-projecting and *Rbp4Cre*-labeled example animals showed that predictions were at chance levels before odor onset, which increased in accuracy after odor onset and lasted through the Reward epoch. Data from cPFC-projecting cells showed a similar pattern, but prediction accuracy improved significantly later, suggesting less choice direction-specific information during the Decision epoch (Figure 6D). To quantify this and compare cell classes, we randomly subsampled the number of cells used in the regression model to 25 per animal, and calculated the average prediction accuracy for choice direction in each class of neurons over time (Figure 6E). This confirmed that the PAG-projecting class contained more information compared to the cPFC-projecting class, particularly during the Decision epoch, with the *Rbp4Cre*-labeled class falling in between. Finally, we asked how this information accumulated over increasing numbers of cells. We computed regression models using data from the Decision epoch that included varying numbers of cells, and plotted the rate at which information accumulated with the number of cells included. Fewer cells in the PAG-projecting class were needed to reach a given prediction accuracy compared with the cPFC-projecting class, indicating a greater amount of information regardless of population size (Figure 6F). Overall, these data demonstrate that PAG-projecting cells more potently encode choice direction compared to cPFC-projecting cells.

### Dissecting Behavioral Variables with Two Additional Tasks

Many of the critical moments of the task occur in the Decision epoch, as this is the period when the animals must interpret the odor identity, make a left vs. right choice, and implement the decision by moving to the correct port. However, because the animals are freely moving, we could not easily dissociate these events in our 2-odor task above. Thus, we devised two additional tasks to analyze these events. First, to determine the extent that different odor cues leading to the same choice outcome are generalized, we designed a 4-odor task: animals were trained to associate one additional odor cue with each side [Left 2: (*S)-*carvone; Right 2: (*R)-*carvone; a pair of enantiomers]. Second, to determine whether activity observed during the Decision epoch was specific to this task, or if similar activity would be evoked whenever a general reward-seeking action toward the left or right ports is made, we designed an uncued task: animals perform similar movements, but have no need to discriminate odors on each trial. During this task, a nose poke at the center port triggered an immediate water reward only at the left port for a block of trials, followed by a block of trials where water reward was only given at the right port.

The same mice that contributed to the 2-odor dataset were trained for an additional 2–3 weeks on the 4-odor task (but not the uncued task), and re-imaged while performing the 4-odor task at expert levels. This was immediately followed by the uncued task within the same imaging session (Figures 7A, S6C). Mice were able to transition quickly from the 4-odor task to the uncued task and attained high performance (>80%) within 10–20 trials (Figure S7A), and performed hundreds of trials in total within one session (165 ± 4 4-odor trials, 50 ± 3 left block trials, 53 ± 3 right block trials, mean ± SEM). We found that the 4-odor task generally recapitulated our findings in the 2-odor task. Sorted, trial-averaged activity in the 4-odor task revealed a similar tiling pattern (Figure 7B) and a largely similar distribution of task-modulated cells across epochs (Figure S7B) as the 2-odor task (Figures 5B, 5C). The ability of different cell classes to predict left vs. right choice directions was also similar to the 2-odor task, with PAG-projecting cells containing more information than cPFC-projecting cells (Figures S7C, S7D).

**Figure 7.**
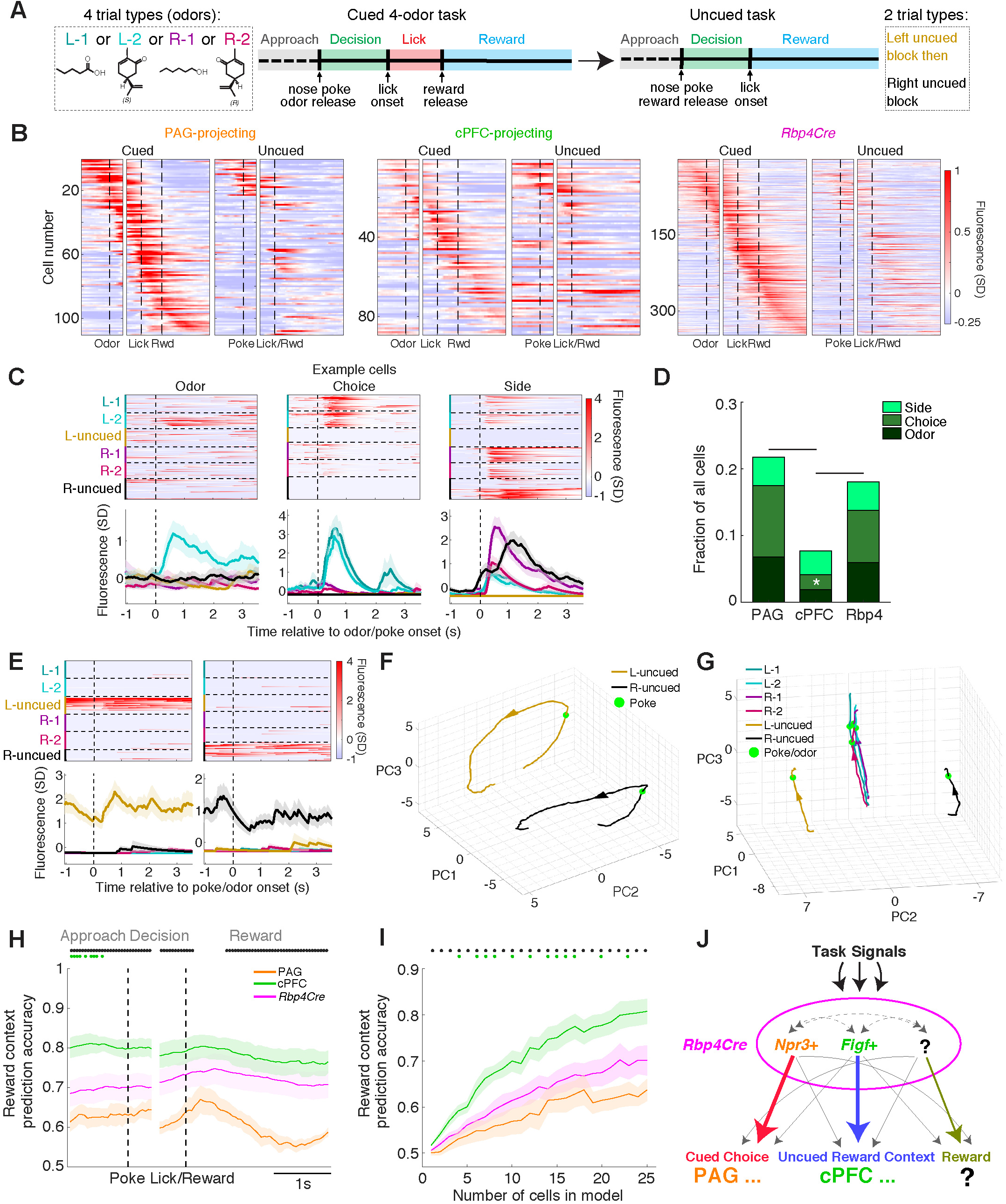
Two Additional Cognitive Tasks Reveal How Cell Classes Differentially Encode Task Signals. (A) Diagram of task design. During imaging, animals first discriminated four possible odors [valeric acid = left 1 or L-1; (*S)*-carvone = L-2; 1-hexanol = R-1; (*R)-*carvone = R-2] to receive 4-μL water rewards. After performing this task, they switched immediately to an uncued task of repeated left (L-uncued) or right (R-uncued) trials in blocks, resulting in six trial types. (B) Trial-averaged activity of all positively-modulated cells during the cued task (left) sorted by the time of maximum activity and grouped by cell class (n = 110 PAG-projecting cells, n = 89 cPFC-projecting cells, n = 348 *Rbp4Cre*-labeled cells) followed by trial-averaged activity of the same cells during the uncued task (right). (C) Example Odor- (L-2 only), Choice- (L-1 + L-2), and Side-selective (R-1 + R-2 + R-uncued) cells with single-trial activity (top) and corresponding trial-averaged activity (bottom, mean ± SEM). Vertical dashed line denotes nose poke/odor onset. (D) Proportions of cells positively-modulated in the Decision epoch, grouped by cell class, and categorized as Odor-, Choice-, or Side-selective. Using linear regression, we determined whether each cell had significant regression coefficients across the six trial types when compared to data that randomized regressor–activity temporal relationships. Cells were categorized based on which coefficient (or group of coefficients) was significant (see STAR Methods). Comparison of the proportion of Choice-selective cells across classes was performed using a permutation test (*, P < 0.05). (E) Example cells with preferential activity during L-uncued trials (left) or R-uncued trials (right). Single-trial activity (top) and corresponding trial-averaged activity (bottom, mean ± SEM) is shown. Vertical dashed line denotes odor/nose poke. (F) Population neural activity trajectories summarizing the trial-averaged traces of left versus right uncued trials represented using the first three principal components (which accounted for >50% of the variance) in activity state space. All imaged cells were included regardless of cell class (n = 1248 cells). (G) Same as (F) except that all six trial types are plotted, and only the Approach epoch leading up to the nose poke is analyzed. (H) Average prediction accuracy of reward context (left vs. right block-type) over the time course of the uncued task, across animals (mean ± SEM; PAG-projecting n = 4, cPFC-projecting n = 5, and *Rbp4Cre* n = 8 animals), from data randomly subsampled to 25 cells per animal (mean ± SEM). Black dots represent timepoints where the cPFC-projecting trace is significantly higher than the PAG-projecting trace, and green dots represent timepoints where the cPFC-projecting trace is significantly higher than the other two traces (P < 0.05; one-way ANOVA, *post-hoc* Tukey’s HSD test). (I) Average prediction accuracy of reward context during the Approach epoch, as a function of the number of cells included in the logistic regression analysis (mean ± SEM). Black dots represent timepoints where the model for cPFC-projecting cells is significantly more accurate than the model for PAG-projecting cells. Green dots represent timepoints where the model for cPFC-projecting cells is significantly more accurate than the other two (P < 0.05, one-way ANOVA, *post-hoc* Tukey’s HSD test). (J) Schematic summary. The heterogeneous population of *Rbp4Cre*-labeled cells in vmPFC is subdivided into cell classes defined by differential gene expression and projection patterns (*Npr3*+; *Figf*+/*Cxcr7*+/*Cd44*+, simplified as *Figf+*), which predominantly route different information (cued choice or uncued reward context) to different targets (PAG or cPFC). However, we note here that PAG and cPFC are not the only sites these neurons project to. Our data also suggest that *Rbp4Cre*-labeled cells contain a subclass that preferentially encodes reward, distinct from PAG- or cPFC-projecting cells. See also Figures S6C and S7.

However, many cells exhibiting task-modulated activity in the 4-odor task exhibited either little task-modulated activity or task-modulated activity with different temporal profiles in the uncued task, supporting the specificity and odor-dependence of the observed neural activity and differences in cognitive demands across the two tasks (Figure 7B). Importantly, this change was not explained by the passage of time, as a comparison of the 4-odor data broken down into first and second halves did not reveal a difference compared with the uncued segment (Figure S7E). Furthermore, inclusion of the uncued data increased the number of principal components required to explain the total variance in the dataset, demonstrating that the uncued task generates activity patterns distinct from the cued task (Figure S7F).

### Differential Encoding of Odor, Choice, and Side Revealed from the 4-Odor and Uncued Tasks

Neural activity enriched in the Decision epoch could be associated with a specific odor (hereafter Odor), both odors for a particular choice direction (hereafter Choice), or movement to one particular side (hereafter Side). The combined data from the 4-odor and uncued tasks provide us with six contrasting trial types to dissociate these variables. Indeed, we found example cells with activity selective for a specific odor, for two odors that predict the reward on a specific side, and for movements towards a specific side, corresponding to the Odor, Choice, and Side definitions above (Figure 7C).

To quantitatively analyze these data, for each cell with positive modulation during the Decision epoch, we performed linear regression analysis with six behavioral regressors, each representing one of the six trial types during the Decision epoch (L-1, L-2, R-1, R-2, L-uncued, R-uncued). We determined which regression coefficient (or group of coefficients) was significant by comparing the true coefficient with those generated from random shuffles of the neural activity traces with respect to the regressors (STAR Methods). If a regression coefficient was significant, we considered that cell modulated by the corresponding trial type. From this, we classified cells into the following categories: Odor (modulated by L-1 or L-2 or R-1 or R-2 regressors), Choice (modulated by L-1 and L-2; or R-1 and R-2 regressors), or Side (modulated by L-1 and L-2 and L-uncued; or R-1 and R-2 and R-uncued regressors) (Figure 7D). Odor-selective cells were present across all three classes and Side-selective cells were in similar proportion across all three classes. Choice-selective cells, however, were represented in PAG-projecting and *Rbp4Cre*-labeled cells at a significantly higher proportion compared to the cPFC-projecting class. These results validated choice as the most abundantly represented signal in the PAG-projecting class.

### vmPFC→ cPFC Neurons Preferentially Represent Reward Context in the Uncued Task

In contrast to the 2- and 4-odor tasks, where the animal is uncommitted to a choice before odor presentation, choice-specific information is potentially represented even before nose poke in the uncued task. By segregating the data into the six different trial types and comparing trial averaged activity, we observed cell examples that exhibited neural activity only during the uncued trials and specific to one of the two block-types (Figure 7E). In contrast to the transient activity characteristic of the cued trials, some of these cells exhibited heightened activity throughout each trial, including the period prior to nose poke (Figure 7E). Representing the aggregate neural data during left vs. right uncued trials as trajectories in activity state space (Figure 7F), we observed that the trajectories never came close to each other. In addition, comparing trajectories during the Approach epoch leading up to nose poke/odor onset across all six trial types demonstrated clear overlap between the cued trials, and segregation between the uncued trial types (Figure 7G). Together, these data demonstrate differences in activity state between the cued and uncued tasks, and provide evidence that information about block-type, or the reward context, is present in vmPFC persistently through each trial.

To address how reward context is represented across cell types in uncued trials, we performed a regression analysis of neural data similar to Figures 6D–F. Left vs. right reward context could be predicted from all three populations, but interestingly, this information was most potent in cPFC-projecting cells (Figures 7H, 7I). The prediction accuracy of reward context by cPFC-projecting cells was consistently higher than that of PAG-projecting cells throughout most of the trial duration, with *Rbp4Cre*-labeled cells falling in between (Figure 7H). Furthermore, the prediction accuracy of reward context by cPFC-projecting cells was generally higher than that of the other two classes across regression models that included varying numbers of cells (Figure 7I). Over the course of this study and prior to the uncued task, we consistently found only weak recruitment of cPFC-projecting cells in the cued tasks, and positively-modulated cPFC cells had no consistent signatures of their behavioral encoding. The results here indicate that cPFC-projecting cells are particularly important in signaling the reward context that the animal is in during the uncued task. Overall, these results emphasize how all three of the cell classes that we studied here have both common information, as well as individual biases in representing specific task signals (Figure 7J).

## DISCUSSION

Molecular neuroscience has invested substantial effort to survey transcriptomic heterogeneity of neurons in a variety of brain regions. On the other hand, systems neuroscience widely ignores cell-type information when positing and testing mechanisms of information encoding in brain circuits, a practice often implicitly justified by the redundant and recurrent nature of neural activity in higher brain regions (Harris and Mrsic-Flogel, 2013; Shenoy et al., 2013). How do we reconcile these contrasting views? Our PFC single-cell dataset uncovered transcriptomic cell types consistent with those from studies that profiled the mouse brain exhaustively (Saunders et al., 2018; Zeisel et al., 2018), other parts of the neocortex (Tasic et al., 2018), or PFC more specifically (Bhattacherjee et al., 2019), but we focused on using Layer 5 pyramidal neuron subtypes and their projection mapping as a case study to interrogate task encoding properties with a cell-type framework. Using PAG-projecting neurons as an example of molecular homogeneity due to its invariant clustering across multiple resolutions and methods (Figures 1E, S1B), we found that even a homogeneous transcriptomic type encodes diverse information. However, different cell classes also preferentially contributed to different aspects of task encoding, suggesting that each transcriptomic type makes quantitatively different contributions to behavior. Given that this mapping is possible even when intentionally focusing on a cortical region known for its complexity, molecular atlas-building efforts (Zeng and Sanes, 2017) will undoubtedly continue to provide a useful framework for analyzing cell function.

### Distribution of Task Information Across Cell Types

Neural responses of individual cells in PFC tend to be highly heterogeneous and represent different combinations of experimental and behavioral variables. This feature, referred to as mixed selectivity, has gained prominence in recent years as a mechanism for how PFC neurons represent task-related signals in a computationally efficient manner to accomplish behavioral tasks (Fusi et al., 2016; Rigotti et al., 2013). Indeed, we found that all three of our examined vmPFC cell classes contained but had relatively small proportions of mixed selective cells, based on their representation of choice direction and/or reward context (Figure S7G; methods similar to Mante et al., 2013). Furthermore, it is also appreciated that task-related signals in goal-directed, cognitive tasks are more generally distributed across the brain than previously thought, in both primate and mouse (e.g., Hernandez et al., 2010; Allen et al., 2017b and 2019; Steinmetz et al., 2019). Our study extends these perspectives by demonstrating that even when looking within a specific brain region, within a single cortical layer, and within a transcriptomically specific cell type, a diversity of information is still present both across the population of cells and within individual cells, including cells that fulfill criteria of mixed selectivity.

Another important feature of our study is that we surveyed the behavioral encoding of our cell classes in detail, across multiple two-alternative choice tasks. By using the 2-odor task to determine general principles of how each class encodes for task signals, and contrasting this with 4-odor and uncued tasks, we obtained insights that could not have been found with only one task. For example, cPFC-projecting cells appeared to be weakly recruited in the 2-odor and 4-odor cued tasks compared with the two other classes. However, our observation that cPFC-projecting cells are most potent at representing reward context in the uncued task is an explicit example of how different cell types come into action during different tasks, underscoring the importance of having sufficient diversity in behavioral repertoire when testing for cell-type specific encoding.

Performance of the uncued task directly after the 4-odor cued task also highlights the flexibility of PFC: choice is more potently represented by PAG-projecting cells in the cued task, but upon changing experimenter-defined rules and switching to the uncued task, animals adapt within tens of seconds, and reward context is more potently represented by cPFC-projecting cells. This ‘division of labor’ between cell types intermingled within the same brain region is echoed at the level of brain regions in a recent study demonstrating that the number of required cortical areas and their dynamics varied across related tasks and the difficulty of cognitive computations (Pinto et al., 2019).

Finally, it is well known that individual neurons within frontal cortices exhibit persistent activity involved in short-term memory (Fuster and Alexander, 1971; Miller et al., 1996). Our finding that reward context is represented preferentially in intracortically-projecting neurons suggests that cortico-cortical networks are likely a key player underlying the maintenance of this activity. While not explicitly tested, our observations may aid in providing a transcriptomic and projection-based cellular context to the body of literature that has linked PFC to working memory (Spellman et al., 2015; Kamigaki and Dan, 2017; Bolkan et al., 2017; Schmitt et al., 2017) and the representation of decision variables such as value (Bari et al., 2019; Hirokawa et al., 2019), both of which could be in play during our uncued task.

### Relationship between Axon Projection Patterns and Transcriptomic Types

The function of neurons is constrained by input–output connectivity patterns. Mesoscale axon collateralization patterns of large neuron populations (Oh et al., 2014; Zingg et al., 2014; Harris et al., 2019) typically present reproducible, highly complex innervation patterns to many areas. However, as these cell populations are subdivided, the extent that axon projection patterns can be subdivided into individual ‘channels’ of output to specific targets, or can retain an overlapping ‘broadcast’ pattern to multiple targets varies substantially across neural systems. For example, projection-defined norepinephrine neurons in the locus coeruleus have diverse and mostly overlapping patterns of output collateralization (Schwarz et al., 2015; Kebschull et al., 2016), whereas projection-defined dopamine neurons in the ventral tegmental area (Beier et al., 2015) or AGRP neurons in the hypothalamus (Betley et al., 2013) have substantially less overlap in their outputs.

A current assumption in the field is that dividing neuron populations based on transcriptomic criteria will be helpful in deciphering the correspondence between transcriptomic vs. projection types. Our data (Figures 3 and S3) indicate that this is not necessarily the case, at least for PFC projection neurons. Based on the mapping of cells between transcriptomic type and downstream target, we find that most regions receive axons from multiple transcriptomic types, and every transcriptomic type projects to multiple targets, at least in part through axon collateralization. This one-to-many and many-to-one pattern undoubtedly underlies the complexity and flexibility of information flow in PFC, and is also in line with similar data focusing on ALM and VISp (Tasic et al., 2018). Emerging evidence from reconstructions of single neuron axon arborization patterns after whole-brain imaging (Lin et al., 2018; Gong et al., 2016; Winnubst et al., 2019; Ren et al., 2019) or sequencing (Kebschull et al., 2016; Han et al., 2018; Chen et al., 2019) demonstrate how axon morphologies of individual neurons can be highly heterogeneous within genetically defined neuronal populations. For example, neurons in the median preoptic nucleus that regulate thirst appear transcriptomically homogeneous, but also target axons to multiple different sites with little collateralization (Allen et al., 2017a). One possible explanation could be that transcriptomic heterogeneity during development that is essential for establishing wiring specificity greatly diminishes in adulthood (Li et al., 2017). Together, these observations suggest a limited extent to which projection patterns can be predicted by transcriptomic data in adult neurons.

In conclusion, a future systems-level understanding of the brain requires integrating multimodal data into a common framework. Our study has aimed to integrate single-cell sequencing and Ca^2+^ imaging data within a highly dynamic behavioral context, and paves the way for future studies that will achieve coregistration of cell type, axon projection, and activity patterns during cognitive behaviors in the same brain.

## STAR METHODS

### Key Resources Table

### Contact for Reagent and Resource Sharing

### Experimental Model and Subject Details

Mice

### Method Details

Single-Cell Sequencing

Histology

Confocal and Slide Scanner Imaging

Surgical Procedures

Behavioral Training

Optogenetics

Collection of Ca^2+^ Imaging Data

Extraction of Ca^2+^ Imaging Data

### Quantification and Statistical Analysis

Single-Cell Sequencing Data Analysis

Differential Expression and Marker Definition

Scrattch.hicat Analysis and Comparison to Seurat

Mapping Query and Reference Datasets

HCR-FISH Puncta Finding and Confocal Imaging Data

Analysis Analysis of Trial-Averaged Activity and Task-Relevant Modulation

Neural Activity Trajectory Analysis

Regression Analysis

Analysis on the Effect of Reward Omission

Classification of Odor-, Choice-, Side-Selective Cells in the Decision Epoch

Classification of Mixed Selective Cells

Statistical Tests

### Limitations

Retrograde Viral Transduction

Optogenetic Perturbation

Age of Mice

### Data and Code Availability

## ACKNOWLEDGMENTS

We thank E. Kremer for the *CAV-Cre* virus; A. Mizrahi, D. Pederick, and L. DeNardo for critical comments on the manuscript; L. Driscoll and members of the Luo Lab for reagents and helpful discussions; L. Schwarz for performing pilot *in situ* hybridization experiments; L. Murrow for advice on RNA sequencing analysis; R. Jones and S. Kolluru for technical help with RNA sequencing; C. Tsang, L. Cardy, and A. Thomas for technical help with the Inscopix microscopes; and M. Andermann for supporting N.D.N. to work on this project after leaving the Luo Lab. J.H.L. is supported by a NIMH K01-MH114022 grant. S.R.Q. is a Biohub investigator. L.L. is a HHMI investigator. This work was also supported by an NIH grant (R01-NS050835) and an NSF NeuroNex grant.

## Author Contributions

J.H.L. and L.L. conceived the project. J.H.L. performed and analyzed the sequencing experiments, performed all of the surgeries, trained animals, and supervised the behavioral experiments and imaging data analysis. N.D.N. designed and wrote all of the code for the behavioral systems, trained animals, performed imaging, and analyzed the imaging data. S.M.G. aided in the optogenetics experiments and performed the HCR-FISH experiments and analysis. S.D. trained J.H.L. to perform the sequencing experiments, and helped in sequencing data analysis. M.J.W. provided critical guidance in imaging data analysis. W.E.A. designed an original version of the olfactometer and behavioral system, and helped in data analysis. J.M.K. aided with sequencing data management and provided critical guidance in sequencing data analysis. E.B.R. aided in image segmentation analyses. J.R. performed the amygdala injections. D.P. and W.T.N. offered critical guidance in the behavioral task design and analysis of imaging data. S.R.Q. supported the sequencing data acquisition, and guided data analysis. L.L. supervised the entire project. J.H.L., N.D.N, and L.L. wrote the manuscript. All authors edited the manuscript.

## STAR METHODS

### Key Resources Table

| Reagent or Resource | Source | Identifier |
| --- | --- | --- |
| **Virus Strains** |  |  |
| CAV-Cre | Soudais et al., 2001 | N/A |
| AAVretro-CMV-Cre-2A-eGFP | Stanford Vector Core | N/A |
| **Antibodies and RNA probes** |  |  |
| anti-GFP chicken polyclonal antibody | Aves Labs | Cat# GFP-1020 |
| Slc17a7 (Vglut1) probe | Molecular Instruments | NM_182993.2 |
| Cd44 probe | Molecular Instruments | NM_009851.2 |
| Vegfd (Figf) probe | Molecular Instruments | NM_010216.2 |
| Otof probe | Molecular Instruments | NM_031875.2 |
| Pld5 probe | Molecular Instruments | NM_176916.4 |
| Ackr3 (Cxcr7) probe | Molecular Instruments | NM_007722.4 |
| Npr3 probe | Molecular Instruments | NM_008728.2 |
| Tshz2 probe | Molecular Instruments | NM_080455.2 |
| **Chemicals** |  |  |
| Isoflurane | Henry Schein | CAS# 26675-46-7; CHEBI: 6015 |
| Avertin (2,2,2-Tribromoethanol) | Sigma | SKU# T48402 |
| Tissue-Plus O.C.T. Compound | Thermo | Cat# 23-730-571 |
| Cholera Toxin B-Alexa488 | Molecular Probes | C-34775 |
| Hoechst | Life Technologies | H3570 |
| DAPI | Thermo | D1306 |
| Papain Dissociation System | Worthington | LK003150 |
| SMARTScribe Reverse Transcriptase | Takara | 639538 |
| KAPA HiFi HotStart ReadyMix | Roche | KK2602 |
| AMPure Beads | Fisher | A63881 |
| Quant-iT™ PicoGreen™ dsDNA Assay Kit | Thermo | P11496 |
| Nextera XT Library Sample Preparation kit | Illumina | FC-131-1096 |
| **Experimental Models: Organisms/Strains** |  |  |
| Mouse: Rbp4-KL100-Cre | GENSAT | KL100 |
| Mouse: Ai14 (Rosa-CAG-LSL-tdTomato-WPRE) | JAX | 7914 |
| Mouse: Gad2-IRES-Cre | JAX | 10802 |
| Mouse: Ai32 (Rosa-CAG-LSL-ChR2(H134R)-EYFP-WPRE) | JAX | 24109 |
| Mouse Ai148 (TIT2L-GC6f-ICL-tTA2)-D | JAX | 30328 |
| Mouse: C57/Bl6 | JAX | 664 |
| Mouse: CD1 | Charles River | N/A |
| **Software and Algorithms** |  |  |
| MATLAB | Mathworks | https://www.mathworks.com |
| Imaris | Bitplane | https://www.bitplane.com |
| Inscopix Data Collection Software | Inscopix | https://www.inscopix.com |
| Inscopix Data Processing Software | Inscopix | https://www.inscopix.com |
| Python | Python | https://www.python.org/ |
| ImageJ (Fiji) | NIH | https://fiji.sc/ |
| Seurat | New York University | https://github.com/satijalab/seurat |
| scrattch.hicat | Allen Brain Institute | https://github.com/AllenInstitute/scrattch.hicat |
| Bpod | CSHL/Sanworks | https://github.com/KepecsLab/Bpod_r0_5 |
| Arduino | Arduino | https://www.arduino.cc |
| SOLIDWORKS | Dassault Systèmes | https://www.solidworks.com |
| Prism | Graphpad | https://www.graphpad.com/scientific-software/prism/ |
| U-Net | UNI Freiburg | https://lmb.informatik.uni-freiburg.de/people/ronneber/u-net/ |
| **Surgical Supplies** |  |  |
| GRIN lens | Inscopix | 1050-002211 |
| Baseplate | Inscopix | 1050-004201 |
| Baseplate cover | Inscopix | 1050-002193 |
| Dummy microscope | Inscopix | 1050-002196 |
| Kwik-Sil | WPI | N/A |
| Kwik-Kast | WPI | N/A |
| Gel Control Superglue | Loctite | 45198 |
| Jet Liquid | Lang Dental | N/A |
| Gel foam | Pfizer | N/A |
| Dual-core implantable Fiber Optic cannulae | ThorLabs | CFM32L10 |
| **Behavioral Training Supplies** |  |  |
| Mineral oil | Fisher Scientific | O122-4 |
| Valeric Acid | Sigma-Aldrich | 240370 |
| 1-hexanol | Sigma-Aldrich | H13303 |
| R-carvone | Sigma-Aldrich | 124931 |
| S-carvone | Sigma-Aldrich | 435759 |
| White acrylic | Plastic-Craft Products | AC1382 |
| Infrared sensor | Adafruit | 2167 |
| Valves | NResearch | 161T011, 161T031 |
| Flowmeter | Cole-Parmer | EW-03227-18 |
| Arduino Due | Arduino | A000062 |
| Arduino TinkerKit | Arduino | K000001 |

### Contact for Reagent and Resource Sharing

Further information and requests for resources and reagents should be directed to and will be fulfilled by the Lead Contact, Liqun Luo.

### Experimental Model and Subject Details

#### Mice

All procedures followed animal care and biosafety guidelines approved by Stanford University’s Administrative Panel on Laboratory Animal Care and Administrative Panel on Biosafety in accordance with NIH guidelines.

To express tdTomato in Layer 5 projection neurons for sequencing, we crossed *Rbp4Cre* (Gerfen et al., 2013, mixed background) mice to *Ai14 (Rosa-CAG-LSL-tdTomato-WPRE,* C57BL/6J background; Madisen et al., 2010) mice. Mice were sacrificed at P34–P40 for single cell isolation and sequencing. A total of 11 mice were used for this purpose.

To express tdTomato in projection-defined neurons for sequencing, we also used *Ai14* mice (mixed CD1, C57BL/6J background), injected *CAV-Cre* (Soudais et al., 2001) at target sites (ipsilateral dorsal striatum, nucleus accumbens, periaqueductal gray, and hypothalamus, as well as contralateral PFC) at P24–P35, and then sacrificed 7 days later at P31–P42, or injected *AAV_retro_-Cre* (Tervo et al., 2016) (amygdala) at P34, and then sacrificed at P49, for single cell isolation and sequencing. Each experiment pooled tissue from 2–4 animals, and each site other than dorsal striatum and nucleus accumbens had two separate batches. A total of 29 mice were used for this purpose. Only female mice were used in sequencing experiments.

To express ChR2 in inhibitory neurons for optogenetic silencing experiments, we crossed *Gad2Cre* (C57BL/6J background; Taniguchi et al., 2011) to *Ai32 (Rosa-CAG-LSL-ChR2(H134R)-EYFP-WPRE,* C57BL/6J background; Madisen et al., 2012) mice. Following this, we performed cannulae implantation, behavioral training, and optogenetics behavioral experiments. A total of 8 mice (3 males, 5 females) were used for this purpose.

To express the Ca^2+^ indicator GCaMP6f (Chen et al., 2013) in neocortical Layer 5 pyramidal cells for imaging, we crossed *Rbp4Cre* (mixed background) to the Cre-dependent GCaMP6f transgenic mouse line *Ai148 (TIT2L-GC6f-ICL-tTA2,* mixed background; Daigle et al., 2018). A total of 8 mice (6 males, 2 females) were used for this purpose. To express GCaMP6f in projection defined neurons for behavioral training and Ca^2+^ imaging, we used *Ai148* mice, injected *CAV-Cre* at target sites at P28–P35, and then performed lens implantation surgeries 1 week later. A total of 15 mice (11 males, 4 females) were used for this purpose. The imbalance in male/female ratio was related to surgery survival rates, and was not intentional. For HCR-FISH validation of sequencing data, adult male and female mice aged P35–P60 on a CD1 and C57BL/6J mixed background were used. Prior to their training on the tasks used to generate the datasets in this study, mice were naive to the behavioral task, and gained their task expertise as described in the ‘Behavioral Training’ section below.

### Method Details

#### Single-Cell Sequencing

The procedure for isolating tdTomato+ cells for single-cell sequencing was identical between those labeled by *Rbp4Cre* (Figures 1, S1) or *CAV-Cre / AAV_retro_-Cre* (Figures 3, S3). Animals were briefly anesthetized with isoflurane and decapitated, and the brain was isolated in ice-cold ACSF (2.5 mM KCl, 7 mM MgCl_2_, 0.5 mM CaCl_2_, 1.3 mM NaH2PO4, 110 mM choline chloride, 25 mM NaHCO3, 1.3 mM sodium ascorbate, 20 mM glucose, 0.6 mM sodium pyruvate, bubbled in 95% O_2_ / 5% CO_2_). Brains were embedded in 3% low-melting point agarose (Fisher BP165-25) in ACSF at 37 °C, cooled to 4 °C, and then cut on a vibratome (Leica VT1200S) in either the coronal (mPFC dissections) or horizontal (OFC dissections) planes into 350-μm floating sections. To microdissect dmPFC, vmPFC, or OFC, we first identified the two adjacent tissue slices (total 700 μm) that most accurately spanned the following anatomical ranges: A–P: ~bregma 1.6 to 2.3 mm (for dmPFC and vmPFC), or D–V: ~bregma −1.5 to −2.2 mm (for OFC). Next we visualized the fluorescent tdTomato labeling and used the atlas as a guide, to cut out the regions of interest as accurately as possible. For dmPFC and vmPFC, we bisected the medial wall (~2.4 mm height × 1 mm width), into upper and lower portions. For *Rbp4Cre*-labeled cells, we collected tissue from both sides of the brain. There are no clear anatomical landmarks that delineate PFC subregions, but based on atlas boundaries, we conservatively estimate that the following subregions are contained in each dissection. dmPFC: cingulate and dorsal prelimbic cortex; vmPFC: ventral prelimbic, infralimbic, and medial orbital cortex; OFC: ventral and lateral orbital cortex. For retrogradely-labeled cells, we only collected cells from the vmPFC ipsilateral to the injection site, except for contralateral PFC injections. Microdissected tissue was incubated at 37 °C in papain enzyme mix + 800 nM kyneurenic acid (Worthington) for 30 minutes, and triturated gently with a P200 pipette every 15 minutes thereafter until fully dissociated, usually within 1 hour of total incubation time. The cell suspension was spun down at 350 *g* for 10 min at room temperature, neutralized with ovomucoid inhibitor, spun again, washed in ACSF, stained with Hoechst for 10 minutes (1:2000, Life Technologies: H3570), washed, filtered (Falcon 532235), and resuspended in 2 mL ACSF.

FACS was performed using the Sony SH800 system with a 130-μm nozzle suitable for the large size of pyramidal neurons. Singlet cells were selected based on low FSC-W, and gated on Hoechst (nuclear stain that penetrates cell membrane) and tdTomato double positivity to identify labeled healthy neurons. Cells fulfilling these criteria were over 100× brighter than background, and were unambiguously identifiable. Single cells were sorted at a low flow rate (<100 events/second), and at the highest purity setting (Single Cell) into 96- or 384-well hard shell PCR plates (BioRad HSP9601 or HSP3901) containing 4 or 0.4 μl lysis buffer [0.5 U Recombinant RNase Inhibitor (Takara Bio, 2313B), 0.0625% TritonX-100 (Sigma, 93443-100ML), 3.125 mM dNTP mix (Thermo Fisher, R0193), 3.125 μM Oligo-dT_30_VN (Integrated DNA Technologies, 5’AAGCAGTGGTATCAACGCAGAGTACT30VN-3’) and 1:600,000 ERCC RNA spike-in mix (Thermo Fisher, 4456740)] in each well, respectively. Following FACS, plates were spun down, sealed and stored at −80 °C.

cDNA synthesis and library preparation protocols were adapted from the SMART-Seq2 protocol (Picelli et al., 2014). 96-well vs. 384-well processing utilized 4 μl or 0.4 μl starting volumes, respectively, and will hereafter be referred to as 1 unit. Plates were first thawed on ice followed by primer annealing (72 °C, for 3 minutes, then on ice). For reverse transcription, 1.5 units of reaction mix [16.7 U/μL SMARTScribe Reverse Transcriptase (Takara Bio, 639538), 1.67 U/μL Recombinant RNase Inhibitor (Takara Bio, 2313B), 1.67× First-Strand Buffer (Takara Bio, 639538), 1.67 μM TSO (Exiqon, 5’-AAGCAGTGGTATCAACGCAGAGTGAATrGrGrG-3’), 8.33 mM dithiothreitol (Bioworld, 40420001-1), 1.67 M Betaine (Sigma, B0300-5VL) and 10 mM MgCl_2_ (Sigma, M1028-10X1ML)], was added to each well either manually (96-well) or with a Formulatrix Mantis liquid handler (384-well). The reaction was then carried out by incubating wells on a thermocycler (Bio-Rad) at 42 °C for 90 min, and stopped by heating at 70 °C for 5 min. Subsequently, 3.75 units of PCR mix [1.67× KAPA HiFi HotStart ReadyMix (Kapa Biosystems, KK2602), 0.17 μM IS PCR primer (IDT, 5’-AAGCAGTGGTAT CAACGCAGAGT-3’), and 0.038 U/μL Lambda Exonuclease (NEB, M0262L)] was added to each well. PCR was then performed using the following program: 1) 37 °C for 30 min, 2) 95 °C for 3 min, 3) 21 cycles of 98 °C for 20 s, 67 °C for 15 s and 72 °C for 4 min, and 4) 72 °C for 5 min. For 96-well plate samples, cDNA from every well was purified using 0.7× AMPure beads (Fisher, A63881), quantified by a Fragment Analyzer (AATI), and diluted to 0.15 ng/μL in Tris-EDTA before tagmentation. For 384-well samples, cDNA from every well was quantified using Quant-iT™ PicoGreen™ dsDNA Assay Kit (Thermo Fisher: P11496), and diluted to 0.4 ng/μL in Tris-EDTA before tagmentation.

For both 96-well and 384-well samples, before tagmentation, we reformatted the samples into a standardized 384-well format, and used the Formulatrix Mantis and Mosquito (TTP Labtech) to automatically perform all liquid handling steps. Tagmentation was performed on double-stranded cDNA using the Nextera XT Library Sample Preparation kit (Illumina, FC-131-1096). Each well was mixed with 0.8 μL Nextera tagmentation DNA buffer and 0.4 μL Tn5 enzyme, then incubated at 55 °C for 10 min. The reaction was stopped by adding 0.4 μL Neutralization Buffer and centrifuging at room temperature at 3,220 *g* for 5 min. Indexing PCR reactions were performed by adding 0.4 μL of 5 μM i5 indexing primer, 0.4 μL of 5 μM i7 indexing primer, and 1.2 μL of Nextera NPM mix. PCR amplification was carried out using the following program: 1) 72 °C for 3 min, 2) 95 °C for 30 s, 3) 12 cycles of 95 °C for 10 s, 55 °C for 30 s and 72 °C for 1 min, and 4) 72 °C for 5 min. After library preparation, wells of each 384-library plate were pooled using a Mosquito liquid handler, and consolidated into one tube. Pooling was followed by two final purifications using 0.8× AMPure beads (Fisher, A63881).

Library quality was assessed using capillary electrophoresis on a Fragment Analyzer (AATI), and libraries were quantified by qPCR (Kapa Biosystems, KK4923) on a CFX96 Touch Real-Time PCR Detection System (Bio-Rad). Libraries were sequenced on NextSeq 500 or NovaSeq 6000 Sequencing Systems (Illumina) using 2 × 75-bp or 2 × 100-bp paired-end reads, respectively. Sequences were de-multiplexed using bcl2fastq. Reads were aligned to the mouse mm10 genome (with Cre and tdTomato genes added) using STAR version 2.5.4 (Dobin et al., 2013). Gene counts were produced using HTseq version 0.10.0 (Anders et al., 2015), for only exons, with the ‘intersection-strict’ flag.

#### Histology

Adult mice were perfused transcardially with phosphate buffered saline (PBS) and 4% paraformaldehyde (PFA). Brains were extracted, post-fixed overnight in 4% PFA, cryoprotected in 30% sucrose/PBS for 48 hours, embedded in OCT, snap-frozen, and stored at −80 °C. For HCR-FISH (Figures 2A, 2C, 2D, S2B, S2C, S2D, 3C, S3C, S3D), 20-μm frozen sections were cut on a cryostat and dried on slides for 4 hours. For immunolabeling and other non-HCR-FISH histology (Figures 3E, S2E, S2F, 4B, S4A, S4B), 50-μm floating sections were cut on a cryostat.

For staining HCR-FISH slides, all probe and wash reagents were from Molecular Instruments, and glassware baked at 180 °C was used. Samples were treated with 4% PFA for 15 min, PBS for 5 min, Proteinase K buffer (1:100 1M TrisHCl, 1:500 0.5 M EDTA, and 14 μg/mL Proteinase K, in dH_2_O) for 6 min, 4% PFA for 10 min, and PBS for 10 min. Slides were then placed in a 37 °C chamber humidified with a 50:50 formamide:dH_2_O mixture and 400 μL of probe hybridization buffer was applied to each slide. After 20 min of pre-hybridization, 400 μL of probe mixture (*Pld5, Tshz2, Cxcr7, Npr3:* 4 nM; *Vglut1:* 8 nM; *Otof:* 10 nM; *Figf, Cd44:* 20 nM) was applied to each slide and incubated for 12–16 hours in the humidified chamber at 37 °C. Slides were then washed in a series of probe wash buffer and 5× SSC-T mixtures (1:0, 3:1, 1:1, 1:3, and 0:1) for 15 minutes each, at 37 °C. Slides were washed again in 5× SSC-T for 5 min before applying 400 μL of amplification buffer to each slide and incubating in the humidified chamber for 30 min. Amplification hairpins were heated quickly and cooled slowly (95 °C for 90 sec, then 30 min at room temperature shielded from light) and mixed into amplification buffer (*Pld5, Tshz2, Cxcr7, Vglut1, Otof:* 50 nM; *Npr3, Figf, Cd44*: 120 nM). 150 μL of amplification mixture was added to each slide, cover slipped, and incubated in a dark humidified chamber at room temperature for 12–16 hours. Finally, slides were immersed in 5× SSC-T for 30 min to remove the coverslips, and washed in 5× SSC-T, before applying the final coverslip.

For immunolabeling (Figure 4B), 50-μm sections collected into PBS were blocked in 10% normal donkey serum (NDS: Jackson)/PBS/0.3%Triton-X overnight at 4 °C, washed 3× in PBS-T (PBS/0.1%Triton-X), incubated in primary antibody [1:1000 chicken anti-GFP (Aves)/5%NDS/PBS/0.3%Triton-X] for 2 days at 4 °C, washed 3× in PBS-T, incubated in secondary antibody (1:500 anti-chicken Alexa488 (Jackson) /5%NDS/PBS/0.3%Triton-X) overnight at 4 °C, washed 3× in PBS-T, adding 1:10000 DAPI in the final wash, and coverslipped.

#### Confocal and Slide Scanner Imaging

For images centered on PFC, we focused on bregma A–P: +1.95mm, as dmPFC, vmPFC, and OFC are all well represented at that coronal level. HCR-FISH images were collected on a Zeiss LSM780 confocal microscope at 20× (Figures 2, S2) or 40× (Figures 3, S3) at 1024 × 1024 or 2048 × 2048 resolution using standard settings, with as many tiles as needed to cover the area for quantification. Images used for quantification in Figures 2 and S2 were single planes. Images used for quantifying co-localization of tdTomato with HCR-FISH in Figures 3 and S3 were maximum intensity projections of 5 images spanning ~20 μm. Other histological images of GFP staining, endogenous tdTomato or endogenous GCaMP6f were collected at 20× using standard settings. The images in Figure 1A for guiding dissections were collected on a dissecting scope with an epifluorescence camera. The images in Figure S2E for the confirmation of injection location were collected at 5× on a Leica Ariol slide scanner with the SL200 slide loader.

For quantification of HCR-FISH in laminar analysis, regions of interest capturing dmPFC (300 μm height × 900 μm width, beginning ~1.1 mm below bregma, which corresponds to cingulate cortex) or vmPFC (300 μm height × 750 μm width, beginning ~2.3mm below bregma, which corresponds to infralimbic cortex) were generated from n = 4 animals (2–3 images per animal). For co-localization, images were generated from n = 3 animals (2–4 images per animal). For OFC, *Pld5* and *Vglut1* staining was quantified in the same manner from regions of interest the same size as and from the same section as vmPFC (300 μm height × 750 μm width), and compared in pairwise fashion over n = 4 animals.

#### Surgical Procedures

We anesthetized mice using isoflurane (1.25–2.5% in 0.7–1.3 L/min of O_2_) during surgeries. We immobilized the head in a stereotaxic apparatus (Kopf Instruments), cleaned the skin with Betadine, injected lidocaine (2%, ~0.3 mL) subcutaneously for local anesthesia, cut open the scalp, peeled back connective tissue/muscle and dried the skull. All virus and dye injections were done at a rate of 100 nL/min. After surgery, animals were injected with carprofen (5 mg/kg), 0.9% saline (1–2 mL/100g body weight), and BuprenorphineSR (0.1 mg/kg) for anti-inflammation, hydration, and pain management, respectively. Mice recovered on a heated pad until ambulatory, were returned to their homecage, and housed in a regular 12 h dark/light cycle with food and water *ad libitum*, unless otherwise noted.

For retrograde labeling experiments, *Ai14* mice were injected with *CAV-Cre* (200–300 nL, 4.2 × 10^12^ gc/mL, all sites but amygdala) or *AAV_retro_-Cre* (300 nL, 8.7 × 10^12^ gc/ml, amygdala only) into contralateral PFC (cPFC: M–L/A–P/D–V, −0.6/+2.2/-2.5), ipsilateral dorsal striatum (DS: +1.7/+0.85–2.8), nucleus accumbens (NAc: +0.6/+1.33/−4.8), amygdala (Amyg: +2.86/−1.3/−4.55), periaqueductal gray (PAG: +0.4/−4.15/−2.8), and hypothalamus (Hypo: +0.55/−1.91/−5.0). For testing injection sites, 100 nL CTB-Alexa488 (Molecular Probes: C-34775) was injected into each site into wild-type mice, and animals were immediately sacrificed for visualization. The images from Figure S2E (top) are from these experiments. The image from Figure 3E results from a dual injection of *CAV-Cre* into PAG, and CTB-Alexa488 into cPFC, with 1-week survival. For insertion of optogenetic cannulae, *Gad2Cre;Ai32* mice (expressing ChR2(H134R)) were implanted with bilateral optical fibers (200-μm core diameter, 0.39NA cannulae, 700-μm spacing that were cut down manually to ~4 mm length, ThorLabs) at the upper border of vmPFC (prelimbic cortex; A–P: +1.95, M–L: ±0.35, D–V: −2.3), for stimulation centered on infralimbic cortex below. The holes in the skull were covered with Kwik-Sil (WPI) for protection, and the cannulae was then secured with superglue (Loctite Gel Control) and dental Jet Liquid (Lang Dental), sealing all of the exposed skull.

GRIN lens implantation surgeries were performed using Resendez et al., 2016 as a guide. In brief, the skull was thoroughly cleaned and roughened with a scalpel blade, and two small screws (stainless eyeglass screws, 1 mm length) were screwed into the skull over posterior areas of cortex, without penetrating the dura, to lend extra support for the headcap. A ~1 × 1 mm craniotomy was cut over the lens target area, which was cleared of any remaining bone and overlying dura using fine forceps. Bleeding was limited with usage of gelfoam surgical sponge (Pfizer), and no further action was taken until bleeding had completely stopped. To visually identify the implantation location, we inserted an empty glass pipette (typically for viral injections) to 75% of the depth of the lens implantation. This served the purpose of creating a ‘starter’ hole, but no further aspiration of brain tissue was performed. Following starter hole generation, a ProView GRIN lens (500 μm width, 6.1 mm length) was loaded onto the ProView lens holder and attached Inscopix nVoke mini-endoscope. Together, this unit would be inserted into the brain with the camera functioning, to assess whether GCaMP6f-labeled cells were present at the final depth. The lens was then centered on bregma, and lowered slowly into the brain at the target location (in mm) A–P: +1.95, M–L: 0.4, D–V: −2.1, at a rate of ~100 μm/min by the stereotax. Once the intended depth was reached, and GCaMP6f labeled cells were confirmed in the field of view, we then proceeded to finalize lens placement by permanently gluing the lens itself in that location to form the ‘headcap’. First, Kwik-Sil (WPI) was used to cover the remaining exposed brain, and subsequently, liberal amounts of superglue (Loctite Gel Control), which was cured using dental Jet Liquid (Lang Dental), was used to firmly attach the lens to the skull and screws, sealing the skin and lens and skull together. After this, the camera and lens holder were carefully released from the lens, and Kwik-Kast (WPI) was used to protect the exposed surface of the glass. After 2 weeks of recovery, animals were re-examined for the presence of GCaMP6f-labeled cells at the lens tip, and if satisfactory, had a permanent baseplate and removable baseplate cover attached (Loctite Gel Control) to the headcap to serve as an adapter of fixed focal distance between camera and lens.

To determine the implant locations of optogenetic cannulae and GRIN lenses in animals after experiments (Figures 4B, S4A, S4B), animals were perfused as described above, but the brain was not isolated from the skull immediately. Rather, the entire head with the cannulae and lens still implanted was processed with 4% PFA and 30% sucrose, to ensure that tracks would remain fixed in place. Hardware was then removed from brains, which were then processed as described above for floating sections. Every section containing a lens or cannulae track was collected, to identify the location of the center point, which was then scored against the standard atlas (Paxinos and Franklin, 2001). Animals for which the center point of the lens or cannulae was outside of prelimbic cortex (not within A–P: 1.7 to 2.1 mm, D–V −1.9 to −2.6 mm) were excluded from analysis.

#### Behavioral Training

Animals were trained to perform at high levels of accuracy on a two-alternative forced choice task in two groups: 1) cannulae-implanted animals for optogenetics experiments (3-s or 1-s laser session), and 2) lens-implanted animals for Ca^2+^ imaging (2-odor task, 4-odor task, and uncued task). In general, mice were water restricted to 1 mL per day and monitored daily to ensure general health by visual inspection and maintenance of >85% of their original weight. Mice were typically able to acquire a minimum amount of water (1 mL) during daily training sessions, and if not, the remainder was supplemented after training. See optogenetics section below for details specific to the optogenetics experiments. Starting one week after the baseplating surgery, all imaging mice were singly housed and chronically wore dummy microscopes with similar weight (1.8 g) and size (8.8 mm × 15 mm × 22 mm) to the Inscopix nVoke miniscope to become accustomed for imaging sessions. The only times the dummy microscopes were removed was to replace with the nVoke miniscope for data collection.

All behavior was performed in custom-built behavioral rigs and controlled by software adapted from the open source Bpod behavioral control system (https://github.com/KepecsLab/Bpod_r0_5). The behavioral box was designed in 3D CAD software with dimensions similar to home cages, and consisted of three main ports: one center odor port and two water delivery ports on the left and right side. The two water ports are 7 cm apart, with the odor port in the center. All ports were at 3.5 cm above the bottom of the box (Figure 4A). Odors were delivered using a custom-built olfactometer. Water was delivered through metal ports coupled to a capacitive sensor that recorded licking at 40 Hz. An IR-sensor was placed in front of the center odor port to control trial initiation. Finally, we also included an air puff port immediately adjacent to each water delivery port to provide mild punishments specific to either side. The behavioral training was adapted from Guo et al., 2014a and Feierstein et al., 2006. Behavioral studies were computer-automated without experimenter input. Three weeks post-lens implantation surgery, or one week post-optogenetic cannulae implantation surgery, water-restricted mice were first habituated to the behavioral rig for two days (~45 minutes per day). During habituation days, mice were given water *ad libitum* through either water port. Following habituation, mice were trained on the two-alternative forced choice task in stepwise protocols with increasing complexity:

1. automatic water rewards dispensed whenever nose poke / IR beam break occurred
2. automatic water rewards coupled to specific odor-side pairings (left with 10% valeric acid in mineral oil and right with 100% 1-hexanol)
3. water reward dispensed only if mice reported the correct odor-side association by licking, but without any consequence for incorrect choices
4. incorrect choices punished by air puff, but mice were allowed to recover and report the correct choice after an initial incorrect choice
5. incorrect choices resulted in a single air puff and termination of the trial

Within ~2–3 weeks of training, mice were able to perform the basic version of the two-alternative forced choice task for optogenetics experiments (Figures 4C, 4D). In this scenario, a typical trial consisted of:

1. nose poking at the center odor port (trial start)
2. immediate release of valeric acid or 1-hexanol for up to one second
3. animals reporting their choice by licking at either the left or right reward port within four seconds from trial start
4. a correct choice resulting in a 4-μL water reward and an incorrect choice resulting in an air puff punishment. No response within the time allotted (4 seconds) terminated the trial.

In contrast to animals used for optogenetics experiments, lens-implanted mice used for imaging were additionally trained to expect water rewards to be delivered following a fixed 750-ms delay after the initial lick response. Mice were imaged on the 2-odor task with reward delay, and 25% of rewards were also randomly omitted (omission trials). For these imaging sessions (Figures 4–6) mice performed at high accuracy (>90%) over hundreds of trials (268 ± 10 trials, mean ± SEM) (Figure 4F). Reward omission trials only occurred during imaging sessions, and did not occur during training.

Following successful imaging during the 2-odor task, mice began training on an analogous 4-odor task. The original two odors (10% valeric acid and 100% 1-hexanol) were replaced with 100% (*R)*-carvone and 100% (*S)*-carvone. Mice trained on the new odor-side pairings until high performance was reached without reward delay or reward omission (~1–2 additional weeks). Following this, the two original odors were reintroduced, and mice were required to discriminate four odors with valeric acid or (*S)*-carvone signaling left and 1-hexanol or (*R)*-carvone signaling right (Figure 7A). Mice typically retained the old odor-side associations. Once high performance was attained, reward delay was reintroduced.

During the 4-odor imaging session, mice performed the 4-odor task at high accuracy (>88%) for 165 ± 4 trials (mean ± SEM), and were then immediately introduced to a novel task variation consisting of a block of ~50 uncued left trials followed by a block of ~60 uncued right trials to provide a contrasting cognitive context for the animals (Figure 7A). During these uncued blocks, nose pokes did not trigger odor release. Rather, each nose poke triggered the automatic release of a water reward at one port, which was repeated over multiple trials. Within 10-20 trials of the start of each block, mice followed a continuous and stereotyped sequence of nose poking at the center port, approaching the reward port, licking/consuming water, approaching the center port, and so on (Figure S7A). Mice on average performed dozens of trials (50 ± 3 left block trials, 53 ± 3 right block trials, mean ± SEM) during imaging. Most mice were able to successfully complete more than 20 correct trials of each block-type during the first introduction of the uncued task. However, 9 out of 23 mice were unable to complete enough trials on the uncued task the first time. These mice were subsequently re-imaged one or two additional times until at least 20 left and 20 right uncued trials were completed correctly within the imaging session. One mouse was never able to complete enough uncued trials. Only data from the mice and sessions where both cued and uncued trials were completed sufficiently (22/23 mice, 1 session each) were combined for the analysis in Figure 7. A small fraction of mice (5/23) were able to complete more than two blocks, and in this case, trials of the same block-type were pooled together.

We quantified animal behavior during trials in terms of percent complete, percent correct, response time, and lick duration (Figures 4C, 4D, 4F). Percent complete is the percentage of trials in which mice reported a response during the allotted 4 second time window after nose poke. Incomplete trials were those where mice nose poked to begin the trial but failed to report a response within 4 seconds. Percent correct is the percentage of trials, out of completed trials, that mice made the correct choice. Incorrect trials are completed trials with the wrong response, which are non-overlapping with incomplete trials. Response time is the time from odor onset to lick response. Lick duration is the time from first to last lick on correct trials. Unless otherwise specified, we only analyzed ‘correct’ trials from the 2- and 4-odor imaging sessions, where mice made the correct choice and remained licking at the reward port throughout the delay period.

During the uncued task, correct trials were those where mice reported their response by licking at the correct port first and within the allotted 4 second time window. To determine the behavioral performance of mice during the transition from the 4-odor task to the uncued task, we calculated the mean performance of mice as they were performing the left and right blocks. Mice were able to achieve high performance in the first left block within ~10 trials and subsequently in the right block within ~20 trials (Figure S7A). The transition between the 4-odor and uncued tasks was immediate (after ~165 4-odor cued trials), and data from both tasks was collected from the same continuous video stream.

#### Optogenetics

Before each session, expert mice were briefly restrained to attach the optogenetics patch cable, and allowed to recover for >30 minutes. During each optogenetics behavioral session, laser stimulation was triggered by the nose poke, and designed to last for 1 or 3 s (separate sessions: Figures 4D, 4C). One second was chosen because the mean response time during normal conditions was approximately one second. Three seconds was chosen as a longer stimulation period that would still allow sufficient time for the mouse to report a choice within the 4 second allotted time. The stimulation laser (473 nm wavelength) was pulsed at 50 Hz with 10 ms pulse durations at an optical power of ~5–6mW at the output of each fiber tip. Based on these settings, the vast majority of laser light is concentrated within 500 μm of the fiber tip. Laser stimulation occurred on 25% of randomly interleaved trials during the task and stimulation lengths were kept constant in each session. Behavioral performance was quantified as described above.

#### Collection of Ca^2+^ Imaging Data

We performed all Ca^2+^ imaging using the Inscopix nVoke miniscope (Figures 4G, S4C, S6C), without refocusing the microscopes across the surgeries and two imaging sessions. This ensured that the same field of view was reproduced over time, to the best of our abilities. However, due to the ~2–3 week period between imaging sessions and minor changes to the fields of view during this time, we were not confident in explicitly aligning cells across imaging sessions and so the datasets were analyzed independently. Before imaging, mice were anesthetized with 2% isoflurane for 2 minutes to attach the miniscope and allowed to recover for 30–60 minutes. To reduce stress, we did not head-fix mice at any point in this procedure.

#### Extraction of Ca^2+^ Imaging Data

Images (360 × 270 pixels) were acquired at 20.01 Hz, LED power of 1 mW/mm^2^, with a gain of 3.0. Each imaging session typically lasted for 30–40 minutes, which began with the self-initiated IR beam break in the animal’s behavior. The resulting imaging data was 2x spatially downsampled and motion corrected using default settings in the Inscopix Data Processing Software. The imaging data was exported, and individual neurons and their respective fluorescence traces were identified using the constrained nonnegative matrix factorization for microendoscopic data (CNMF-E) algorithm (Zhou et al., 2018). For each neuron, the denoised fluorescence trace or the deconvolved Ca^2+^ events was extracted using the OASIS algorithm [AR(1) model, Friedrich et al., 2017]. Every extracted cell was manually checked for circular spatial footprints and Ca^2+^ transients characterized by sharp rises and slow decays. Extracted cells were also excluded if they could not be clearly cross-checked between the correlation and maximum intensity images due to low signal. Elongated-shaped ROIs representing dendrites that correlated with the more circular-shaped cell bodies were removed from analysis. All analysis used either the standard deviation of the denoised fluorescence trace of each cell or the corresponding deconvolved Ca^2+^ events. We used denoised fluorescence traces for the general display of all data over time and logistic regression analysis. We used deconvolved Ca^2+^ events for the determination of task-modulated cells, as well as for all other linear regression-based analysis in Figures S5D, 7D, and S7G. Under our imaging conditions, GCaMP6f transients are likely the result of multiple spikes (Chen et al., 2013).

### Quantification and Statistical Analysis

#### Single-Cell Sequencing Data Analysis

To generate the transcriptomic map featured in Figure 1, we used standard procedures for filtering, variable gene selection, dimensionality reduction, and clustering in Seurat v3.0 (Butler et al., 2018; Stuart et al., 2019). The analyzed *Rbp4Cre>tdTomato+* cells originated from 11 different samples/batches (dmPFC, n = 3; vmPFC, n = 4; OFC, n = 4). Cells were removed if they expressed fewer than 2000 genes, and genes were removed if they were detected in fewer than 3 cells. Cells expressing the inhibitory neuron marker *Gad2* (<1%) were also removed from consideration. This resulted in a dataset of 3139 cells × 17535 genes. All cells were processed by the same SMART-Seq2 chemistry, but were collected in 96- or 384-format, and sequenced on either the NextSeq 500 or NovaSeq 6000 systems. Thus, we used batch correction within Seurat v3.0 to sequentially define pairwise anchors across the 11 batches (Stuart et al., 2019), and integrate the data together to remove possible batch effects. The assumption that there were physiologically analogous cells across batches should apply because the batches were experimental replicates of adjacent PFC subregions. In brief, each batch/sample was split into its own Seurat object, 3000 variable genes were selected with ‘vst’, anchors were found using FindIntegrationAnchors (k.anchor = 5, k.filter = 50, k.score = 30, dims = 1:15, max.features = 100, anchor.features = 3000) and the data was integrated with IntegrateData (k.weight = 100, dims = 1:15). In the integrated object, counts were log-normalized for each cell using the natural logarithm of (1 + counts per 10000), and scaled using ScaleData while regressing out the effects of the # of genes and the # of reads. Cells were visualized using a 2-dimensional t-distributed Stochastic Neighbour Embedding (tSNE, van der Maaten and Hinton, 2008) of the PC-projected data using the FeaturePlot and VlnPlot functions. PCA was performed on the integrated data (npcs = 15), and 8 PCs were used in FindNeighbors based on a steep dropoff in variance explained by visual inspection of the elbow plot. We used the FindClusters function in Seurat v3.0, and compared the resultant clusters from 9 different levels of resolution (0.1–0.9, 0.1 increments) using the ‘Clustree’ function (Zappia and Oshlack, 2018) to visualize the way in which individual cells either retained or changed their classifications as the resolution parameter was increased (Figure 1E). We eventually settled on a resolution parameter of 0.3 for the majority of the paper, after noting that most clusters classified at this resolution were stable at higher resolution, but could be defined, from a practical standpoint, by individual markers. We note that there are no cases where major and unexpected rearrangements occur in the organization, and that only the *Cd44*, *Figf*, and *Otof* clusters are further split at higher resolution. However, given that in the retrograde mapping studies shown in Figures 3 and S3, we saw one-to-many and many-to-one matching between transcriptomic types and projection types, we did not intentionally further subdivide these clusters.

To generate the transcriptomic map used to visualize intermingling of retrograde (n = 1155) cells and *Rbp4Cre*>tdTomato+ (n = 3139) cells (Figure S3A), it was not possible to perform a batch correction based on every individual sample because the cell number in some of the retrograde-labeling samples was too low. Therefore, we performed batch correction after segregating the samples by ‘sequencer’, and then regressed out the effect of ‘plate’ (96-vs. 384-well) subsequently in the ScaleData function. All other parameters were the same, and this transcriptomic map was primarily for the purpose of visualization.

#### Differential Expression and Marker Definition

We used the FindAllMarkers function in Seurat v3.0 and applied this to the ‘RNA’ assay in the Seurat object of 3139 high-quality cells classified at Seurat resolution = 0.3 (Figure 1B). This function performs differential expression analysis (Wilcoxon rank sum) between a specific cluster, compared with all other cells not in the cluster, and then iterates through all seven clusters. Because of the immense number of marker genes with statistically significant p-values, we filtered genes not only on the basis of differential expression levels and average log fold change (avg_logfc > 0.1), but also on the consistency in which a gene was expressed across different cells in the same cluster or population. ‘PCT’ refers to the percentage of cells within a population expressing a specific gene. We first required that the difference in this membership between the assayed cluster and all other cells (PCT_delta_ = PCT_cluster_–PCT_other_) for each considered gene to be >0.35. We next required that PCT_cluster_> 0.5, meaning that a gene must be expressed in greater than 50% of the cells in a cluster, in order to be considered a marker for that cluster. Conversely, PCT_other_ was required to be <0.25, meaning that at maximum, 25% of cells outside of the cluster can express the marker gene. Together, these requirements enforce that marker genes exhibit close to binary ‘on-off expression in and out of the cluster. This resulted in a list of 133 genes (Cluster 1: 6 genes; Cluster 2: 4 genes; Cluster 3: 17 genes; Cluster 4: 7 genes; Cluster 5: 14 genes; Cluster 6: 39 genes; Cluster 7: 46 genes, see Table S1 Tab 1). With this list, we visually inspected available online *in situ* hybridization resources, further prioritized genes based on expression level differences, binary expression (PCTcluster close to 1, and PCTother close to 0), and practical considerations of whether commercially available HCR probes gave good signal-to-noise ratio in histological tissue. From this, we settled on the combination of genes: *Cd44, Figf, Otof, Pld5, Cxcr7, Npr3,* and *Tshz2* to delineate the seven clusters, but recognize that this is not the only combination that achieves this purpose.

We also computed differentially expressed genes for the same dataset, but at a higher clustering resolution [Seurat resolution = 0.9 (Table S1 Tab 2)]. Here, we were interested more broadly in differentially expressed genes rather than only ‘on-off’ expression. Thus, we lowered the threshold to PCT_delta_ > 0.25, and removed the requirement for PCTother. This resulted in a longer list of 867 genes. However, we highlighted genes that fulfilled the criteria of Tab 1 (avg_logfc > 0.1, PCT_cluster_ > 0.5, PCT_other_ < 0.25, and PCT_delta_ > 0.35) in red in Tab 2.

#### Scrattch.hicat Analysis and Comparison to Seurat

We used scrattch.hicat analysis (Tasic et al., 2018) as a contrasting method to analyze and validate our *Rbp4Cre*>tdTomato+ PFC clusters (Figures S1B, S1C). As input, we used the same filtered data matrix from Seurat (3139 cells × 17535 genes), and the 0.3 resolution Seurat cluster labels as a reference classification. In brief, raw counts were normalized to log2(1+ counts per million), and the threshold for differential expression and clustering was set based on the following combination of parameters: padj.th=0.05, lfc.th=1, low.th=1, q1.th=0.5, q2.th=NULL, q.diff.th=0.7, min.cells=15, de.score.th=100. We adhered to these parameters as they were the recommendation by the Allen Institute for data of this size and complexity. We next removed technical artifacts associated with the # of genes, # of reads, type of plate, and the specific sample, to mirror the quality control done in Seurat. Finally, we performed consensus clustering over 20 iterations, which resulted in a new classification that could be compared with the Seurat (resolution = 0.3) classification (Figure S1B), and also visualize the probabilities of cell co-clustering over the different iterations (Figure S1C).

To compare our PFC data to the published *Rbp4Cre-*labeled cell data from ALM and VISp (Tasic et al., 2018), we first preprocessed the raw data to make it compatible with our data. In brief, we downloaded (https://portal.brain-map.org/atlases-and-data/rnaseq) and considered only SMART-Seq2 data, reads mapped to exons, and cells derived from *Rbp4Cre* animals. We filtered cells and detected genes by the same criteria as above (genes detected in >3 cells, cells expressing > 2000 genes), and then only considered genes expressed across all three datasets. Finally, cells labeled from Allen cluster annotations as ‘doublets’, ‘high intronic’, or ‘low quality’ were removed and not considered. This resulted in a dataset of 14611 considered genes, and n = 3137 PFC, n = 546 ALM, n = 697 VISp neurons, which was used for the mapping analysis in Figure 1F.

#### Mapping Query and Reference Datasets

We used Seurat v3.0 to assign cells with classification labels based on their proximity to previously classified cells within a reference dataset. For mapping *Rbp4Cre*-labeled PFC cells to Allen ALM and VISp labels, we first generated a Seurat object containing only the ALM and VISp cells to serve as a reference (data normalization, variable feature finding, data scaling, and PCA done as described above). We next subsetted the PFC data by cluster (*Cd44, Figf, Otof, Pld5, Cxcr7, Npr3, or Tshz2*), and queried each of these subsets with the reference data. We used FindTransferAnchors (dims=1:9, k.filter=25) and TransferData (dims 1:9) to first find anchors between datasets, and then assign classification labels, respectively. To generate Figure 1F, an alluvial diagram was made using the ‘ggalluvial’ R package, based on the number of cells that were assigned to different cluster labels, and normalized to the same number of cells in each PFC cluster. Reference classifications in Figure 1F were pooled from the lists of individual cluster labels shown in Figure S1D. The data in Figures 3B and S3B was generated in a similar way, but query sets were retrogradely-labeled cells from each individual target. The reference classifications were the integrated *Rbp4Cre>*tdTomato*+* Seurat object from Figure 1, at 0.3 (Figure 3B) or 0.9 (Figure S3B) clustering resolutions, and normalized to the number of retrogradely-labeled cells.

#### HCR-FISH Puncta Finding and Confocal Imaging Data Analysis

To quantify the expression of each marker gene and its co-localization, we used *Slc17a7 (Vglut1)* as a counterstain because it is a marker of excitatory neurons. We trained the deep learning network U-Net (Falk et al., 2019) to identify and segment *Vglut1+* cells into individual ROIs, using a total of 12 (1077–2409 wide × 120-693 pixels tall) manually annotated images as training data (using the default caffemodel as input, 5000 iterations, validation interval of 20, 348 × 348 tile size). For determining marker gene expression levels, fluorescent puncta were identified using the Fiji plugin TrackMate (Tinevez et al., 2017: Laplacian of Gaussian detector with sigma suited to each marker’s punctum size). These puncta were binarized and the individual masks for each neuron from the U-Net segmentation were applied, allowing the area and position of each punctum for each gene to be determined. Using custom python scripts, puncta size was normalized to the minimum size observed in each image, and the number of puncta in a neuron was normalized to the maximum in each image. This resulted in a comparable score of the amount of gene expression in each neuron across images and genes. A threshold was set to consider any neuron to be positive for the marker of interest if it contained enough puncta to be within 75% of the maximum number of puncta observed. Additionally, for stains of multiple markers, co-localization was calculated in the same manner with a percentage breakdown of all single, double, and triple labeled cells. To illustrate the medial–lateral spatial distribution of markers, we counted positive cells using a sliding window of 50 μm width and 10 μm per slide. For each image, we identified the first bin that contained >17 (vmPFC) or >11 (dmPFC) cells for 3 consecutive bins, as we found empirically that this was a robust method for defining the transition between the very sparse cells found in L1, and the much denser cells found in L2/3. Subsequently, all per bin cell counts were aligned based on the ‘beginning of L2/3’ as an anatomical landmark close to the midline. The traces of each image were then averaged and plotted in Figures 2B and S2A.

Co-localization of marker genes with tdTomato+ cells (Figures 3C, 3D, S3C, S3D) was processed in a similar manner, but *Slc17a7* and U-Net was not used as the basis for defining cell masks. Instead, we thresholded tdTomato signal to include only the cell body, and then converted these into cell outlines using the ‘Analyze Particles’ function in Fiji. Fluorescent puncta were identified as described above with TrackMate, overlaid with tdTomato cell outlines, and the number of puncta was quantified per cell. Similar to above, a cell was considered staining positive if it contained within 75% of the maximum number of puncta observed within a tdTomato+ cell in the image. If there was low to no co-localization, and hence no tdTomato cell clearly contained puncta in the image, then this maximum was calculated based on tdTomato–cells. The 75% threshold was defined from careful inspection of the data, as it was effective in ruling out puncta contributions from slightly overlapping neighboring cells or background staining, but sensitive enough to pick up lowly expressed genes that did fill the cell outlines. For each combination of marker gene and retrograde labeling, 2–4 windows of 440 × 600 μm size were scored for n = 3 animals, in vmPFC.

#### Analysis of Trial-Averaged Activity and Task-Relevant Modulation

Mice were freely moving and made decisions with slightly variable response times. To assess the temporal specificity of activity in each cell relative to the task structure, we computed trial-averaged activity using the denoised fluorescence trace of each neuron over all considered trials (all correct trials, or all left trials, etc.), and aligned to either the odor onset or the first lick times (Figures 5A, S5A, 6A). To determine which cells had significant task-modulated activity within a particular trial type (e.g., left trials), we defined a set of behavioral regressors representing the four task epochs (Approach, Decision, Lick, and Reward), and for left and right trial types separately (Figure 4I). For each cell, we linearly regressed its deconvolved Ca^2+^ events during the entire recording period onto the set of eight (2-odor task) or 16 (4-odor task) behavioral regressors. Cells with significant regression coefficients were then considered ‘modulated’ in that epoch. The set of all cells modulated in any task epoch was considered the population of ‘task-modulated’ cells. To determine whether a cell had a significant regression coefficient for any behavioral regressor, we used permutation tests. We randomized the regressor–activity relationship by shuffling the neural activity matrix with respect to the regressors, and performed linear regression to produce coefficients derived from the shuffled data. If the true coefficient was greater (or lower) than the shuffled coefficient for a given cell for over 99% of iterations, it was considered significant. Positively-modulated cells were defined as those which exhibited higher regression coefficients than chance and negatively-modulated cells were defined as those which exhibited lower regression coefficients than chance. Cells with significant modulation in two consecutive epochs (for example, Decision and Lick) were categorized by the first significant epoch. Cells with significant modulation for multiple trial types (for example, right Decision *and* left Reward) counted twice for the calculations in Figures 5C and S7B. Trial-averaged heat maps of positively-modulated cell populations were sorted by the time of maximum denoised fluorescence (Figures 5B, S5B, 7B, S7E). When comparing heat maps between two different trial types, we sorted the activity based on the left most panel (Figures S5B, S6A, 7B, S7E). All Ca^2+^ denoised fluorescence traces are z-scored with standard deviation (SD) as the unit.

#### Neural Activity Trajectory Analysis

To visualize neural activity trajectories, we used PCA for dimensionality reduction on the trial-averaged data of cells. For each cell, we computed and concatenated the trial-averaged trace of left and right trial types such that each cell’s activity was represented as a 2T × 1 dimensional vector, where T is the total number of time points per trial type. For the aggregate trajectory in Figure 6B, we combined all imaged cells into a single matrix with dimensions 2T × N where N is the total number of cells. We used PCA for dimensionality reduction and projected the data onto the first three principal components to visualize left and right neural activity trajectories. For the cell class-specific neural trajectories shown in Figure 6C, we randomly subsampled each cell class to an equal number of cells (200, a number lower than the class with the least number of imaged cells) and used PCA on the resulting matrix with dimensions 2T × (3 × 200). We projected the activity of each cell class on their respective PC loadings [for example, projection of cell class 1 = activity matrix (:,1:200) × PC loadings (1:200,:)] to visualize each cell class’ left and right neural activity trajectories independently. We randomly subsampled cells hundreds of times and the example trajectory in Figure 6C is representative of trajectories commonly observed. For Figure S6B (left) and Figure S7F, we computed the cumulative variance explained as a function of the number of PCs included, for all the data without subsampling. For Figure S6B (right) we randomly subsampled equal amounts of cells for each class. For the aggregate trajectory in Figure 7F, we performed PCA on the concatenated trial-averaged trace of correct left and right uncued trials. For the aggregate trajectory in Figure 7G, we performed PCA on the concatenated trial-averaged trace of the Approach epoch for all six trial types (L-1, L-2, R-1, R-2, L-uncued, R-uncued).

#### Regression Analysis

We used logistic regression for the analyses in Figures 6D–F, 7H, 7I, S7C, S7D, to determine how accurately left vs. right choice direction or reward context could be predicted from the neural data. For each animal, we constructed a data matrix with dimensions N × M × T where N is the number of cells, M is the number of trials and T is the number of timepoints in the z-scored fluorescence trace. For each animal, five segments of the trial were analyzed separately [−1 s to −0.05 s (Approach epoch), and 0 s to 0.45 s (early Decision epoch) using the odor onset alignment, −0.5 s to −0.05 s (late Decision epoch), 0 s to 0.7 s (Lick epoch), and 0.75 s to 2.5 s (Reward epoch) using the lick onset alignment] resulting in five data matrices per animal. To maximize the number of samples to train our classifier, we first reshaped each data matrix along the last 2 dimensions to obtain an N × (M × T) matrix. This allowed us to treat every sample across each epoch equally, meaning we would train only one classifier across all T timepoints in each epoch. We also constructed a categorical response vector for each epoch with dimensions (M × T) × 1, with left trial time points labeled as 1 and right trial time points labeled as 0.

For each data matrix, the responses of all neurons in 80% of the trials were fit with a logistic model (fitclinear in MATLAB, lasso regularization) and the remaining 20% of the trials were tested using the trained model (predict in MATLAB, 5-fold cross validation) (Figure 6D). The same process was used for each window and the predicted leftward choice accuracy was recorded. This yielded a prediction accuracy trace with 50-ms resolution for each animal and epoch. Both correct and error trials were included in the analysis for Figures 6D–F.

For the example animals in Figure 6D, we randomly subsampled the number of trials to be the same between animals (50 left trials and 50 right trials) but did not subsample the number of cells. To compare the amount of choice direction information between cell classes, we repeated the same analysis as above and randomly subsampled the number of trials and cells to be the same between all animals (50 left trials, 50 right trials, and 25 cells). We then calculated the choice direction prediction accuracy. We repeated this for 30 iterations, for each animal and for each epoch, and then plotted the mean ± SEM for each cell class (Figure 6E). To quantify the choice direction prediction accuracy as a function of the number of cells included, we repeated the same steps as above for the Decision epoch, and plotted the mean ± SEM as a function of the number of cells included for each cell class (Figure 6F).

Similar methods were used in Figures S7C and S7D for training classifiers to predict left or right choice direction, and Figures 7H and 7I for training classifiers to predict reward context in the uncued task. This required creating a new (M × T) × 1 vector of labels, with left time points labeled as 1 and right time points labeled as 0. Both correct and error trials were included for Figures S7C and S7D whereas only correct trials were included for Figures 7H and 7I because error trials in the uncued task had high behavioral variability. To compare the amount of reward context information between cell classes, we used the last 20 correct left and right trials and randomly subsampled the number of cells to be the same between all animals (25 cells). Here, two segments of the trial were analyzed separately [−1 s to 0.4 s (Approach and early Decision epochs) using the odor onset alignment, and −0.4 s to 2.5 s (late Decision and Reward epochs) using the lick onset alignment] resulting in two data matrices per animal. To quantify reward context prediction accuracy as a function of the number of cells included, we repeated the same steps as above for the Approach epoch, and plotted the mean ± SEM as a function of the number of cells included for each cell class (Figure 7I).

#### Analysis on the Effect of Reward Omission

To determine whether a cell was significantly modulated by reward omission, we used linear regression. We defined a set of four behavioral regressors that matched with the trial types: left reward, left reward omission, right reward, and right reward omission that spanned the period of reward/omission onset + 2 seconds. For each cell, we linearly regressed its deconvolved Ca^2+^ events during the entire recording period onto the set of four behavioral regressors. Cells with significant regression coefficients were then considered ‘modulated’ in that time window. To determine whether a cell had a significant regression coefficient for any of the four behavioral regressors, we used permutation tests. We randomized the regressor–We randomized theactivity relationship by shuffling the neural activity matrix with respect to the regressors, and performed linear regression to produce coefficients derived from the shuffled data. If the true coefficient was greater than the shuffled coefficient for a given cell in over 99% of iterations, it was considered significant. This allowed us to determine whether a cell was significantly modulated by reward or omission (Figure S5D). To determine whether a cell had selective activity for either reward or omission, we used the same permutation test but calculated whether the difference between the reward regressor coefficient and the omission regressor coefficient was greater than the difference found in the shuffled data. If the true difference was greater than the shuffled difference in over 99% of iterations, then the difference was considered significant. ‘Reduced activity’ cells had a significant reward regressor coefficient, which was also significantly greater than the omission regressor coefficient. ‘Elevated activity’ cells had a significant omission regressor coefficient, which was also significantly greater than the reward regressor coefficient. ‘No change’ cells had a significant reward regressor coefficient, which was not significantly different than the omission regressor coefficient (Figure S5D).

To determine whether the proportion of cells modulated by reward omission (total of ‘Reduced activity’ and ‘Elevated activity’ cells, defined above) differed between cell classes, we first calculated the true difference in the proportion of cells modulated by reward omission between classes. We then pooled all cells and randomly sampled from this pool and calculated the differences in proportions of cells modulated by reward omission between the randomized populations. If the true difference in proportions was greater (or less) than cells drawn randomly from this pool in over 99% of iterations, this was considered significant. To determine the population-level impact of reward omission, we plotted the mean response of all cells during reward and omission trials (Figure S5E).

#### Classification of Odor-, Choice-, Side-Selective Cells in the Decision Epoch

To determine whether a cell that was positively-modulated during the Decision epoch should be categorized as an Odor-, Choice-, or Side-selective cell, we used linear regression (Figure 7D). We defined a set of six behavioral regressors: left odor 1 (L-1), left odor 2 (L-2) right odor 1 (R-1), right odor 2 (R-2), left uncued (L-uncued), and right uncued (R-uncued) that spanned the decision period (odor onset to first lick). For each cell, we linearly regressed its deconvolved Ca^2+^ events during the entire recording period onto the set of six behavioral regressors. To determine whether a cell had a significant regression coefficient for any of the six behavioral regressors, we used permutation tests. We randomized the regressor–activity relationship by shuffling the neural activity matrix with respect to the regressors and performed linear regression to produce coefficients derived from the shuffled data. A cell was classified as a Side cell (L-1 and L-2 and L-uncued or R-1 and R-2 and R-uncued) if the lowest of the three left or three right coefficients was greater than the lowest of the three corresponding shuffled coefficients in over 99% of iterations. A cell was classified as a Choice cell (L-1 and L-2 or R-1 and R-2) if the lowest of the two left or two right coefficients was greater than the lowest of the two corresponding shuffled coefficients in over 99% of iterations, and that cell was not already classified as a side cell. Lastly, a cell was classified as an Odor cell (L-1 or L-2 or R-1 or R-2) if any of the coefficients was greater than the corresponding shuffled coefficients in over 99% of iterations, and that cell was not already classified as a Side or Choice cell.

To determine whether the proportion of Choice cells differed between the three cell classes, we first calculated the true difference in the proportion of choice-selective cells between the PAG-projecting, cPFC-projecting or *Rbp4Cre* classes. We then pooled all cells and randomly sampled from this pool, and calculated the differences in proportions of choice-selective cells between the randomized populations. If the true difference in proportions was greater (or less) than cells drawn randomly from this pool in over 99% of iterations, this was considered significant.

#### Classification of Mixed Selective Cells

To determine whether a cell represented information about choice direction, reward context, or both (mixed selective), we used linear regression (Figure S7G, similar to Mante et al., 2013). We defined a choice regressor that labels all time points within left trials as 1 and all time points within right trials as −1 during the 4-odor task. We similarly defined a reward context regressor during the un-cued task. For each cell, we then linearly regressed its deconvolved Ca^2+^ events onto either the choice or reward context regressors. To determine whether a cell had a significant regression coefficient for either the choice or reward context regressors, we used permutation tests. We randomized left and right trial types and performed linear regression to produce coefficients derived from the shuffled data. A cell was considered to represent information about choice direction if the true coefficient was greater (or less) than the shuffled coefficients in over 99% of iterations. This was performed similarly for reward context. Cells that had significant regressors for both choice direction and reward context were considered mixed selective cells.

To determine whether the proportion of mixed selective cells differed between cell classes, we first calculated the true difference in the proportion of mixed selective cells between classes. We then pooled all cells and randomly sampled from this pool and calculated the differences in proportions of mixed selective cells between the randomized populations. If the true difference in proportions was greater (or less) than cells drawn randomly from this pool in over 99% of iterations, this was considered significant.

#### Statistical Tests

We used MATLAB for all statistical tests unless otherwise stated. To compare differences between two groups, we used paired or unpaired t-tests, as appropriate. To compare differences between three groups, we used the one-way ANOVA or the Kruskal-Wallis test (also non-parametric) with Tukey’s honest significant difference criterion for multiple comparisons correction. In the event that standard statistical tests were not applied, we used custom permutation tests as described within each section above. All significance thresholds were set at P < 0.05 unless otherwise stated.

### Limitations

In this section we acknowledge and discuss several technical caveats raised by the reviewers that we were unable to address with new experiments, in part due to the COVID-19 pandemic.

#### Retrograde Viral Transduction

Regarding the targeting accuracy of the retrograde viruses used for the single-cell RNA-seq experiments (Figure 3), we practiced our injections extensively and were confident in our accuracy before the real experiments, and also used injection volumes and concentrations consistent with previous studies (Schwarz et al., 2015; Beier et al., 2015; Ren et al., 2018). However, because the procedure to dissect, dissociate, and FAC-sort the region of interest for sequencing was highly time-sensitive from the standpoints of cell viability and mRNA integrity, we did not prioritize saving the actual tissue from injection sites for presentation in the paper. Thus, we used atlas diagrams to illustrate our intended injection sites (Figure 3A).

Additionally, the extent of spatial spread of these retrograde viruses is not possible to quantify precisely, which is a known issue in the field. Because *CAV* and *AAV_retro_* transduce axon terminals as well as axons-in-passage (Schwarz et al., 2015; Tervo et al., 2016), both cell bodies and axons projecting to the injection site from long and short ranges are labeled. Therefore, close to the injection site, it is not possible to determine whether labeling is contributed from local circuitry or from the injection itself. Finally, we acknowledge that both *CAV* and *AAV_retro_* have tropisms in different neuronal classes from different brain regions (Schwarz et al., 2015; Tervo et al., 2016), likely due to the differential expression levels of receptors for these viruses (Li et al., 2018). In general, this problem is likely less severe in our study because it compares the same general class of neurons from the same region (L5 excitatory projection neurons in PFC). However, we cannot rule out that these factors play a role in our data.

#### Optogenetic Perturbation

Regarding the illumination zone and the extent of the optogenetics effect, we implanted our fiber tips in prelimbic cortex approximately 750 μm above the bottom of infralimbic cortex, the lower boundary of the “intended” illumination zone in vmPFC. Based on the optical settings we selected (200-μm core diameter, 0.39 NA cannulae, 5–6 mW at fiber tip), we estimated light attenuation based on the calculator at *optogenetics.org,* which indicated that the vast majority of laser light is concentrated within 500 μm of the fiber tip. This is squarely within the region of interest. Furthermore, we used laser power similar to or more conservative than previous mPFC studies in our lab and others (DeNardo et al., 2019; Huang et al., 2018; Selimbeyoglu et al., 2017; Rajasethupathy et al., 2015). Beyond this, however, we do not have empirical data demonstrating the exact extent of the optogenetics effect, but believe it is unlikely for our illuminated tissue volume to have grossly spilled beyond vmPFC.

A related point is that long-range projecting GABAergic cells are expected to exist in PFC, which could exert an inhibitory effect to projection targets outside of PFC. However, in our own sequencing data of retrogradely-labeled cells, we found that out of 1160 labeled cells from 6 regions, only 5 were GABAergic (expressed *Gad2,* = ~0.4%). This included 2 cells projecting to contralateral PFC, 2 cells projecting to hypothalamus, and 1 cell projecting to amygdala. This very low percentage in PFC is also consistent with studies from the Allen Institute that quantify other cortical areas as well (Tasic et al., 2018, also less than 1%). Furthermore, transcriptomic characterization of long-projecting GABA neurons suggest that they are likely very different from canonical inhibitory neurons, using slower neuroendocrine signaling in the place of fast neurotransmission (Paul et al., 2017). Taken together, while we acknowledge that the precise functional contribution of long-range GABAergic neurons in our assay is difficult to estimate, we believe it is unlikely to have exerted a major inhibitory effect brain-wide on the timescale of our assay.

Finally, optogenetic inhibition of projection-defined cells could test for causality in our choice behaviors. We did not pursue this avenue because: (1) Technically, the success of this experiment relies on our ability to label a large fraction of projection-defined cells with retrograde viruses for expressing inhibitory opsins, which may not be easily achievable (based on the relative abundance of PAG-*CAV*-Cre>tdTomato labeling compared with *Npr3* labeling, we estimate that our current strategy labels less than 50%); (2) Conceptually, we already demonstrated a high degree of redundancy in the encoding of task signals across cell types, and under those circumstances, it is not clear what the interpretation of such functional experiments would be. Nevertheless, we acknowledge that these experiments could be a valuable future direction.

#### Age of Mice

Regarding the usage of different ages of mice, all animals used were adults and the reason for different ages was technical. We performed all the sequencing in younger adults so that we did not have to wait for mice longer than absolutely necessary, and also because cell dissociation is relatively easier in younger animals. For the dataset in Figure 1 that we used as a reference, all animals used were very close in age. For the optogenetics and imaging mice, we needed to wait until adulthood to perform surgeries, and they were older by the time their extensive behavioral training, and optogenetics/imaging was completed. We cannot rule out the effect of age in our experiments, but consider it very unlikely to have played a major role.

### Data and Code Availability

Sequencing data will be deposited in a publicly available database such as GEO. Imaging data and analysis code is available upon request.

---

**Figure S1.**
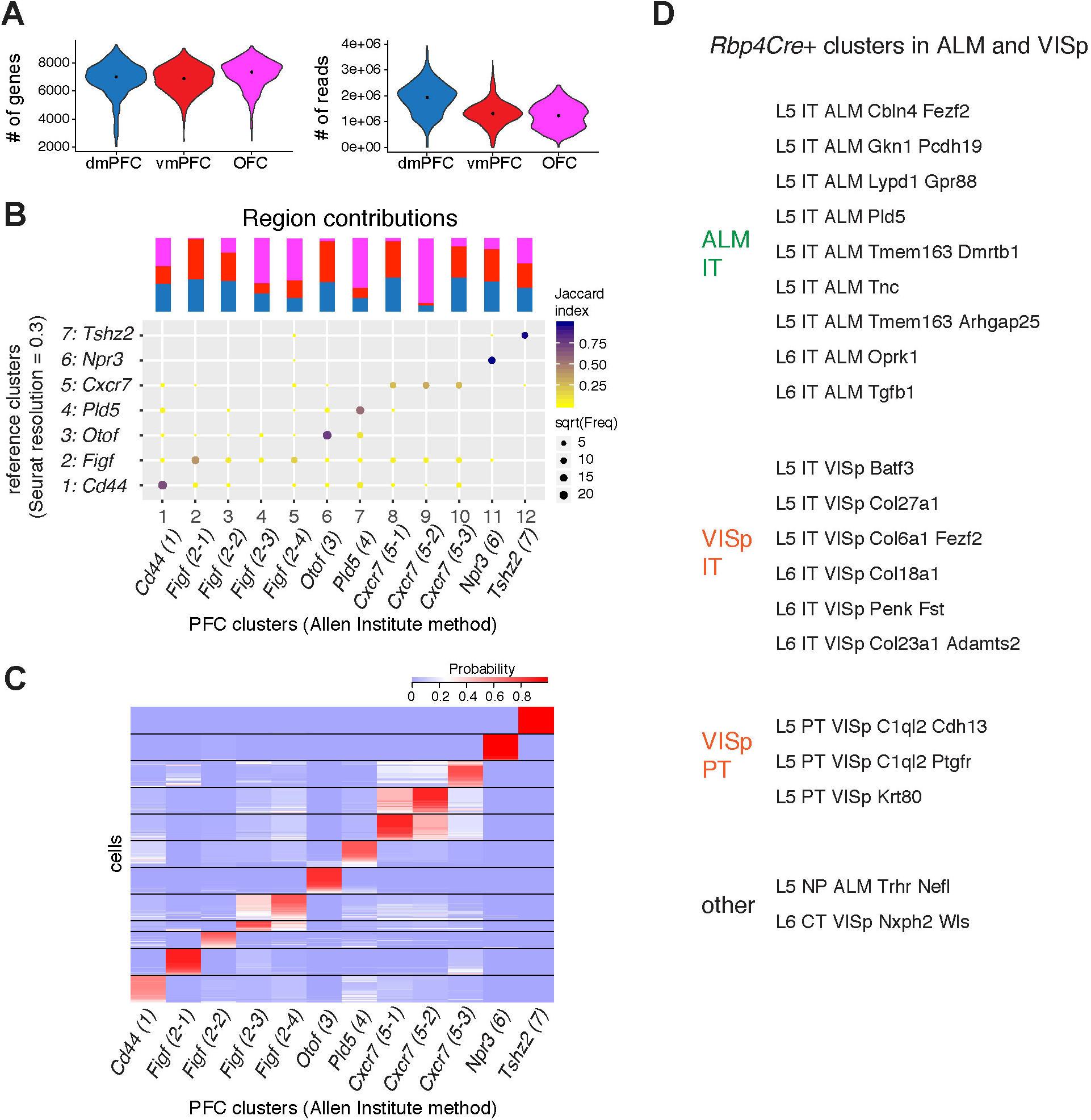
Sequencing Quality Control, Validation of PFC Neuron Clustering, and Other RNA-Sequencing Analysis, Related to Figure 1. (A) Violin plots showing the distribution of the number of genes detected (left) and the number of reads (right) across single cells derived from each of the three prefrontal cortex regions. The dots within each violin denote the median of the distribution (~7000 genes, ~1–2 million reads per cell). Effects in the gene expression data driven specifically by differences in the number of detected genes and the number of reads were regressed out from further analysis by the Seurat ScaleData function as part of the standard pipeline. (B) Comparison of *Rbp4Cre>tdTomato+* PFC neuron (n = 3139 cells, as in Figure 1) classification between reference clusters (Figure 1, Seurat resolution = 0.3) and clusters generated from the Allen Institute scrattch.hicat analysis (Tasic et al., 2018). As a quality control measure, principal components correlating with log2 detected genes, # of reads, plate format, and sample replicate were removed, see STAR Methods for additional parameters. Allen clusters are named based on similarity and subdivision from the Seurat clusters. Color scale represents the Jaccard similarity index (intersection divided by the union) of the two groups, and the size of each dot is the square root (sqrt) of the cell number. The percentage contribution of each region to each cluster (normalized to the size of the cell population) is displayed on top, with the same color code as (A). Only one cluster was highly region-specific *[Cxcr7(5-2)* with >85% contribution from OFC]. (C) Matrix depicting the robustness of *Rbp4Cre>tdTomato+* PFC neuron classification across multiple iterations of consensus clustering in scrattch.hicat. Rows represent cells, columns represent clusters, and color scale represents the probability of being assigned to the cluster based on 20 iterations of consensus clustering. (D) List of clusters from Tasic et al., 2018 that contained cells derived from *Rbp4Cre*-labeled animals, and met quality control measurements (see STAR Methods). Clusters were then grouped into the four larger categories and used for display purposes in Figure 1F.

**Figure S2.**
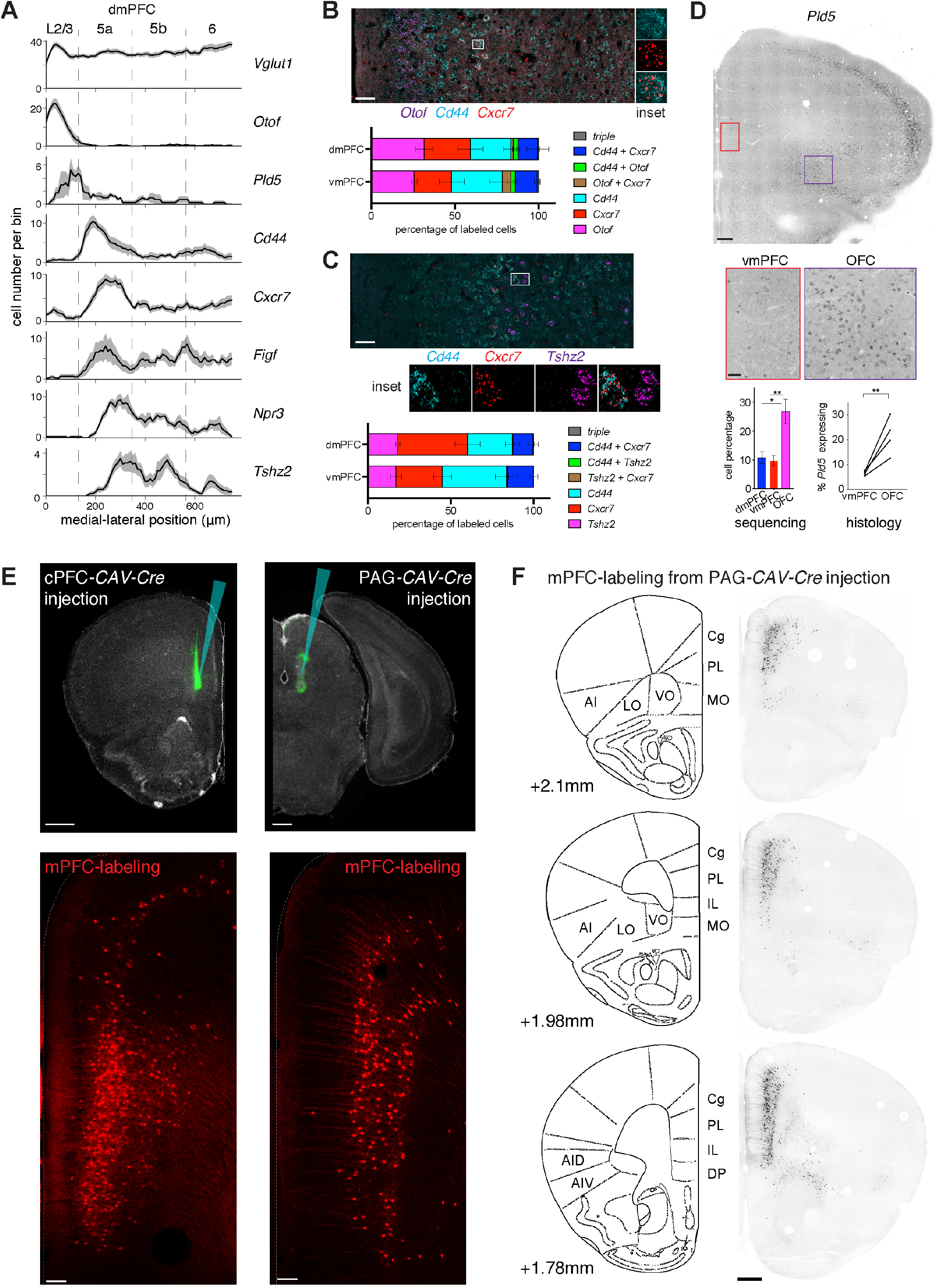
Additional HCR-FISH Validations and Example Retrograde Labeling, Related to Figures 2 and 3. (A) Laminar distribution of cells expressing cluster-specific marker genes across dmPFC, analogous to Figure 2B. Dashed lines are approximate cortical layer boundaries according to the Allen Atlas (L5a, 130 μm; L5b, 345 μm; L6, 565 μm, aligned to the beginning of L2/3). (B) Triple HCR-FISH between *Otof, Cd44,* and *Cxcr7* in vmPFC. Quantification of overlap is shown for both dmPFC and vmPFC, and averaged across n = 4 animals (543 dmPFC, 603 vmPFC cells). Error bars are SEM. Scale bar, 50 μm. (C) Triple HCR-FISH between *Cd44, Cxcr7, and Tshz2* in vmPFC. Quantification of overlap is shown for both dmPFC and vmPFC, and averaged across n = 4 animals (380 dmPFC, 372 vmPFC cells). Error bars are SEM. Scale bar, 50 μm. (D) Comparison of *Pld5* expression in mPFC vs. OFC. Sequencing data shows the percentage of cells that belonged to the *Pld5* cluster, comparing dmPFC and vmPFC with OFC (dmPFC: 910 cells, n = 3 animals; vmPFC: 1234 cells, n = 4 animals; OFC: 995 cells, n = 4 animals; *, P<0.05, **, P<0.01, unpaired t-test). Histology data shows the percentage of *Vglut1+* cells that were *Pld5+,* comparing vmPFC and OFC. Cells were quantified within a 300 × 750 μm area, and a ratio paired t-test was used, **, P<0.01. Scale bar, left: 250 μm, insets: 50 μm. (E) Injection sites of *CAV-Cre* into the contralateral PFC (−0.6/+2.2/−2.5) and PAG: (+0.4/−4.15/−2.8) of *Ai14* animals (above). Resulting retrograde labeling of tdTomato+ cells 1 week post injection at bregma A–P +1.95 mm (below). Dotted lines indicate the edge of the tissue slice. Scale bar, 500 μm (top), 100 μm (bottom). Coordinates are expressed in (M–L / A–P / D–V) axes with respect to bregma, in mm. (F) Overall distribution of PFC cells labeled by *CAV-Cre* injection into PAG: (+0.4/−4.15/−2.8) in an *Ai14* animal, highlighting presence and absence across PFC subregions. Cg, cingulate; PL, prelimbic; IL, infralimbic; MO, medial orbital; VO, ventral orbital; LO, lateral orbital; AI (D/V): agranular insular (dorsal/ventral); DP (dorsal peduncular) cortices. Scale bar, 500 μm.

**Figure S3.**
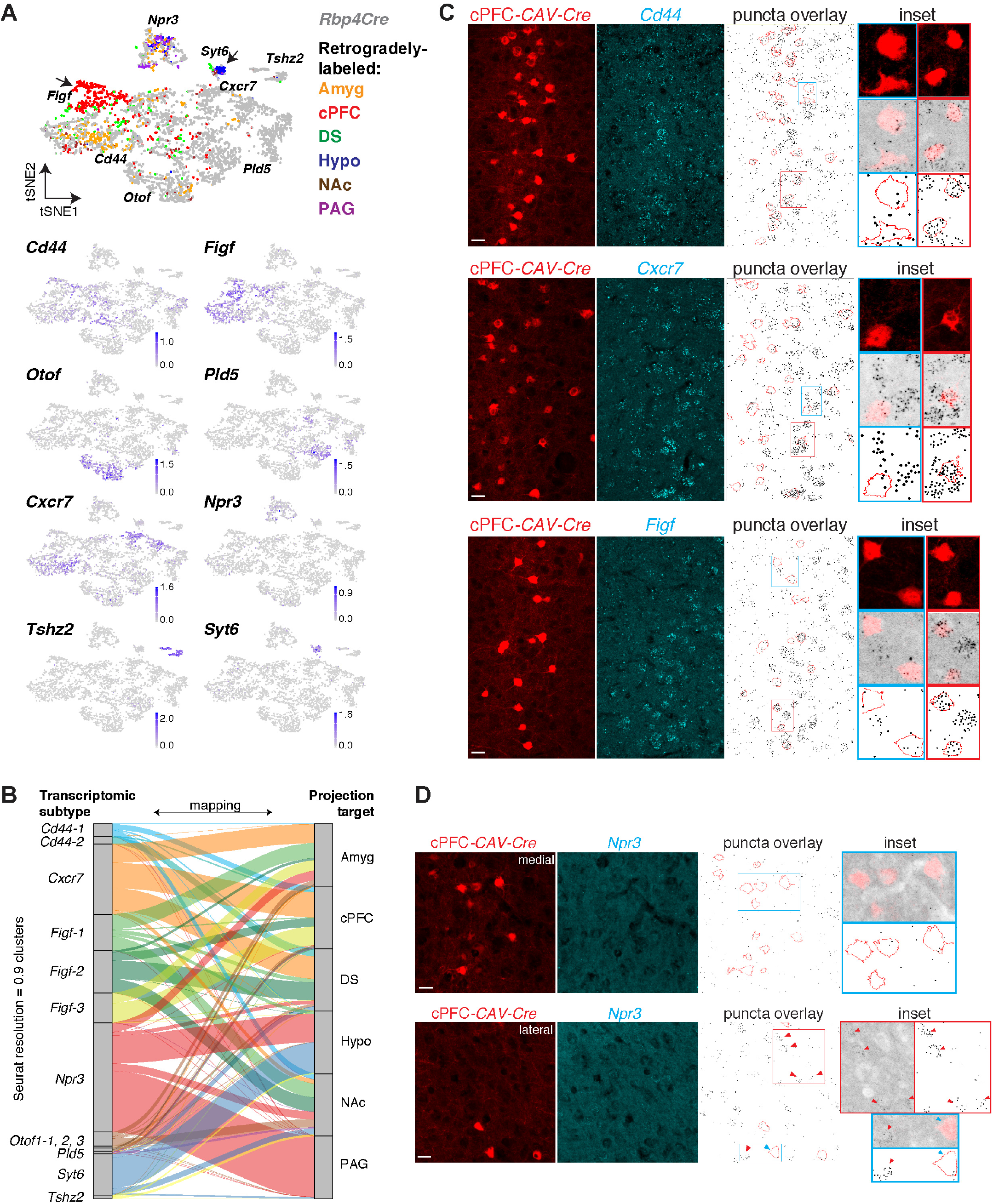
*Rbp4Cre>tdTomato+* and Projection-defined Neurons Intermingle in tSNE Space, and Additional Histological Validation of Marker Expression in Projection-defined Neurons, Related to Figure 3. (A) tSNE cluster map (top) combining all *Rbp4Cre>tdTomato+* neurons from Figure 1 (n = 3139) and retrogradely-labeled vmPFC neurons from all 6 injection sites (n = 1155), labeled by color. Areas where retrograde cells do not intermingle well with *Rbp4Cre*>tdTomato+ cells are highlighted by arrows. A *Syt6* cluster (known Layer 6 marker) is gained, and additional cells are in the *Figf* cluster. Feature plots of the seven marker genes (bottom) demonstrate that this map still roughly segregates by the marker genes identified in Figure 1, albeit with some interaction between *Figf* and *Cxcr7* clusters. Areas with relatively low retrograde cell intermingling are predominantly in *Figf* and *Syt6* clusters. Color scale represents expression level in the unit of ln[1+ (reads per 10000)]. (B) Nearest neighbor mapping of retrograde cells to transcriptomic cluster identities, at higher resolution, as an alternative to Figure 3B, represented by an alluvial diagram. Using 0.9 resolution in Seurat resulted in 12 clusters + *Syt6* as possible transcriptomic types to map retrograde cells to. Under these conditions, differences in the relative contributions of *Cd44* and *Figf* subclusters to cPFC, DS, NAc, or Amyg projections began to emerge, but no one-to-one relationships were observed. While these differences are potentially important, clustering at high resolution did not alter the general conclusions regarding the relation between transcriptomic type and projection target. Furthermore, mapping retrogradely-labeled cells to reference datasets containing only *Rbp4+* or only vmPFC cells gave similar results (data not shown). For differentially expressed marker genes of the 12 clusters based on Seurat resolution = 0.9, see Table S1 Tab 2. (C) Confocal images double-labeled with retrograde tracing from cPFC (red for the tdTomato Cre reporter) and HCR-FISH against *Cd44, Cxcr7,* or *Figf* (cyan). Scale bar, 20 μm. HCR-FISH signal was converted to binary puncta and overlaid with tdTomato cell outlines for quantification. Red insets highlight staining-positive cells (within 75% of maximum puncta observed), and cyan insets highlight staining-negative cells (<25% of maximum puncta observed), at higher magnification. (D) Confocal images double-labeled with retrograde tracing from cPFC (red for the tdTomato Cre reporter) and HCR-FISH against *Npr3* (cyan), showing enrichment of cPFC-projecting neurons in medial (superficial) and *Λpr3*-expressing cells in lateral (deeper) parts of vmPFC Layer 5. Red insets highlight *Npr3+* cells and cyan insets highlight *Npr3*- cells, at higher magnification. Red arrowheads point to *Npr3+* cells. Scale bar, 20 μm.2+

**Figure S4.**
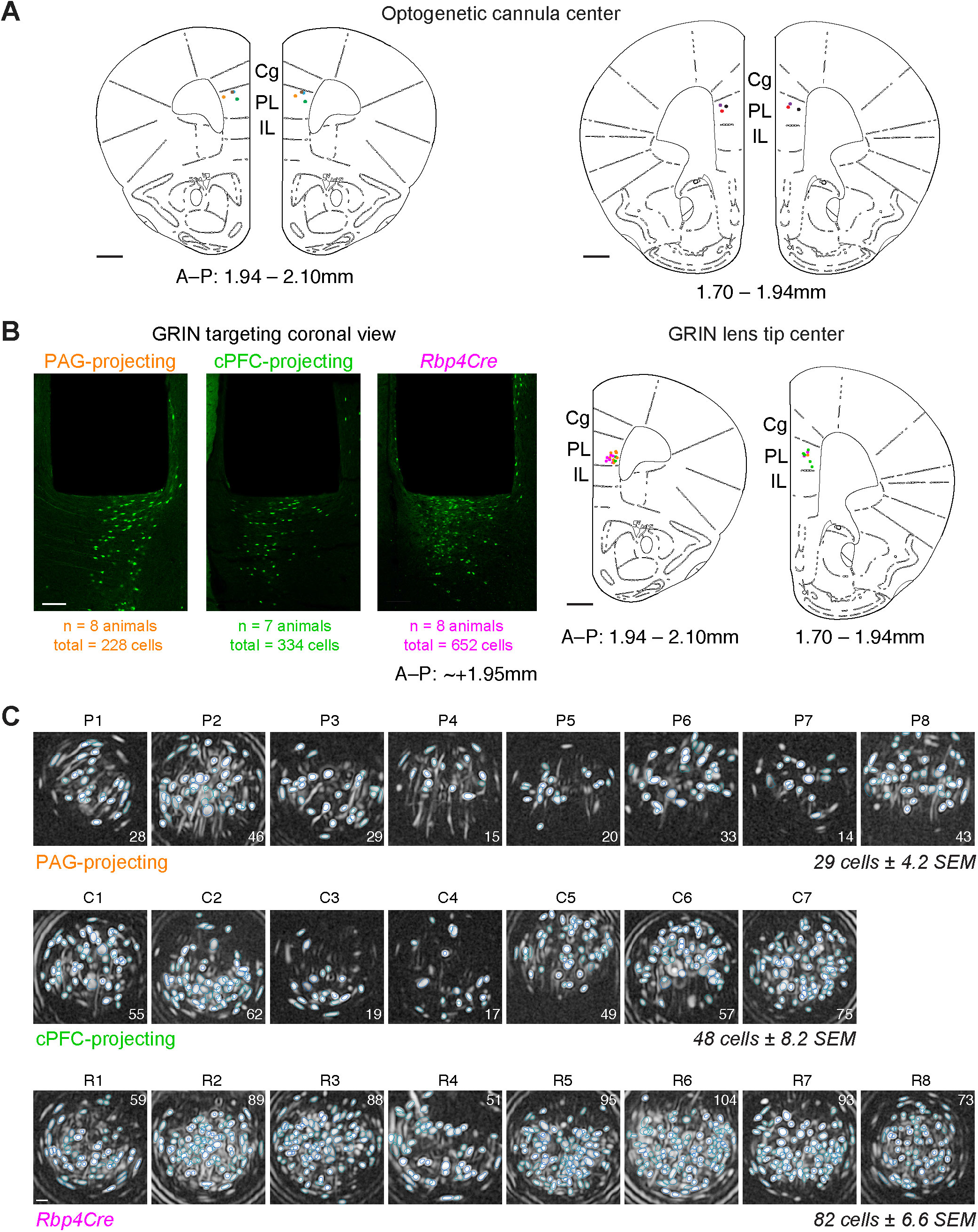
Optogenetic Cannulae Placement, GRIN Lens Targeting, and Ca^2+^ Imaging Fields of View for the 2-Odor Task, Related to Figure 4. (A) Positions of bilateral optogenetic fibers registered to a standard atlas (Paxinos and Franklin, 2001). For all animals, fibers were located within prelimbic cortex (PL; A–P: +1.95, M–L: ±0.35, D–V: −2.3). Each different colored dot denotes the location of the tip center for individual animals, and anterior-posterior (A–P) position is summarized from two adjacent atlas outlines. Scale bar, 500 μm. (B) Coronal sections of GRIN lens tracts (A–P: ~+1.95) and placement (500 μm width, 6.1 mm length, implanted at A–P: +1.95, M–L: +0.4, D–V: −2.4) in the right hemisphere registered to a standard atlas, highlighting GCaMP6f-labeled cells sitting below lens tip. Dots denote the GRIN lens tip center for each animal, and are colored by cell class (Orange: PAG-projecting, Green: cPFC-projecting, Magenta: *Rbp4Cre*). Focal plane is ~200–300 μm below the lens tip. Scale bar, 100 μm (left, images), 500 μm (right, atlas). Cg, cingulate cortex; PL, prelimbic cortex; IL, infralimbic cortex. (C) Correlation image and contour plot of neurons detected by CNMF-E in the fields of view of all imaged animals used in the 2-odor task, labeled with the number of cells analyzed and animal name (including those in Figure 4G: animals P1, C1, R2, where maximum intensity projections are displayed and rotated to be in line with the A–P axis). Colored rings are the regions of interest of individual extracted cells. Scale bar, ~25 μm.2+

**Figure S5.**
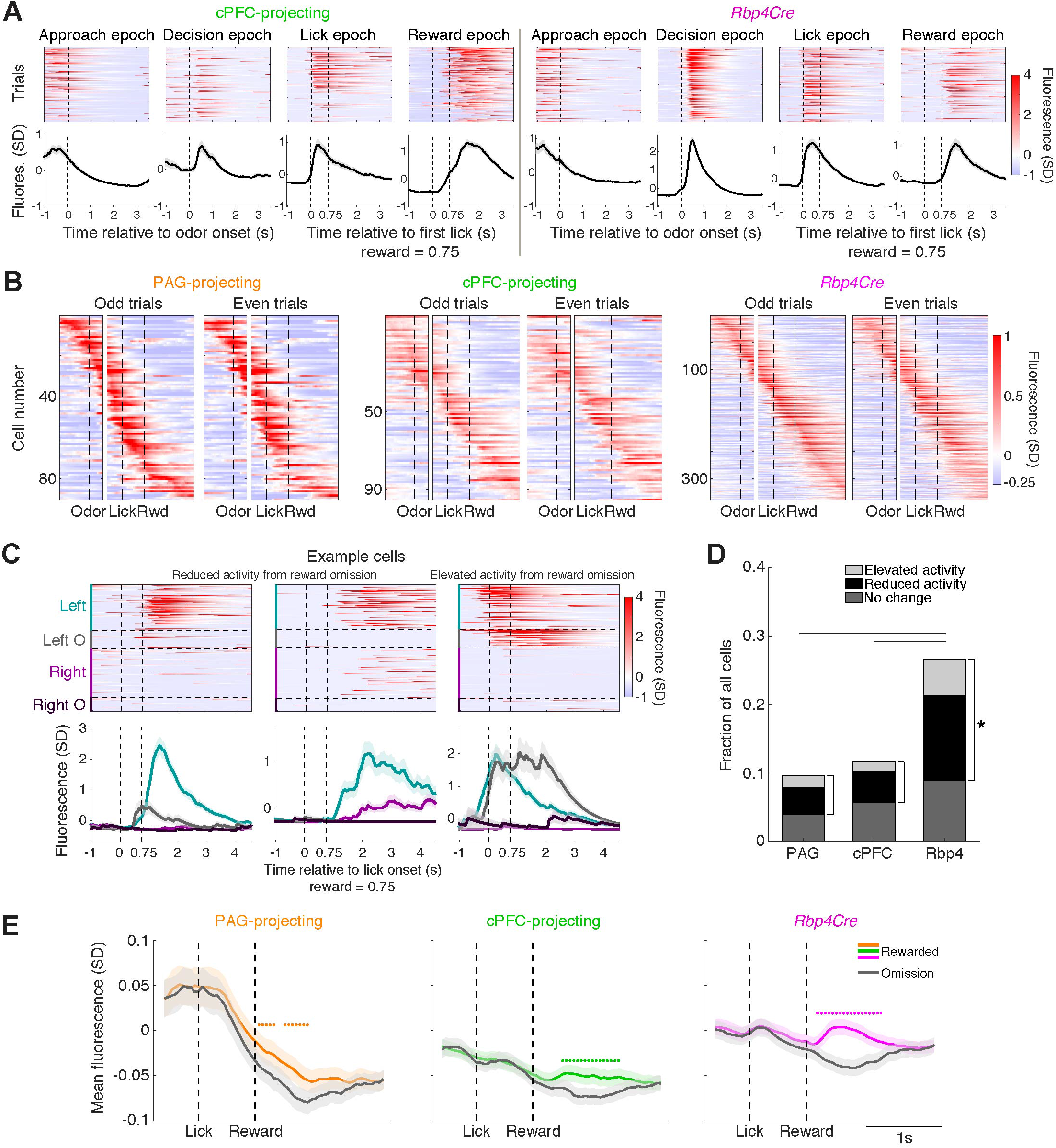
Task-Relevant Ca^2+^ Activity Dynamics Vary by Cell Class in vmPFC, and Encoding of Reward Omission, Related To Figure 5. (A) Example single-trial activity (top) and corresponding trial-averaged activity (bottom, mean ± SEM) of four task-modulated cPFC-projecting (left) and *Rbp4Cre*-labeled (right) cells during each of the four task epochs (Approach, Decision, Lick, and Reward). Traces include all correct rewarded trials. Reward was delivered 0.75 s after the first lick. Vertical dashed line in Approach/Decision epochs denotes odor onset. Vertical dashed lines in Lick/Reward epochs denote first lick and reward release (from left to right). (B) Trial-averaged activity of odd trials (left) for all positively-modulated cells sorted by the time of maximum activity, adjacent to trial-averaged activity of even trials with the same cell sorting order. Heatmap rasters are grouped by cell class, and aligned to odor onset (left) and first lick (right) (n = 90 PAG-projecting cells, n = 95 cPFC-projecting cells, n = 339 *Rbp4Cre*-labeled cells). Similar patterns of activity observed independently between odd and even trials demonstrate that the tiling pattern is not due to the sorting itself. (C) Example cells with reduced (left and middle panel) or elevated (right panel) activity during reward omission trials. Single-trial activity (top) and corresponding trial-averaged activity (bottom, mean ± SEM) is shown, color-coded by trial type. Vertical dashed lines denote lick onset and reward release. (D) Fraction of cells that: 1) had elevated activity during reward omission trials, 2) had activity during the Reward epoch that was reduced during reward omission trials, or 3) had activity during the Reward epoch that was unchanged during reward omission trials, across three cell classes. Example cells that had significantly elevated or reduced activity (Figure S5C) during omission trials were observed in all classes, indicating that information about reward omission is broadly present. Comparison of the proportion of cells modulated by reward omission (brackets: total of elevated activity and reduced activity cells) across classes was performed using a permutation test (*, P < 0.05). Reduced activity was more common than elevated activity during omission, and *Rbp4Cre*-labeled cells were most likely to be modulated. (E) Average population activity of each cell class during rewarded (colored) vs. omission trials (grey) (mean ± SEM) in the Lick and Reward epochs. The average signal within each class exhibits a net negative modulation during omission trials in the Reward epoch. Colored dots represent time points in the Reward epoch where average activity in rewarded trials is significantly higher than in omission trials (P < 0.05, paired t-test).

**Figure S6.**
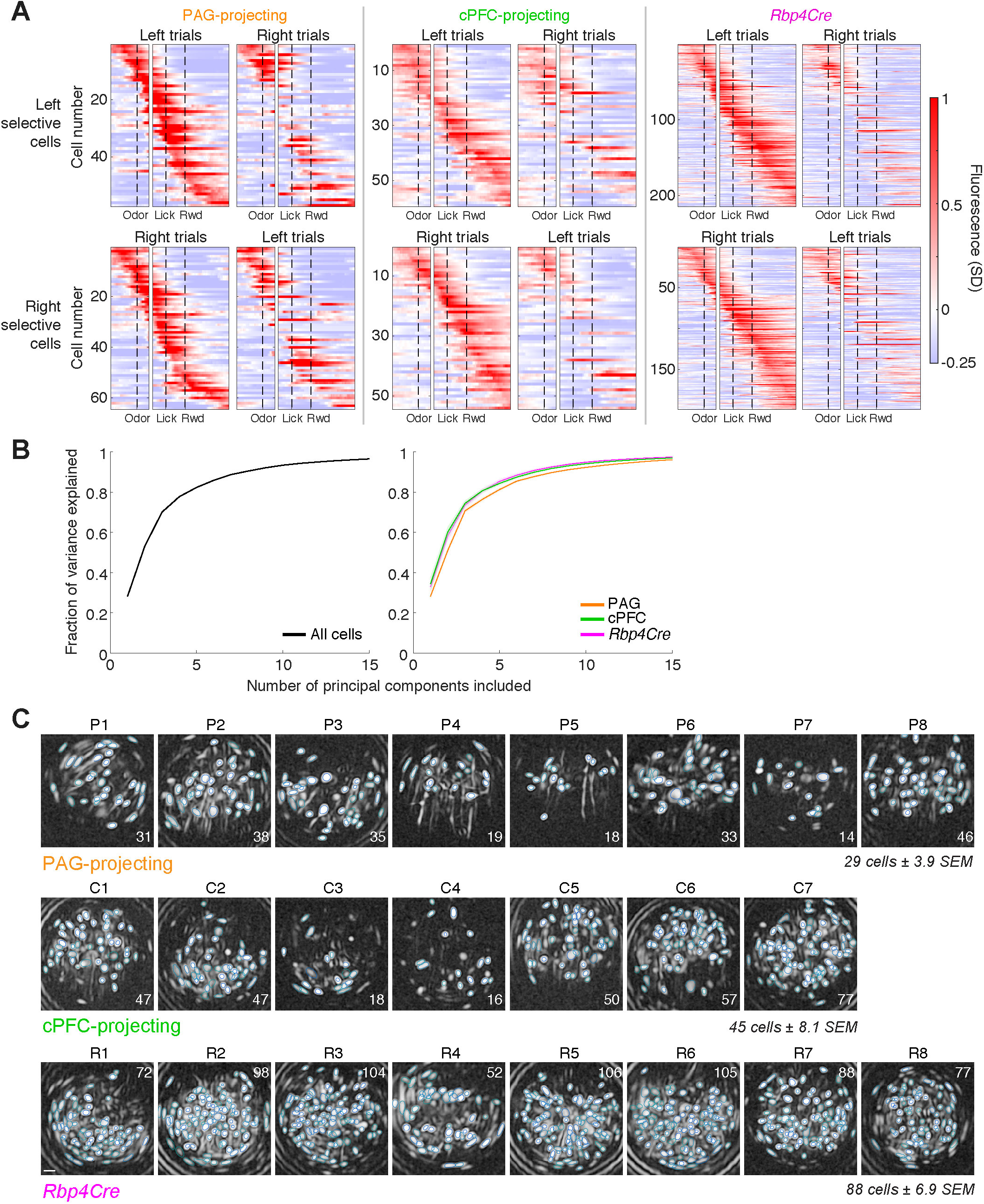
Analysis of Side-Selectivity and Dimensionality in the 2-Odor Task, and Imaging Fields of View in the 4-odor Cued and Uncued Tasks, Related to Figures 6 and 7. (A) Trial-averaged activity of left-selective cells (top) or right-selective cells (bottom). Left-selective cells are sorted by the time of peak activity on left trials and right-selective cells are sorted by that on right trials (left panels) and then compared to activity of the same cell on the opposing trial type (right panels). The loss of activity observed when comparing right panels with their corresponding left panels demonstrate the extent of side-selectivity in all three cell classes. (B) Fraction of variance explained as a function of the number of principal components included, for all cells regardless of cell class (left, related to Figure 6B) or subdivided by cell class (right, related to Figure 6C). These results indicate that the data derived from each cell class has similar dimensionality, suggesting a similar diversity of activity patterns in all classes. (C) Correlation image and contour plot of neurons detected by CNMF-E in the fields of view of all imaged animals used in the 4-odor cued and uncued tasks, labeled with the number of cells analyzed and animal name. Colored rings are the regions of interest of individual extracted cells. Scale bar, ~25 μm.

**Figure S7.**
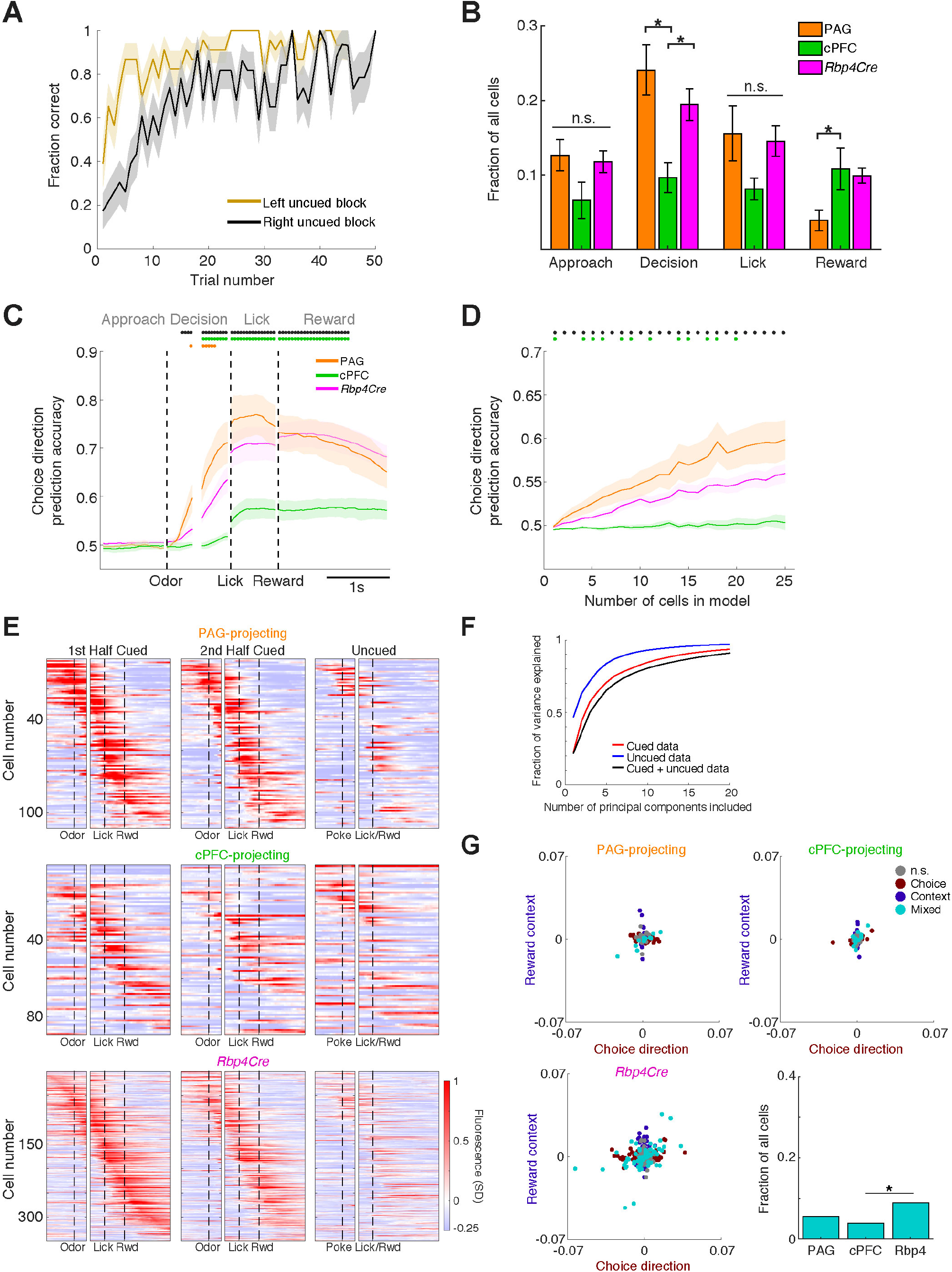
Neural Data Is Similar between the 2- and 4-Odor Tasks but Differs in the Uncued Task, and Analysis of Mixed Selectivity, Related to Figure 7. (A) Fraction of uncued trials that were correct as animals progressed through the task (mean ± SEM). (B) Cells positively-modulated in each of the four task epochs in the 4-odor task (Approach, Decision, Lick, Reward) as a fraction of all cells, on a per-animal basis (mean ± SEM; *, P < 0.05; one-way ANOVA, *post-hoc* Tukey’s HSD test; n.s., not significant). (C) Average choice direction prediction accuracy across animals in the 4-odor task (mean ± SEM, PAG-projecting n = 5, cPFC-projecting n = 5, and *Rbp4Cre* n = 8 animals). Because of the 4 trial types, data from left (L-1 + L-2) trials and right (R-1 + R-2) trials were pooled for this analysis, and randomly subsampled to 25 cells per animal (mean ± SEM). Black dots represent timepoints where the PAG-projecting trace is significantly greater than the cPFC-projecting trace. Orange dots represent timepoints where the PAG-projecting trace is significantly greater than the other two traces, and green dots represent timepoints where the cPFC-projecting trace is significantly lower than the other two traces (P < 0.05, one-way ANOVA, *post-hoc* Tukey’s HSD test). Models were computed independently for each epoch with 5-fold cross validation (randomly selected 80% of trials for training and the remaining 20% for testing). (D) The mean prediction accuracy of choice direction during the Decision epoch, as a function of the number of cells included in the logistic regression analysis (mean ± SEM) in the 4-odor task. Black dots represent timepoints where the model for PAG-projecting cells is significantly more accurate than the model for cPFC-projecting cells. Green dots represent timepoints where the model for cPFC-projecting cells is significantly less accurate than the other two (P < 0.05, one-way ANOVA, *post-hoc* Tukey’s HSD test). (E) Trial-averaged activity of all positively-modulated cells subdivided into the first half of cued trials, the second half of cued trials, and uncued trials. Heatmap rasters were sorted by the time of maximum activity and grouped by cell class, aligned to odor onset (left) and first lick (right) (n = 110 PAG-projecting cells, n = 89 cPFC-projecting cells, n = 348 *Rbp4Cre*-labeled cells). The observed similarity in activity between the 1^st^ and 2^nd^ half of the cued trials, and the dissimilarity in activity between the 2^nd^ half of the cued trials and the uncued trials, demonstrates that the difference between uncued and cued trials is not purely due to the passage of time. (F) Variance in Ca^2+^ data explained as a function of the number of principal components included, across task type (cued, uncued, and combined). The increase in dimensionality when combining the two datasets indicates that the neural activity patterns found in the two tasks are not the same. For the same imaged cells in the same field of view, novel activity patterns emerge when transitioning from the cued task to the uncued task. (G) Scatter plot of regression coefficients for choice direction in the 4-odor task vs. reward context in the uncued task for every imaged cell. We determined which cells had a significant regression coefficients) for choice direction (dark red), reward context (dark blue), or both (mixed selective, turquoise) when compared to data with randomized regressor–activity temporal relationships (see STAR Methods). Bar plot represents the proportion of mixed selective cells grouped by cell class. Comparison of the proportion of mixed selective cells across classes was performed using a permutation test (*, P < 0.05).

**Table S1. Marker Genes of the Transcriptomic Clusters in Figure 1 and Figure S3B.**

Results of differential expression analysis (Wilcoxon rank sum test, see STAR Methods) across the 7 clusters in Figure 1B (Tab 1; Seurat resolution = 0.3), or across the 12 clusters in Figure S3B (Tab 2; Seurat resolution = 0.9). This analysis takes cells within each specific cluster, compares them to all other cells not in the cluster, and then iterates through all clusters (7 or 12). In the columns of the table: ‘Gene’ is the name of the gene, ‘p_val’ is the p-value from the comparison, ‘avg_logFC’ is the average log fold change, ‘pct_cluster’ is the fraction of cells within the cluster that expresses the gene, ‘pct_other’ is the fraction of cells outside the cluster that expresses the gene, ‘p_val_adj’ is the p-value with Bonferroni correction, ‘pct_delta’ is the difference between ‘pct_cluster’ and ‘pct_other’, ‘cluster’ is the number of the cluster, and ‘cluster_name’ is the name based on the chosen exemplar marker gene used in the manuscript. To arrive at this list for Seurat resolution = 0.3 (Tab 1), we filtered differentially expressed genes by avg_logfc > 0.1, pct_cluster > 0.5, pct_other < 0.25, and pct_delta > 0.35. Together, these requirements enforce that marker genes exhibit close to binary ‘on-off’ expression in and out of the cluster. Genes actually used as markers in the paper are labeled in red. To arrive at the list for Seurat resolution = 0.9 (Tab 2), we were interested more broadly in differentially expressed genes rather than only ‘on-off’ expression. Thus, we lowered the threshold to pct_delta > 0.25, and removed the requirement for pct_other. This resulted in a longer list of 867 genes. However, we highlighted genes that fulfilled the criteria of Tab 1 (avg_logfc > 0.1, pct_cluster > 0.5, pct_other < 0.25, and pct_delta > 0.35) in red.

## REFERENCES

Allen, W.E., Chen, M.Z., Pichamoorthy, N., Tien, R.H., Pachitariu, M., Luo, L., and Deisseroth, K. (2019). Thirst regulates motivated behavior through modulation of brainwide neural population dynamics. Science 364, 253.

Allen, W.E., DeNardo, L.A., Chen, M.Z., Liu, C.D., Loh, K.M., Fenno, L.E., Ramakrishnan, C., Deisseroth, K., and Luo, L. (2017a). Thirst-associated preoptic neurons encode an aversive motivational drive. Science 357, 1149–1155.

Allen, W.E., Kauvar, I.V., Chen, M.Z., Richman, E.B., Yang, S.J., Chan, K., Gradinaru, V., Deverman, B.E., Luo, L., and Deisseroth, K. (2017b). Global Representations of Goal-Directed Behavior in Distinct Cell Types of Mouse Neocortex. Neuron 94, 891–907.e896.

Anders, S., Pyl, P.T., and Huber, W. (2015). HTSeq--a Python framework to work with high-throughput sequencing data. Bioinformatics 31, 166–169.

Asaad, W.F., Rainer, G., and Miller, E.K. (1998). Neural activity in the primate prefrontal cortex during associative learning. Neuron 21, 1399–1407.

Bari, B.A., Grossman, C.D., Lubin, E.E., Rajagopalan, A.E., Cressy, J.I., and Cohen, J.Y. (2019). Stable Representations of Decision Variables for Flexible Behavior. Neuron 103, 922–933.e927.

Beier, K.T., Steinberg, E.E., DeLoach, K.E., Xie, S., Miyamichi, K., Schwarz, L., Gao, X.J., Kremer, E.J., Malenka, R.C., and Luo, L. (2015). Circuit Architecture of VTA Dopamine Neurons Revealed by Systematic Input-Output Mapping. Cell 162, 622–634.

Betley, J.N., Cao, Z.F., Ritola, K.D., and Sternson, S.M. (2013). Parallel, redundant circuit organization for homeostatic control of feeding behavior. Cell 155, 1337–1350.

Bhattacherjee, A., Djekidel, M.N., Chen, R., Chen, W., Tuesta, L.M., and Zhang, Y. (2019). Cell type-specific transcriptional programs in mouse prefrontal cortex during adolescence and addiction. Nat Commun 10, 4169.

Bolkan, S.S., Stujenske, J.M., Parnaudeau, S., Spellman, T.J., Rauffenbart, C., Abbas, A.I., Harris, A.Z., Gordon, J.A., and Kellendonk, C. (2017). Thalamic projections sustain prefrontal activity during working memory maintenance. Nat Neurosci 20, 987–996.

Butler, A., Hoffman, P., Smibert, P., Papalexi, E., and Satija, R. (2018). Integrating single-cell transcriptomic data across different conditions, technologies, and species. Nat Biotechnol 36, 411–420.

Chen, T.W., Wardill, T.J., Sun, Y., Pulver, S.R., Renninger, S.L., Baohan, A., Schreiter, E.R., Kerr, R.A., Orger, M.B., Jayaraman, V., et al. (2013). Ultrasensitive fluorescent proteins for imaging neuronal activity. Nature 499, 295–300.

Chen, X., Sun, Y.C., Zhan, H., Kebschull, J.M., Fischer, S., Matho, K., Huang, Z.J., Gillis, J., and Zador, A.M. (2019). High-Throughput Mapping of Long-Range Neuronal Projection Using In Situ Sequencing. Cell 179, 772–786.e719.

Choi, H.M.T., Schwarzkopf, M., Fornace, M.E., Acharya, A., Artavanis, G., Stegmaier, J., Cunha, A., and Pierce, N.A. (2018). Third-generation in situ hybridization chain reaction: multiplexed, quantitative, sensitive, versatile, robust. Development 145, 12.

Churchland, M.M., Yu, B.M., Ryu, S.I., Santhanam, G., and Shenoy, K.V. (2006). Neural variability in premotor cortex provides a signature of motor preparation. J Neurosci 26, 3697–3712.

Daigle, T.L., Madisen, L., Hage, T.A., Valley, M.T., Knoblich, U., Larsen, R.S., Takeno, M.M., Huang, L., Gu, H., Larsen, R., et al. (2018). A Suite of Transgenic Driver and Reporter Mouse Lines with Enhanced Brain-Cell-Type Targeting and Functionality. Cell 174, 465–480.e422.

DeNardo, L.A., Liu, C.D., Allen, W.E., Adams, E.L., Friedmann, D., Fu, L., Guenthner, C.J., Tessier-Lavigne, M., and Luo, L. (2019). Temporal evolution of cortical ensembles promoting remote memory retrieval. Nat Neurosci 22, 460–469.

Dobin, A., Davis, C.A., Schlesinger, F., Drenkow, J., Zaleski, C., Jha, S., Batut, P., Chaisson, M., and Gingeras, T.R. (2013). STAR: ultrafast universal RNA-seq aligner. Bioinformatics 29, 15–21.

Driscoll, L.N., Pettit, N.L., Minderer, M., Chettih, S.N., and Harvey, C.D. (2017). Dynamic Reorganization of Neuronal Activity Patterns in Parietal Cortex. Cell 170, 986–999.e916.

Economo, M.N., Viswanathan, S., Tasic, B., Bas, E., Winnubst, J., Menon, V., Graybuck, L.T., Nguyen, T.N., Smith, K.A., Yao, Z., et al. (2018). Distinct descending motor cortex pathways and their roles in movement. Nature 563, 79–84.

Euston, D.R., Gruber, A.J., and McNaughton, B.L. (2012). The role of medial prefrontal cortex in memory and decision making. Neuron 76, 1057–1070.

Falk, T., Mai, D., Bensch, R., Çiçek, Ö., Abdulkadir, A., Marrakchi, Y., Böhm, A., Deubner, J., Jäckel, Z., Seiwald, K., et al. (2019). U-Net: deep learning for cell counting, detection, and morphometry. Nat Methods 16, 67–70.

Feierstein, C.E., Quirk, M.C., Uchida, N., Sosulski, D.L., and Mainen, Z.F. (2006). Representation of spatial goals in rat orbitofrontal cortex. Neuron 51, 495–507.

Friedrich, J., Zhou, P., and Paninski, L. (2017). Fast online deconvolution of calcium imaging data. PLoS Comput Biol 13, e1005423.

Fusi, S., Miller, E.K., and Rigotti, M. (2016). Why neurons mix: high dimensionality for higher cognition. Curr Opin Neurobiol 37, 66–74.

Fuster J.M. (2008). The prefrontal cortex: anatomy, physiology, and neuropsychology of the frontal lobe. 4th edn (Philadelphia: Academic Press).

Fuster, J.M., and Alexander, G.E. (1971). Neuron activity related to short-term memory. Science 173, 652–654.

Gabbott, P.L., Warner, T.A., Jays, P.R., Salway, P., and Busby, S.J. (2005). Prefrontal cortex in the rat: projections to subcortical autonomic, motor, and limbic centers. J Comp Neurol 492, 145–177.

Gerfen, C.R., Paletzki, R., and Heintz, N. (2013). GENSAT BAC cre-recombinase driver lines to study the functional organization of cerebral cortical and basal ganglia circuits. Neuron 80, 1368–1383.

Ghosh, K.K., Burns, L.D., Cocker, E.D., Nimmerjahn, A., Ziv, Y., Gamal, A.E., and Schnitzer, M.J. (2011). Miniaturized integration of a fluorescence microscope. Nat Methods 8, 871–878.

Gokce, O., Stanley, G.M., Treutlein, B., Neff, N.F., Camp, J.G., Malenka, R.C., Rothwell, P.E., Fuccillo, M.V., Südhof, T.C., and Quake, S.R. (2016). Cellular Taxonomy of the Mouse Striatum as Revealed by Single-Cell RNA-Seq. Cell Rep 16, 1126–1137.

Gong, H., Xu, D., Yuan, J., Li, X., Guo, C., Peng, J., Li, Y., Schwarz, L.A., Li, A., Hu, B., et al. (2016). High-throughput dual-colour precision imaging for brain-wide connectome with cytoarchitectonic landmarks at the cellular level. Nat Commun 7, 12142.

Greig, L.C., Woodworth, M.B., Galazo, M.J., Padmanabhan, H., and Macklis, J.D. (2013). Molecular logic of neocortical projection neuron specification, development and diversity. Nat Rev Neurosci 14, 755–769.

Guo, Z.V., Hires, S.A., Li, N., O’Connor, D.H., Komiyama, T., Ophir, E., Huber, D., Bonardi, C., Morandell, K., Gutnisky, D., et al. (2014a). Procedures for behavioral experiments in head-fixed mice. PLoS One 9, e88678.

Guo, Z.V., Li, N., Huber, D., Ophir, E., Gutnisky, D., Ting, J.T., Feng, G., and Svoboda, K. (2014b). Flow of cortical activity underlying a tactile decision in mice. Neuron 81, 179–194.

Han, Y., Kebschull, J.M., Campbell, R.A.A., Cowan, D., Imhof, F., Zador, A.M., and Mrsic-Flogel, T.D. (2018). The logic of single-cell projections from visual cortex. Nature 556, 51–56.

Harris, J.A., Mihalas, S., Hirokawa, K.E., Whitesell, J.D., Choi, H., Bernard, A., Bohn, P., Caldejon, S., Casal, L., Cho, A., et al. (2019). Hierarchical organization of cortical and thalamic connectivity. Nature 575, 195–202.

Harris, K.D., and Mrsic-Flogel, T.D. (2013). Cortical connectivity and sensory coding. Nature 503, 51–58.

Harvey, C.D., Coen, P., and Tank, D.W. (2012). Choice-specific sequences in parietal cortex during a virtual-navigation decision task. Nature 484, 62–68.

He, M., Tucciarone, J., Lee, S., Nigro, M.J., Kim, Y., Levine, J.M., Kelly, S.M., Krugikov, I., Wu, P., Chen, Y., et al. (2016). Strategies and Tools for Combinatorial Targeting of GABAergic Neurons in Mouse Cerebral Cortex. Neuron 91, 1228–1243.

Hernández, A., Nácher, V., Luna, R., Zainos, A., Lemus, L., Alvarez, M., Vázquez, Y., Camarillo, L., and Romo, R. (2010). Decoding a perceptual decision process across cortex. Neuron 66, 300–314.

Hirokawa, J., Vaughan, A., Masset, P., Ott, T., and Kepecs, A. (2019). Frontal cortex neuron types categorically encode single decision variables. Nature 576, 446–451.

Huang, W.H., Wang, D.C., Allen, W.E., Klope, M., Hu, H., Shamloo, M., and Luo, L. (2018). Early adolescent Rai1 reactivation reverses transcriptional and social interaction deficits in a mouse model of Smith-Magenis syndrome. Proc Natl Acad Sci U S A 115, 10744–10749.

Hodge, R.D., Bakken, T.E., Miller, J.A., Smith, K.A., Barkan, E.R., Graybuck, L.T., Close, J.L., Long, B., Johansen, N., Penn, O., et al. (2019). Conserved cell types with divergent features in human versus mouse cortex. Nature 573, 61–68.

Jorgenson, L.A., Newsome, W.T., Anderson, D.J., Bargmann, C.I., Brown, E.N., Deisseroth, K., Donoghue, J.P., Hudson, K.L., Ling, G.S., MacLeish, P.R., et al. (2015). The BRAIN Initiative: developing technology to catalyse neuroscience discovery. Philos Trans R Soc Lond B Biol Sci 370.

Kamigaki, T., and Dan, Y. (2017). Delay activity of specific prefrontal interneuron subtypes modulates memory-guided behavior. Nat Neurosci 20, 854–863.

Kebschull, J.M., Garcia da Silva, P., Reid, A.P., Peikon, I.D., Albeanu, D.F., and Zador, A.M. (2016). High-Throughput Mapping of Single-Neuron Projections by Sequencing of Barcoded RNA. Neuron 91, 975–987.

Kiani, R., Cueva, C.J., Reppas, J.B., and Newsome, W.T. (2014). Dynamics of neural population responses in prefrontal cortex indicate changes of mind on single trials. Curr Biol 24, 1542–1547.

Kim, C.K., Ye, L., Jennings, J.H., Pichamoorthy, N., Tang, D.D., Yoo, A.W., Ramakrishnan, C., and Deisseroth, K. (2017). Molecular and Circuit-Dynamical Identification of Top-Down Neural Mechanisms for Restraint of Reward Seeking. Cell 170, 1013–1027.e1014.

Kim, D.W., Yao, Z., Graybuck, L.T., Kim, T.K., Nguyen, T.N., Smith, K.A., Fong, O., Yi, L., Koulena, N., Pierson, N., et al. (2019). Multimodal Analysis of Cell Types in a Hypothalamic Node Controlling Social Behavior. Cell 179, 713–728.e717.

Lein, E.S., Hawrylycz, M.J., Ao, N., Ayres, M., Bensinger, A., Bernard, A., Boe, A.F., Boguski, M.S., Brockway, K.S., Byrnes, E.J., et al. (2007). Genome-wide atlas of gene expression in the adult mouse brain. Nature 445, 168–176.

Li, H., Horns, F., Wu, B., Xie, Q., Li, J., Li, T., Luginbuhl, D.J., Quake, S.R., and Luo, L. (2017). Classifying Drosophila Olfactory Projection Neuron Subtypes by Single-Cell RNA Sequencing. Cell 171, 1206–1220.e1222.

Li, N., Chen, T.W., Guo, Z.V., Gerfen, C.R., and Svoboda, K. (2015). A motor cortex circuit for motor planning and movement. Nature 519, 51–56.

Li, S.J., Vaughan, A., Sturgill, J.F., and Kepecs, A. (2018). A Viral Receptor Complementation Strategy to Overcome CAV-2 Tropism for Efficient Retrograde Targeting of Neurons. Neuron 98, 905–917.e905.

Lin, R., Wang, R., Yuan, J., Feng, Q., Zhou, Y., Zeng, S., Ren, M., Jiang, S., Ni, H., Zhou, C., et al. (2018). Cell-type-specific and projection-specific brain-wide reconstruction of single neurons. Nat Methods 15, 1033–1036.

Luo, L., Callaway, E.M., and Svoboda, K. (2018). Genetic Dissection of Neural Circuits: A Decade of Progress. Neuron 98, 865.

Madisen, L., Mao, T., Koch, H., Zhuo, J.M., Berenyi, A., Fujisawa, S., Hsu, Y.W., Garcia, A.J., Gu, X., Zanella, S., et al. (2012). A toolbox of Cre-dependent optogenetic transgenic mice for light-induced activation and silencing. Nat Neurosci 15, 793–802.

Madisen, L., Zwingman, T.A., Sunkin, S.M., Oh, S.W., Zariwala, H.A., Gu, H., Ng, L.L., Palmiter, R.D., Hawrylycz, M.J., Jones, A.R., et al. (2010). A robust and high-throughput Cre reporting and characterization system for the whole mouse brain. Nat Neurosci 13, 133–140.

Mante, V., Sussillo, D., Shenoy, K.V., and Newsome, W.T. (2013). Context-dependent computation by recurrent dynamics in prefrontal cortex. Nature 503, 78–84.

Miller, E.K., and Cohen, J.D. (2001). An integrative theory of prefrontal cortex function. Annu Rev Neurosci 24, 167–202.

Miller, E.K., Erickson, C.A., and Desimone, R. (1996). Neural mechanisms of visual working memory in prefrontal cortex of the macaque. J Neurosci 16, 5154–5167.

Moffitt, J.R., Bambah-Mukku, D., Eichhorn, S.W., Vaughn, E., Shekhar, K., Perez, J.D., Rubinstein, N.D., Hao, J., Regev, A., Dulac, C., et al. (2018). Molecular, spatial, and functional single-cell profiling of the hypothalamic preoptic region. Science 362.

Murugan, M., Jang, H.J., Park, M., Miller, E.M., Cox, J., Taliaferro, J.P., Parker, N.F., Bhave, V., Hur, H., Liang, Y., et al. (2017). Combined Social and Spatial Coding in a Descending Projection from the Prefrontal Cortex. Cell 171, 1663–1677.e1616.

Oh, S.W., Harris, J.A., Ng, L., Winslow, B., Cain, N., Mihalas, S., Wang, Q., Lau, C., Kuan, L., Henry, A.M., et al. (2014). A mesoscale connectome of the mouse brain. Nature 508, 207–214.

Otis, J.M., Namboodiri, V.M., Matan, A.M., Voets, E.S., Mohorn, E.P., Kosyk, O., McHenry, J.A., Robinson, J.E., Resendez, S.L., Rossi, M.A., et al. (2017). Prefrontal cortex output circuits guide reward seeking through divergent cue encoding. Nature 543, 103–107.

Paul, A., Crow, M., Raudales, R., He, M., Gillis, J., and Huang, Z.J. (2017). Transcriptional Architecture of Synaptic Communication Delineates GABAergic Neuron Identity. Cell 171, 522–539.e520.

Paxinos, G. and Franklin, K.B.J. (2001). The Mouse Brain in Stereotaxic Coordinates. Second Edition (Academic Press).

Pfeffer, C.K., Xue, M., He, M., Huang, Z.J., and Scanziani, M. (2013). Inhibition of inhibition in visual cortex: the logic of connections between molecularly distinct interneurons. Nat Neurosci 16, 1068–1076.

Picelli, S., Faridani, O.R., Björklund, A.K., Winberg, G., Sagasser, S., and Sandberg, R. (2014). Full-length RNA-seq from single cells using Smart-seq2. Nat Protoc 9, 171–181.

Pinto, L., and Dan, Y. (2015). Cell-Type-Specific Activity in Prefrontal Cortex during Goal-Directed Behavior. Neuron 87, 437–450.

Pinto, L., Rajan, K., DePasquale, B., Thiberge, S.Y., Tank, D.W., and Brody, C.D. (2019). Task-Dependent Changes in the Large-Scale Dynamics and Necessity of Cortical Regions. Neuron 104, 810–824.e819.

Preuss, T.M. (1995). Do rats have prefrontal cortex? The rose-woolsey-akert program reconsidered. J Cogn Neurosci 7, 1–24.

Rajasethupathy, P., Sankaran, S., Marshel, J.H., Kim, C.K., Ferenczi, E., Lee, S.Y., Berndt, A., Ramakrishnan, C., Jaffe, A., Lo, M., et al. (2015). Projections from neocortex mediate top-down control of memory retrieval. Nature 526, 653–659.

Ren, J., Friedmann, D., Xiong, J., Liu, C.D., Ferguson, B.R., Weerakkody, T., DeLoach, K.E., Ran, C., Pun, A., Sun, Y., et al. (2018). Anatomically Defined and Functionally Distinct Dorsal Raphe Serotonin Sub-systems. Cell 175, 472–487.e420.

Ren, J., Isakova, A., Friedmann, D., Zeng, J., Grutzner, S.M., Pun, A., Zhao, G.Q., Kolluru, S.S., Wang, R., Lin, R., et al. (2019). Single-cell transcriptomes and whole-brain projections of serotonin neurons in the mouse dorsal and median raphe nuclei. Elife 8.

Resendez, S.L., Jennings, J.H., Ung, R.L., Namboodiri, V.M., Zhou, Z.C., Otis, J.M., Nomura, H., McHenry, J.A., Kosyk, O., and Stuber, G.D. (2016). Visualization of cortical, subcortical and deep brain neural circuit dynamics during naturalistic mammalian behavior with head-mounted microscopes and chronically implanted lenses. Nat Protoc 11, 566–597.

Rigotti, M., Barak, O., Warden, M.R., Wang, X.J., Daw, N.D., Miller, E.K., and Fusi, S. (2013). The importance of mixed selectivity in complex cognitive tasks. Nature 497, 585–590.

Rose, J.E., and Woolsey, C.N. (1948). The orbitofrontal cortex and its connections with the mediodorsal nucleus in rabbit, sheep and cat. Res Publ Assoc Res Nerv Ment Dis 27 (1 vol.), 210–232.

Rushworth, M.F., Noonan, M.P., Boorman, E.D., Walton, M.E., and Behrens, T.E. (2011). Frontal cortex and reward-guided learning and decision-making. Neuron 70, 1054–1069.

Saunders, A., Macosko, E.Z., Wysoker, A., Goldman, M., Krienen, F.M., de Rivera, H., Bien, E., Baum, M., Bortolin, L., Wang, S., et al. (2018). Molecular Diversity and Specializations among the Cells of the Adult Mouse Brain. Cell 174, 1015–1030.e1016.

Schmitt, L.I., Wimmer, R.D., Nakajima, M., Happ, M., Mofakham, S., and Halassa, M.M. (2017). Thalamic amplification of cortical connectivity sustains attentional control. Nature 545, 219–223.

Schwarz, L.A., Miyamichi, K., Gao, X.J., Beier, K.T., Weissbourd, B., DeLoach, K.E., Ren, J., Ibanes, S., Malenka, R.C., Kremer, E.J., et al. (2015). Viral-genetic tracing of the input-output organization of a central noradrenaline circuit. Nature 524, 88–92.

Selimbeyoglu, A., Kim, C.K., Inoue, M., Lee, S.Y., Hong, A.S.O., Kauvar, I., Ramakrishnan, C., Fenno, L.E., Davidson, T.J., Wright, M., et al. (2017). Modulation of prefrontal cortex excitation/inhibition balance rescues social behavior in. Sci Transl Med 9.

Shekhar, K., Lapan, S.W., Whitney, I.E., Tran, N.M., Macosko, E.Z., Kowalczyk, M., Adiconis, X., Levin, J.Z., Nemesh, J., Goldman, M., et al. (2016). Comprehensive Classification of Retinal Bipolar Neurons by Single-Cell Transcriptomics. Cell 166, 1308–1323.e1330.

Shenoy, K.V., Sahani, M., and Churchland, M.M. (2013). Cortical control of arm movements: a dynamical systems perspective. Annu Rev Neurosci 36, 337–359.

Siciliano, C.A., Noamany, H., Chang, C.J., Brown, A.R., Chen, X., Leible, D., Lee, J.J., Wang, J., Vernon, A.N., Vander Weele, C.M., et al. (2019). A cortical-brainstem circuit predicts and governs compulsive alcohol drinking. Science 366, 1008–1012.

Soudais, C., Laplace-Builhe, C., Kissa, K., and Kremer, E.J. (2001). Preferential transduction of neurons by canine adenovirus vectors and their efficient retrograde transport in vivo. FASEB J 15, 2283–2285.

Spellman, T., Rigotti, M., Ahmari, S.E., Fusi, S., Gogos, J.A., and Gordon, J.A. (2015). Hippocampal-prefrontal input supports spatial encoding in working memory. Nature 522, 309–314.

Stamatakis, A.M., Schachter, M.J., Gulati, S., Zitelli, K.T., Malanowski, S., Tajik, A., Fritz, C., Trulson, M., and Otte, S.L. (2018). Simultaneous Optogenetics and Cellular Resolution Calcium Imaging During Active Behavior Using a Miniaturized Microscope. Front Neurosci 12, 496.

Stanley, G., Gokce, O., Malenka, R.C., Südhof, T.C., and Quake, S.R. (2020). Continuous and Discrete Neuron Types of the Adult Murine Striatum. Neuron 105, 688–699.e688.

Steinmetz, N.A., Zatka-Haas, P., Carandini, M., and Harris, K.D. (2019). Distributed coding of choice, action and engagement across the mouse brain. Nature 576, 266–273.

Stuart, T., Butler, A., Hoffman, P., Hafemeister, C., Papalexi, E., Mauck, W.M., Hao, Y., Stoeckius, M., Smibert, P., and Satija, R. (2019). Comprehensive Integration of Single-Cell Data. Cell 177, 1888–1902.e1821.

Taniguchi, H., He, M., Wu, P., Kim, S., Paik, R., Sugino, K., Kvitsiani, D., Kvitsani, D., Fu, Y., Lu, J., et al. (2011). A resource of Cre driver lines for genetic targeting of GABAergic neurons in cerebral cortex. Neuron 71, 995–1013.

Tasic, B., Yao, Z., Graybuck, L.T., Smith, K.A., Nguyen, T.N., Bertagnolli, D., Goldy, J., Garren, E., Economo, M.N., Viswanathan, S., et al. (2018). Shared and distinct transcriptomic cell types across neocortical areas. Nature 563, 72–78.

Tervo, D.G., Hwang, B.Y., Viswanathan, S., Gaj, T., Lavzin, M., Ritola, K.D., Lindo, S., Michael, S., Kuleshova, E., Ojala, D., et al. (2016). A Designer AAV Variant Permits Efficient Retrograde Access to Projection Neurons. Neuron 92, 372–382.

Tinevez, J.Y., Perry, N., Schindelin, J., Hoopes, G.M., Reynolds, G.D., Laplantine, E., Bednarek, S.Y., Shorte, S.L., and Eliceiri, K.W. (2017). TrackMate: An open and extensible platform for single-particle tracking. Methods 115, 80–90.

Tosches, M.A., Yamawaki, T.M., Naumann, R.K., Jacobi, A.A., Tushev, G., and Laurent, G. (2018). Evolution of pallium, hippocampus, and cortical cell types revealed by single-cell transcriptomics in reptiles. Science 360, 881–888.

Uchida, N., and Mainen, Z.F. (2003). Speed and accuracy of olfactory discrimination in the rat. Nat Neurosci 6, 1224–1229.

Uylings, H.B., Groenewegen, H.J., and Kolb, B. (2003). Do rats have a prefrontal cortex? Behav Brain Res 146, 3–17.

van der Maaten, L.J.P, Hinton, G.E. (2008). Visualizing high dimensional data using t-SNE. Journal of Machine Learning Research. 9(Nov):2579–2605.

Vander Weele, C.M., Siciliano, C.A., Matthews, G.A., Namburi, P., Izadmehr, E.M., Espinel, I.C., Nieh, E.H., Schut, E.H.S., Padilla-Coreano, N., Burgos-Robles, A., et al. (2018). Dopamine enhances signal-to-noise ratio in cortical-brainstem encoding of aversive stimuli. Nature 563, 397–401.

Wagner, M.J., Kim, T.H., Kadmon, J., Nguyen, N.D., Ganguli, S., Schnitzer, M.J., and Luo, L. (2019). Shared Cortex-Cerebellum Dynamics in the Execution and Learning of a Motor Task. Cell 177, 669–682.e624.

Winnubst, J., Bas, E., Ferreira, T.A., Wu, Z., Economo, M.N., Edson, P., Arthur, B.J., Bruns, C., Rokicki, K., Schauder, D., et al. (2019). Reconstruction of 1,000 Projection Neurons Reveals New Cell Types and Organization of Long-Range Connectivity in the Mouse Brain. Cell 179, 268–281.e213.

Yuste, R., Hawrylycz, M., Aalling, N., Aguilar-Valles, A., Arendt, D., Arnedillo, R.A., Ascoli, G., Bielza, C., Bokharie, V., Bergmann, T.B., et al. (2020). A community-based transcriptomics classification and nomenclature of neocortical cell types. Nat Neurosci in press.

Zappia, L, Oshlack, A. (2018). Clustering trees: a visualization for evaluating clusterings at multiple resolutions. GigaScience. 7. DOI:gigascience/giy083

Zeisel, A., Hochgerner, H., Lönnerberg, P., Johnsson, A., Memic, F., van der Zwan, J., Häring, M., Braun, E., Borm, L.E., La Manno, G., et al. (2018). Molecular Architecture of the Mouse Nervous System. Cell 174, 999–1014.e1022.

Zeng, H., and Sanes, J.R. (2017). Neuronal cell-type classification: challenges, opportunities and the path forward. Nat Rev Neurosci 18, 530–546.

Zhou, P., Resendez, S.L., Rodriguez-Romaguera, J., Jimenez, J.C., Neufeld, S.Q., Giovannucci, A., Friedrich, J., Pnevmatikakis, E.A., Stuber, G.D., Hen, R., et al. (2018). Efficient and accurate extraction of in vivo calcium signals from microendoscopic video data. Elife 7, e28728.

Zingg, B., Hintiryan, H., Gou, L., Song, M.Y., Bay, M., Bienkowski, M.S., Foster, N.N., Yamashita, S., Bowman, I., Toga, A.W., et al. (2014). Neural networks of the mouse neocortex. Cell 156, 1096–1111.

